# Molecular phylogenetics and micromorphology of Australasian Stipeae (Poaceae), and the interrelation of whole-genome duplication and evolutionary radiations in this grass tribe

**DOI:** 10.1101/2020.06.05.129320

**Authors:** Natalia Tkach, Marcin Nobis, Julia Schneider, Hannes Becher, Grit Winterfeld, Mary E. Barkworth, Surrey W. L. Jacobs, Martin Röser

**Affiliations:** Martin Luther University Halle-Wittenberg, Institute of Biology, Geobotany and Botanical Garden, Dept. of Systematic Botany, Neuwerk 21, 06108 Halle, Germany; Institute of Botany, Faculty of Biology, Jagiellonian University, Gronostajowa 3 st., 30-387 Kraków, Poland; Institute of Evolutionary Biology, School of Biological Sciences, University of Edinburgh, Charlotte Auerbach Road, Edinburgh EH9 3FL, UK; Biology Department, Utah State University, 5305 Old Main Hill, Logan Utah, 84322, USA

**Author notes:** Addresses for correspondence: Martin Röser,; Natalia Tkach. deceased (Surrey W. L. Jacobs).

**Keywords:** *Achnatherum*, *Anemanthele*, *Austrostipa*, chromosome number, dysploidy, long-distance dispersal, micromorphology, phylogeny, Poaceae, polyploidy, SEM, Stipeae, taxonomy, whole-genome duplication

## Abstract

The mainly Australian grass genus *Austrostipa* with ca. 64 species represents a remarkable example of an evolutionary radiation. To investigate aspects of diversification, macro- and micromorphological variation in this genus we conducted a molecular phylogenetic and scanning electron microscopy (SEM) analysis including representatives from all of its accepted subgenera.

Plastid DNA variation within *Austrostipa* was low and only few lineages were resolved. Nuclear ITS and *Acc1* yielded comparable groupings of taxa and resolved subgenera *Arbuscula*, *Petaurista*, *Bambusina* in a common clade and as monophyletic. In summary, we suggest recognizing nine subgenera in *Austrostipa*.

Because of its taxonomic significance in Stipeae, we studied the lemma epidermal structure in 34 representatives of *Austrostipa.* In most species, the lemma epidermal pattern (LEP) was relatively uniform (maize-like LEP), but in six species it was more similar to that of *Stipa* s.str., *Neotrinia*, *Ptilagrostis* and *Orthoraphium.* The species representing subgenera *Lobatae*, *Petaurista*, *Bambusina* and *A. muelleri* from subg. *Tuberculatae* were well-separated from all the other species included in the analysis.

Two different sequence copies of *Acc1* were found in polyploid *Austrostipa* and *Anemanthele*. Each of the copy types formed a single clade. This was also true of the sampled species of *Stipa* s.str., but their clades were strongly separated from those of *Austrostipa* and *Anemanthele*. This underlines the statement of Tzvelev (1977) that most if not all contemporary Stipeae are of hybrid origin and demonstrates it for the first time unambiguously on the molecular level.

Chromosome number variation is surveyed and reviewed for the whole tribe Stipeae and interpreted in a molecular phylogenetic framework. The rather coherent picture of chromosome number variation underlines the phylogenetic and evolutionary significance of this character.

The closest extant relatives of *Austrostipa* and *Anemanthele* are in the clade of *Achnatherum* s.str., *Celtica*, *Oloptum* and *Stipellula*. These genera are most abundant in Central and Eastern Asia, which makes a colonization of Australian and New Zealand from this region more likely, perhaps via long-distance dispersal, than colonization of Australia from southern South America via Antarctica as previously invoked.

**Supporting Information** may be found online in the Supporting Information section at the end of the article.

## INTRODUCTION

The Australasian grass genus *Austrostipa* S.W.L.Jacobs & J.Everett encompasses ca. 64 species, most of which occur in Australia and Tasmania (Vickery & al., 1986; Jacobs & Everett, 1996; Everett & al., 2009; Williams, 2011). Only one species, *A. stipoides* (Hook.f.) S.W.L.Jacobs & J.Everett, is considered native to New Zealand; it is also present in southeastern Australia. Other species of Australian origin are naturalized in New Zealand (Jacobs & al., 1989; Edgar & Connor, 2000). One species, *A. scabra* (Lindl.) S.W.L.Jacobs & J.Everett, is present on Easter Island but it is thought to have been introduced there after 1860 (Everett & Jacobs, 1990).

*Austrostipa* belongs to the tribe Stipeae, a group of ca. 600 species, with almost worldwide distribution with major centers of radiation in Eurasia (especially in *Stipa* L. s.str., *Piptatherum* P.Beauv.) and the Americas (in *Nassella* (Trin.) É.Desv., *Eriocoma* Nutt., *Piptochaetium* J.Presl, *Pappostipa* (Speg.) Romasch., P.M.Peterson & Soreng). *Austrostipa* and the endemic New Zealand genus *Anemanthele* Veldkamp (one species) are together with New Zealand *Achnatherum petriei* (Buchanan) S.W.L.Jacobs & J.Everett the only indigenous representatives of this tribe in the Australian Plant Kingdom.

The grasses of tribe Stipeae have long attracted the interest of scientists not only because of their enormous morphological variation but also because of their worldwide ecological significance as important constituents of grasslands (steppes, prairies) under mesic to xeric, sometimes in rather cold climates where they are often important source of food for the livestock of humans. The delineation of this tribe, its taxonomic structure with regards to major lineages and the circumscription of its genera has been investigated during the past three decades using morphological, anatomical and, more recently, molecular phylogenetic approaches (Freitag, 1975, 1985; Tzvelev, 1976; Barkworth, 1983, 1990, 1993, 2007; Vickery & al., 1986; Barkworth & Everett, 1987; Jacobs & Everett, 1996; Cialdella & al., 2007, 2010, 2014; Edgar & Connor, 2000; Jacobs & al., 2000, 2007; Barkworth & Torres, 2001; Peñailillo, 2002, 2003; Soreng & al., 2003; Wu & Phillips, 2006; Barkworth & al., 2008; Romaschenko & al., 2008, 2010, 2011, 2012, 2013, 2014; Barber & al., 2009; Everett & al., 2009; Schneider & al., 2009, 2011; Vásquez Pardo & al., 2011; Hamasha & al., 2012; Sclovich & al., 2015; Krawczyk & al., 2017, 2018; Nobis & al., 2019a,b, 2020; Peterson & al., 2019).

Ecologically, *Austrostipa* is adapted to the warm-to hot-summer Mediterranean climate of SW, S and SE Australia and the more Oceanic climate of SE Australia and Tasmania. This rather isolated southern outpost of Stipeae is widely separated from temperate Eurasia and distant to the Americas, both regions with high diversity in Stipeae genera. It is notable for its large number of species, 64, which is indicative of a fast evolutionary radiation. The species also display a tremendous morphological diversity in, for example, habit, growth form, the size and form of individual structures such as spikelets, glumes, etc. Moreover, the rich evolutionary diversification occurred sympatric primarily in a comparatively narrow coastal strip of southwestern, southern and southeastern Australia, the seemingly climatically suitable part of this continent for plants preferring Mediterranean-type to steppe climate with open vegetation as typical of Stipeae. Many of the species seem to be edaphically specialized, being restricted to specific soil types (Everett & al., 2009; Williams, 2011).

Although different types of lemma indumentum, visible even under low magnification, are best known and frequently used in identification of many species within Poaceae, the taxonomic value of micromorphological characters of the lemma epidermis is also substantial in many genera of grasses (for example, Thomasson, 1978, 1981, 1986; Terrell & Wergin, 1981; Barkworth & Everett, 1987; Valdés-Reyna & Hatch, 1991; Snow, 1996; Acedo & Llamas, 2001; Terrell & al., 2001; Mejía Saulés & Bisby, 2003; Ortúñez & de la Fuente, 2010; Nobis, 2013). Thomasson (1978, 1981) was the first to use several epidermal ultrastructural characters of the lemma, such as the presence of hooks, the shape of the long cells and the presence of silica cells, for identification of fossil and extant representatives of Stipeae and documented the phylogenetic importance of epidermal pattern in this tribe. Building on his work, various lemma epidermal patterns have been distinguished in Stipeae, with two, *Stipa*-like (also called saw-like) and *Achnatherum*-like (maize-like), being the most common (Barkworth & Everett, 1987; Romaschenko & al., 2012; Nobis & al., 2019b,c, 2020). Despite, however, the taxonomic significance of lemma epidermal patterns (LEP) in the Stipeae being well established, only the pattern of *Austrostipa scabra* has been described so far (Romaschenko & al., 2012). It confirmed its affiliation with the achnatheroid species. Bustam (2010) examined LEPs in 33 *Austrostipa* species but did not report on the patterns observed in the individual species, focusing instead on the species clusters she identified. Several authors pointed out that, even if LEP is relatively uniform within a genus, it may still be useful in identifying particular species as well as in delineating relationships among and between different subgenera or sections (Ortúñez & de la Fuente, 2010; Nobis, 2013; Olonova & al., 2016; Nobis & al., 2019b).

Many of the more recent studies converged in recognizing *Austrostipa* as separate from, and only distantly related to, *Stipa* s.str. in Eurasia and North Africa (Barkworth & al., 2008; Romaschenko & al., 2012; Hamasha & al., 2012), thus supporting its status as separate genus proposed by Jacobs & Everett (1996). Morphologically, *Austrostipa* has several floret characteristics that, although not individually unique to the genus, in combination distinguish it from other genera, including the rather similar and poorly understood genus *Achnatherum*. These characters are its long, sharp calluses and often dark and tough the lemma margins, the glabrous and prow-tipped palea (Jacobs & Everett, 1996). The closest extant relatives of *Austrostipa* within the Stipeae, however, have not yet been unequivocally identified. Analyses of morphological and anatomical data placed *Austrostipa* in a clade together with *Achnatherum* P.Beauv. and *Ptilagrostis* Griseb. (Jacobs & Everett, 1996; Jacobs & al., 2000: Fig. 2). Previous molecular phylogenetic studies showed *Austrostipa* forming together with the main part of *Achnatherum*, the American genera *Nassella*, *Jarava*, and several smaller genera (for example, *Amelichloa* Arriaga & Barkworth, *Celtica* F.M.Vázquez & Barkworth, *Stipellula* Röser & Hamasha) one of the well supported major lineages within the tribe (Barkworth & al., 2008; Romaschenko & al., 2008, 2010, 2012; Cialdella & al., 2010; Hamasha & al., 2012). Most studies sampled only one or a few species of *Austrostipa*, making it hard to assess the monophyly of this genus. One species of *Austrostipa* was sampled for the internal transcribed spacer (ITS) regions of nrDNA by Hsiao & al. (1999), 13 for ITS and five plastid DNA regions by Romaschenko & al. (2008, 2010, 2012), six for ITS1 and seven for four plastid DNA regions by Barkworth & al. (2008), two for four plastid DNA regions by Cialdella & al. (2010) and five for ITS and two for one plastid DNA region by Hamasha & al. (2012). The ITS studies of Jacobs & al. (2000, 2007) encompassed 15 and 37 species, respectively. While monophyly of *Austrostipa* was supported by the former study, sequences of some species of *Achnatherum*, *Nassella*, and *Stipa* were interspersed in the *Austrostipa* clade of the latter. In both studies, the New Zealand endemic *Anemanthele* was included in the *Austrostipa* clade, but its position was unstable. The most comprehensive study conducted so far included 31 taxa for ITS and 52 for two plastid DNA regions (Syme & al., 2012).

Overall variation between individual *Austrostipa* ITS sequences was low and the differences between sequences from different accessions of the same species was often not much smaller than between sequences of different species (Jacobs & al., 2007). This overall low variation made it also difficult to compare their results with classification of *Austrostipa* into 13 subgenera (Jacobs & Everett, 1996; Everett & al., 2009). Some of the subgenera were reflected in the ITS data (for example, subg. *Falcatae* S.W.L.Jacobs & J.Everett), whereas others were mixed up (for example, subg. *Austrostipa* and subg. *Tuberculatae* S.W.L.Jacobs & J.Everett or subg. *Arbuscula* S.W.L.Jacobs & J.Everett and subg. *Bambusina* S.W.L.Jacobs & J.Everett, respectively), or were entirely unresolved (Jacobs & al., 2007; Syme & al., 2012). The plastid DNA analyses resolved two main clades, none of which corresponded to the recognized subgenera, and further resolution was low (Syme & al., 2012).

By using diversified morpho-molecular approaches, this study intends to address some of the main phylogenetic and evolutionary problems connected with *Austrostipa* and not yet fully resolved, namely its monophyly and its internal phylogenetic structure. These questions are addressed on a broader sample of *Austrostipa* taxa than in previous studies by generating a taxonomically overlapping set of nr ITS and plastid DNA sequences of the 3’*trnK* region. Both molecular markers are frequently utilized and well-established in molecular phylogenetic studies (Baldwin & al., 1995; Liang & Hilu, 1996), although ITS from the repetitive 18S–26S nrDNA can be polymorphic in individual genomes for several reasons., This leads to paralogous sequence relationships that can potentially confound phylogenetic reconstruction (Buckler & al., 1997; Álvarez & Wendel, 2003; Bailey & al., 2003; Razafimandimbison & al., 2004; Bayly & Ladiges, 2007; Nieto Feliner & Rosselló, 2007; Schneider & al., 2009, 2011). The 3’*trnK* region, comprising the 3’part of the chloroplast *matK* gene with following intron and 3’*trnK* exon, was selected as sequence marker from the plastid DNA mainly because of its comparatively high substitution rate. Moreover, these sequences are straightforward to align and are already available in many potential outgroup taxa from within Stipeae and neighboring tribes (Döring & al., 2007; Schneider & al., 2009, 2011, 2012; Hamasha & al., 2012; Blaner & al., 2014; Wölk & Röser, 2014, 2017; Hochbach & al., 2015, 2018; Tkach & al., 2020). For DNA extraction, we make use of plant material, which was collected and preserved in the field in saturated NaCl/CTAB buffer solution especially for DNA work (see Material and Methods).

A further objective of this study was to explore the value of the nuclear single-copy gene *Acc1* encoding plastid acetyl-CoA carboxylase 1 (Huang & al., 2002; Fan & al., 2007, 2009; Sha & al., 2010; Hochbach & al., 2015) for molecular phylogenetic work in Stipeae. This gene cannot be successfully amplified from DNA isolated from normal herbarium specimens, even if such plant material is suitable for amplification of ITS or plastid DNA sequences. The presence of different copies of gene *Acc1* in single specimens of *Austrostipa* soon became evident, thus we were forced to clone these sequences via bacterial plasmids. Additionally, we conducted cytogenetic work in *Austrostipa*, most of which was already published by Winterfeld & al. (2015), to find out if our specimens were diploids or polyploids. We use the resulting increased knowledge on chromosomes of Stipeae to update and complement a previous survey of chromosome numbers (see Romaschenko & al., 2012), to discuss the basic chromosome number(s), dysploid variation and the evolutionary role of whole-genome duplications in Stipeae.

## MATERIAL AND METHODS

### Plant material

The sample for the molecular phylogenetic study included 51 species and subspecies of *Austrostipa*. Geographic origin, collector or seed exchange locality and herbarium vouchers for the taxa used in this study are listed in Appendix 1. For half of the species more than one specimen were included. Sampling density among the 13 subgenera for the analysed DNA regions 3’*trnK* (3’part of the chloroplast *matK* gene with the following intron and 3’*trnK* exon), ITS and *Acc*1 is summarized in Table 1. The dataset of the 3’*trnK* region encompassed 43, that of ITS 48 *Austrostipa* species, representing 71% and 75% of the total species number, respectively. All subgenera except subg. *Lanterna* S.W.L.Jacobs & J.Everett were represented by at least half their species. For the analysis of the nuclear single-copy gene *Acc*1 sequence data we selected at least one specimen of each subgenus and studied a total of 22 (33%) species. In addition, we studied representatives of 8 stipoid genera (*Achnatherum*, *Anemanthele*, *Celtica*, *Nassella*, *Neotrinia* (Tzvelev) M.Nobis, P.D.Gudkova & A.Nowak, *Oloptum* Röser & Hamasha, *Stipa*, *Stipellula*) that previous studies have shown to be the most closely related to *Austrostipa* (Jacobs & al., 2000, 2007; Barkworth & al., 2008; Romaschenko & al., 2008, 2010, 2012; Hamasha & al., 2012). Genera from the tribes Bromeae (*Bromus* L.), Duthieeae (*Anisopogon*) and Triticeae (*Henrardia* C.E.Hubb.*, Hordeum* L., *Secale* L.) were chosen as outgroups for phylogenetic reconstructions based on ITS and 3’*trnK* regions based on recent studies in phylogenetic relationships within subf. Pooideae (for example, Catalán & al., 1997; Hilu & al., 1999; Mathews & al., 2000; Soreng & Davis, 2000; GPWG, 2001; Davis & Soreng, 2007; Döring & al., 2007; Soreng & al., 2007; Schneider & al., 2009, 2011; Saarela & al., 2015, 2018). For the *Acc*1 dataset exemplary species from Bromeae (*Bromus inermis* Leyss.) and Triticeae (*Henrardia persica* (Boiss.) C.E.Hubb., *Hordeum chilense* Roem. & Schult.*, H. vulgare* L.) were used as outgroup taxa taken from ENA/GenBank.

**Table 1.**
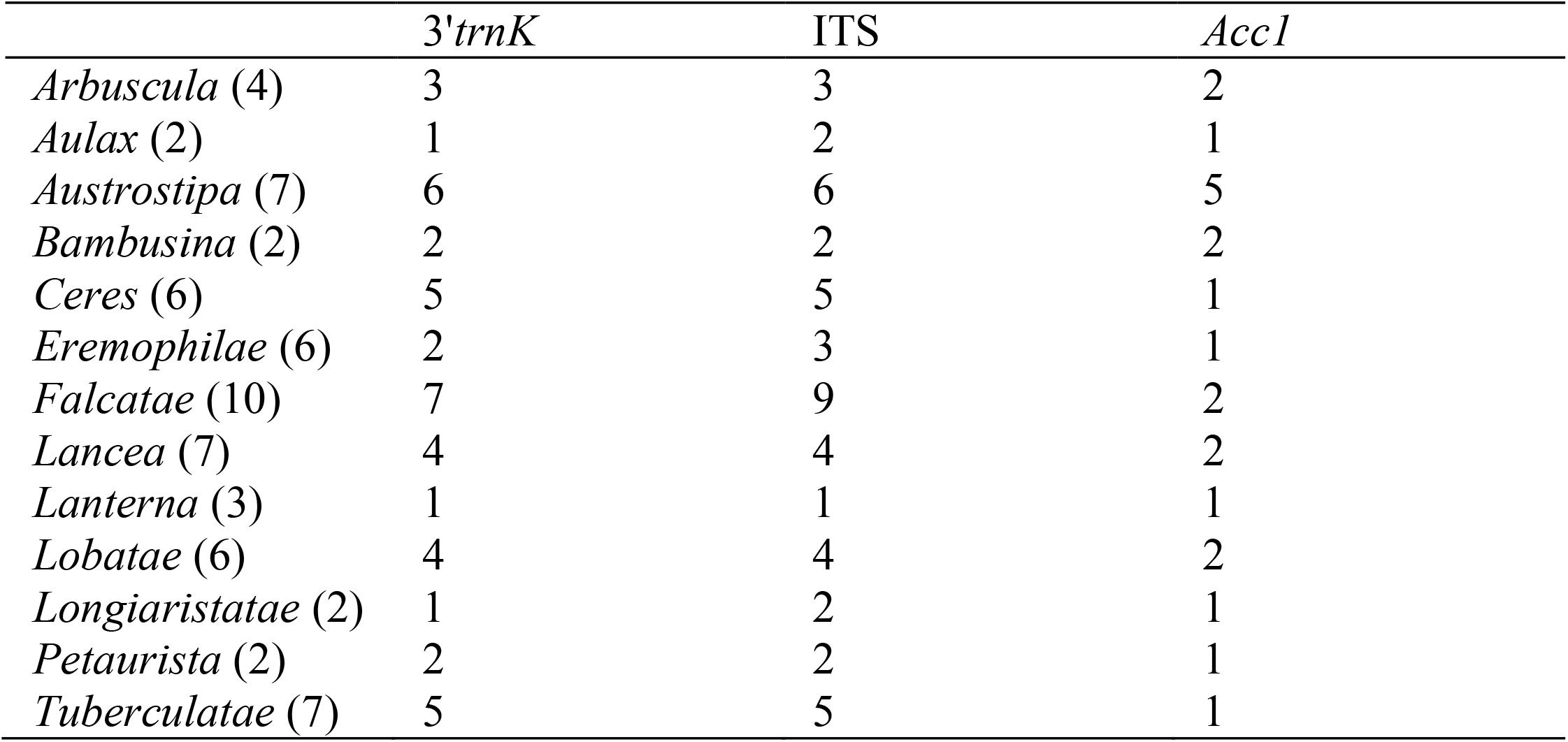
Overview of sampling density among the 13 previous subgenera of *Austrostipa* used for molecular phylogenetic analyses. The number of species sampled relative to the total number of species (in brackets) according to Jacobs & Everett (1996), Everett & al. (2009) and Williams (2011) is shown for the three DNA regions analyzed.

Most plant material used in this study for DNA extraction was collected in the field in 2007 by SWLJ and MEB along with herbarium specimens and duplicates, which have been subsequently distributed to various herbaria (Appendix 1). The leaf samples were preserved in saturated NaCl/CTAB buffer solution prepared according to Štorchová & al. (2000). Further leaf material for DNA extraction were collected from living pot plants grown from seeds stored at the Millennium Seed Bank (Wakehurst Place, Royal Botanic Gardens, Kew, UK). These caryopses were collected from natural populations with verified identifications and voucher specimens deposited at herbaria K and partly PERTH (Appendix 1). The pot plants were cultivated in the greenhouses of the Botanical Garden of the University Halle-Wittenberg (vouchers at HAL). Leaves for DNA extractions were silica gel-dried (Chase & Hills, 1991) and the same plants were used for cytogenetic studies (Winterfeld & al., 2015).

### DNA extraction, PCR amplification and sequencing

For DNA extraction leaves preserved in NaCl/CTAB buffer were removed from the solution, rinsed in water, immersed in liquid nitrogen, and then ground to fine powder using mortar and pestle. Silica gel-dried fresh leaves were shredded in a FastPrep FP 120 bead mill homogenizer (Qbiogene, Heidelberg, Germany). The ready-to-use NucleoSpin Plant Kit (Macherey-Nagel, Düren, Germany) was used for extraction.

The ITS and 3’*trnK* region were amplified and sequenced according our previous studies with primers listed in Table 2 (Schneider & al., 2009, 2011, 2012; Hamasha & al., 2012; Winterfeld & al., 2015). The amplification of *Acc*1 (exons 6–13 and intervening introns) was carried out using primers also listed in Table 2. An overview of the gene *Acc1* is shown in Fig. 1 together with the locations, directions and designations of the primers used in this study.

**Fig. 1.**
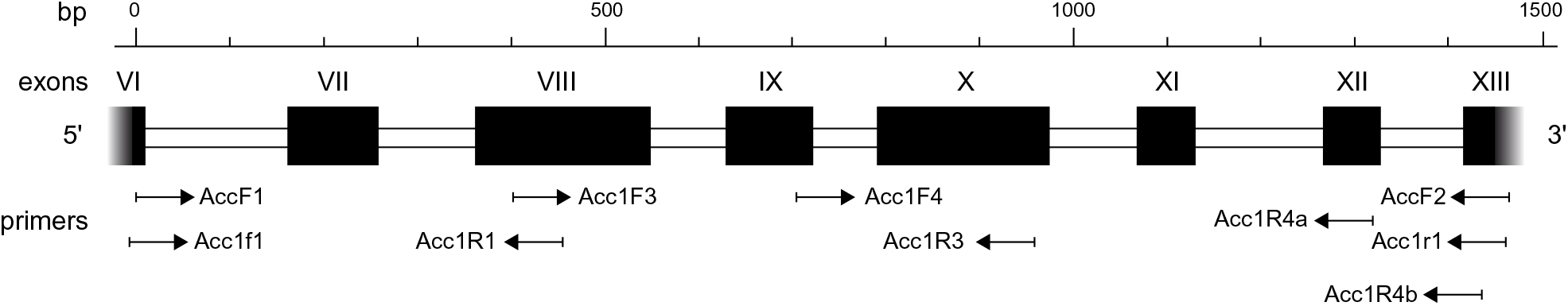
Schematic of the acetyl-CoA carboxylase gene (*Acc1*) modified from Huang & al. (2002) with designations, locations and directions of the primers used.

**Table 2.**
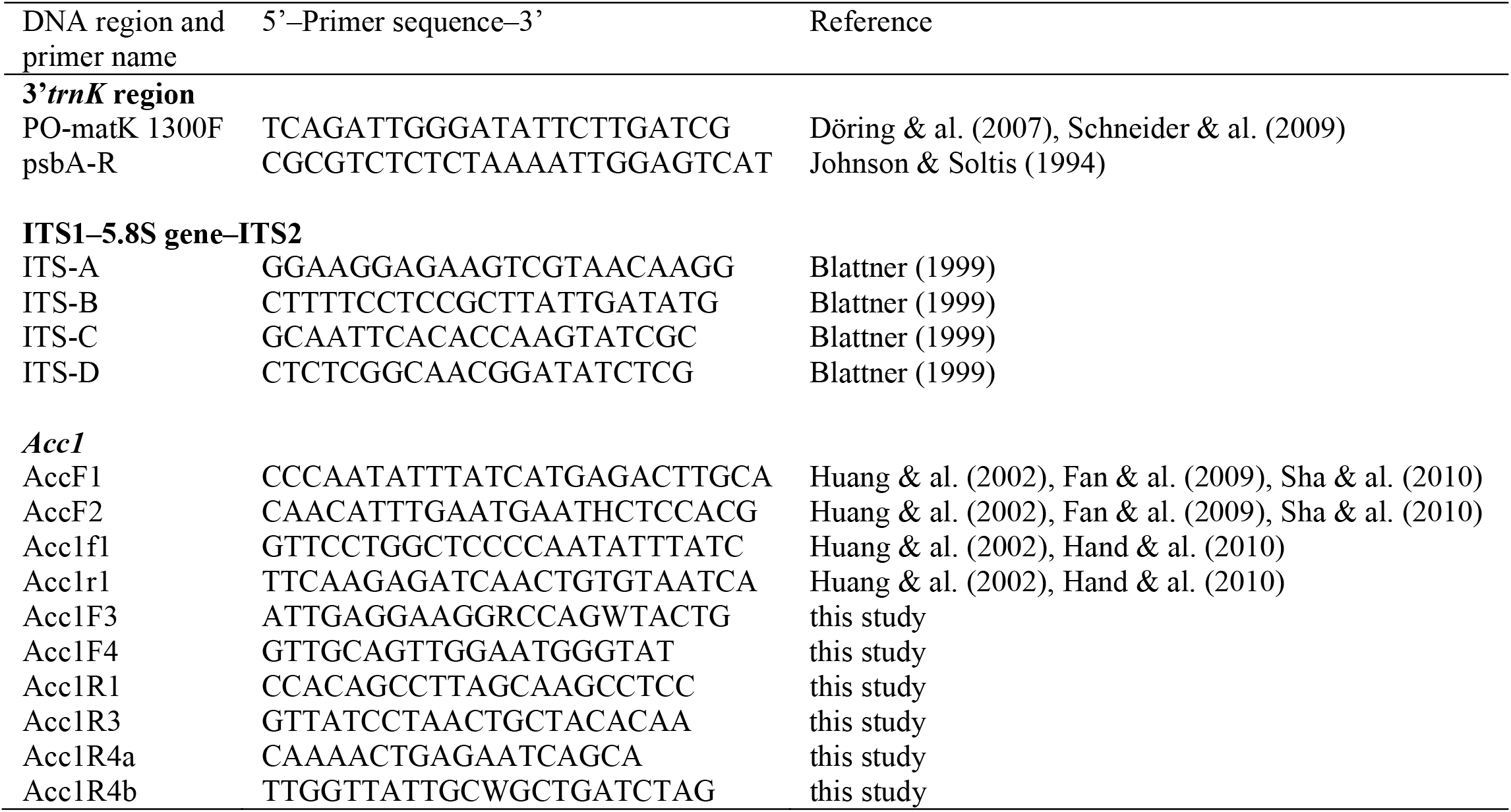
Primers used to amplify and sequence the plastid 3’*trnK* region, nuclear ITS1–5.8S gene–ITS2 and the *Acc1* gene (exons 6‒13 and intervening introns). For *Acc1* primer positions see Fig. 1.

The PCR reactions of 20 μl usually contained 0.5 μM of each primer, 2 μl of 10 × PCR buffer, 1.9 mM MgCl_2_, 0.8–1 U *Taq* DNA polymerase (all MP Biomedicals, Heidelberg, Germany), 5% DMSO (AppliChem, Darmstadt, Germany), 100 μM dNTPs (GeneCraft, Lüdinghausen, Germany), 1–2 μl of template DNA (∼ 50 ng) and distilled water. For DNA samples, which were obtained by extraction from leaves preserved in saturated NaCl/CTAB buffer solution, the PCR reaction was performed with 3 min at 94°C, followed by 35 cycles of 30 s at 94°C, 30 s–2 min at 50°C, 5 min at 68°C, and a final extension for 20 min at 68°C. The DNA extracted from silica gel-dried leaf material was amplified by the following PCR program: 3 min at 94°C, followed by 35 cycles of 30 s at 94°C, 30 s–2 min at 50°C, 2 min at 72°C, and a final extension for 20 min at 72°C. PCR products of *Acc*1 were column-purified with the NucleoSpin Extract ΙΙ Kit (Macherey-Nagel).

The purified PCR-products of the analysed diploids species (*Achnatherum paradoxum* (L.) Banfi, Galasso & Bartolucci ≡ *Piptatherum paradoxum* (L.) P.Beauv., *A. sibiricum* (L.) Keng ex Tzvelev) and *Nassella trichotoma* (Nees) Hack. ex Arechav.), were sequenced directly by StarSEQ GmbH (Mainz, Germany) or Eurofins MWG Operon (Ebersberg, Germany). Because species of *Austrostipa* and *Anemanthele lessoniana* are tetra- or hexaploid (Appendix 2; Winterfeld & al., 2015), *Acc*1 amplicons were cloned into the pGEM-T Easy Vector (Promega, Mannheim, Germany) according to the manufacturer’s protocol. In the next step 10–30 individual white colonies containing the insert were picked. The isolation of plasmid DNA was performed with the Wizard Plus SV Minipreps DNA Purification System (Promega). The insert of the purified plasmid DNA was sequenced using the standard primers T7 and SP6.

### Alignment and phylogenetic analysis

All sequences were edited by eye in Sequencher v.5.0 (Gene Codes, Ann Arbor, MI, USA). The automatically performed alignments by using ClustalW2 (Larkin & al., 2007) were manually adjusted in Geneious v.9.1.6 (https://www.geneious.com; Kearse & al., 2012). We identified few double peaks in chromatograms of the ITS dataset already documented in our previous study (Winterfeld & al., 2015). It was possible to edit these single nucleotide positions by IUPAC code and include all obtained ITS sequences.

All clone sequences of the *Acc1* dataset were visually checked in terms of the presence of chimerical sequences or PCR artefacts (see Brassac & al., 2012). Furthermore, we tested the protein sequence of the exon regions (696 bp) for each clone and compared the translation to the *Acc1* sequence of the diploid outgroup, *Bromus inermis*, taken from ENA/GenBank (Appendix 1). We excluded chimerical sequences and clone sequences different from that of *Bromus inermis* in more than 20 amino acid positions of the exon regions. To reduce the number of singletons in the alignment, we summarized for each specimen highly similar *Acc1* sequences of the remaining individual clones to consensus sequences.

Sequences of ITS and 3’*trnK* region as well as the individual *Acc*1 clone sequences used for assembling consensus sequences were submitted to ENA/GenBank under the accession numbers ####-#### (Appendix 1).

All DNA sequence datasets were analysed using the phylogenetic approaches of maximum likelihood (ML), maximum parsimony (MP) and Bayesian inference (BI) following Tkach & al. (2019). The trees were visualized with FigTree v.1.4.3 (http://tree.bio.ed.ac.uk/software/figtree/). Support values are cited in the text in the following sequence: ML bootstrap support/MP bootstrap support/Bayesian posterior probability (PP).

### Morphological analyses

We estimated or scored nine morphological characters in each of 65 taxa (species and subspecies) of *Austrostipa* examined: mean length of ligules of the culm leaves; surface of inflorescence branches (glabrous, with prickles or macrohairs); lengths of glumes, calli, lemmas, lemma lobes and awns; length ratio lemma:palea; shape of palea apex (without or with 2‒4 teeth) (Table 3, suppl. Tables S1, S2). These characters were chosen to elucidate phylogenetic groupings of *Austrostipa* taxa in reference to the intergeneric division proposed by Jacobs & Everett (1996) and Everett & al. (2009). For some species not available for morphological study the descriptions of Everett & al. (2009) were used (see Appendix 1).

**Table 3.** Morphological characters and character states.

|  | Character states |
| --- | --- |
| **Macromorphological characters** |  |
| Ligules | mean length [mm] |
| Glumes | mean length [mm] |
| Callus | mean length [mm] |
| Lemma | mean length [mm] |
| Lemma lobes | mean length [mm] |
| Awn | mean length [mm] |
| Length lemma:palea | ratio |
| Inflorescence branches | glabrous or scabrous (0); pilose (1) |
| Palea apex | 2–4-toothed (0); without teeth (1) |
| **Micromorphological characters of the lemma epidermis** |  |
| Basal cells | wider than long to as long as wide (1); up to 1.8 times longer than wide (2); 2–3 times longer than wide (3) |
| Hooks | present (1); absent (2) |
| Silica cells | elongated, 2–3 times longer than wide (1); ovate to elliptic, up to 1.5 times longer than wide (2) |
| Cork cells | present (1); absent (2) |

### Lemma micromorphology

The ultrastructure of the lemma epidermis was studied in 34 taxa (species and subspecies) of *Austrostipa* (Appendix 1). For scanning electron microscopy (SEM) observation, dry samples were coated with a thin layer of gold using a JFC-1100E ion sputter (JEOL), then observed and photographed on a Hitachi S-4700 scanning electron microscope. Six diagnostic micromorphological characters, namely basal cells, silica bodies, cork cells, hooks, prickles and macrohairs were examined in the middle part of the abaxial lemma surface and photographs were taken.

The numerical analyses were performed on the same 34-taxa set based on (1) four micromorphological characters, and (2) a combination of eight macromorphological and four micromorphological characters (Table 3, suppl. Tables S1, S2). The complete data matrix is presented in suppl. Table S1. Each taxon was treated as an Operational Taxonomic Unit (OTU), in accordance with the methods used in numerical taxonomy (Sokal & Sneath, 1963). The similarities among OTUs were calculated using Gower’s general similarity coefficient. Cluster analysis, using PAST software (Hammer & al., 2001), was performed on all OTUs to estimate morphological similarities among the species.

## RESULTS

### Molecular phylogenetics

We analysed a dataset of 110 DNA sequences for the 3’*trnK* region and 111 for ITS, respectively. The *Acc1* dataset comprised a total of 266 clone sequences. After evaluation of all clone sequences of the polyploid genera *Austrostipa* and *Anemanthele*, as described in Material and Methods, we created 61 consensus sequences for the final dataset. We obtained two or three distinct *Acc1* consensus sequences for each *Austrostipa* species with exception of *A. breviglumis*, which had only one consensus sequence. For tetraploid *Stipa capillata* L. and *S. tirsa* Steven (both 2*n* = 44; Appendix 2) we identified two different *Acc1* copy types after analysing the clone sequences. For diploid *Achnatherum paradoxum* and *A. sibiricum* (both 2*n* = 24) as well as polyploid *Nassella trichotoma* (2*n* = 36, 38; Appendix 2), only one sequence with clear peaks in the chromatograms was identified from direct sequencing of the PCR products.

The topology of the trees inferred by ML, MP and BI analyses were largely identical although their statistical supports differed slightly. Fig. 2 shows trees with plastid and nuclear ITS DNA data reduced to a single accession per taxon. The complete phylograms with all studied accessions are presented in suppl. Figs. S1 and S2.

**Fig. 2.**
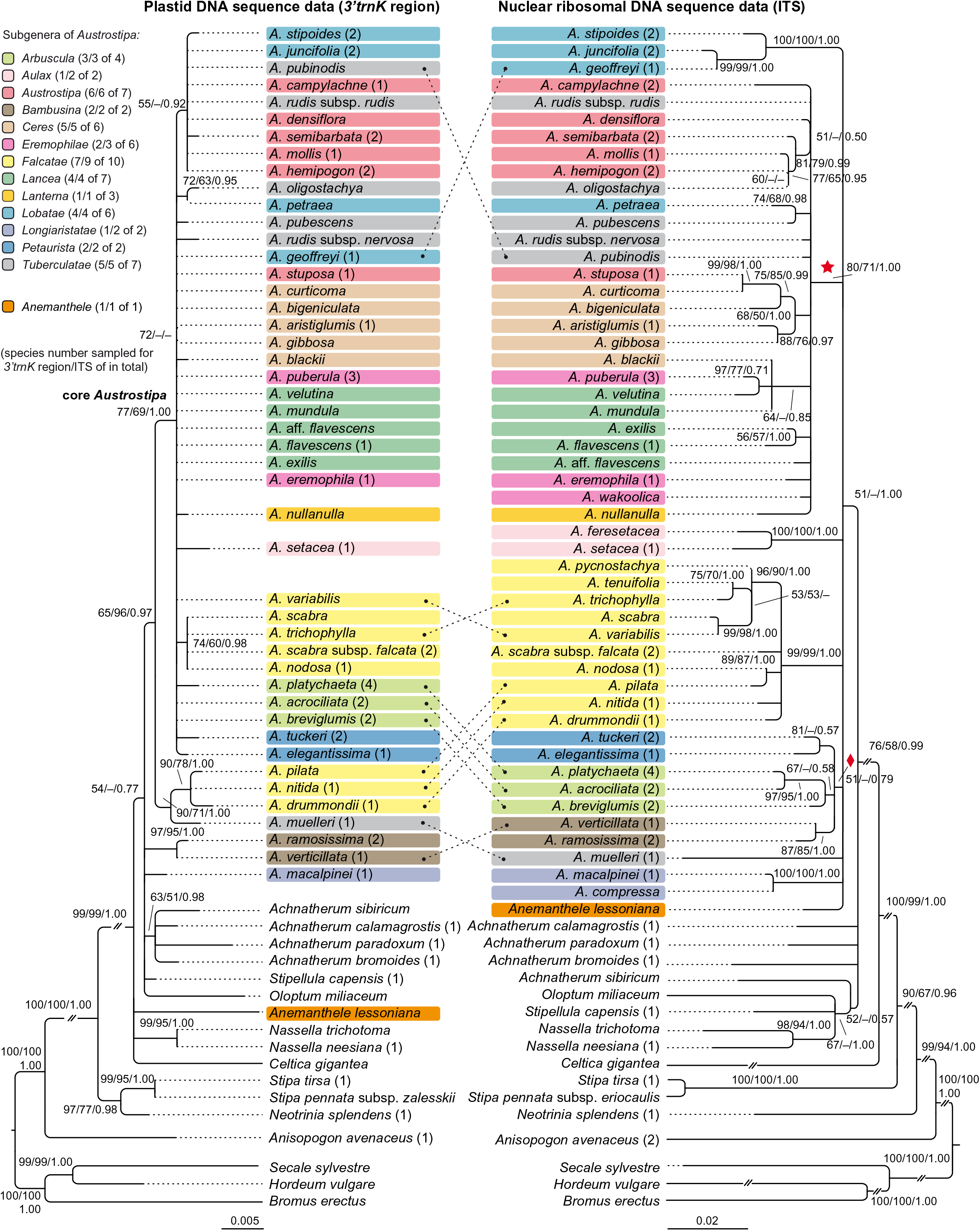
Maximum likelihood phylograms of *Austrostipa* species, *Anemanthele lessoniana* and exemplary other taxa of tribe Stipeae inferred from plastid (3’*trnK* region) and nuclear ribosomal (ITS1–5.8S gene–ITS2) DNA sequences with *Bromus erectus* (Bromeae), *Hordeum vulgare*, *Secale sylvestre* (both Triticeae) and *Anisopogon avenaceus* (Duthieeae) used as outgroup. Reduced datasets with each taxon represented by a single representative accession. ML and MP bootstrap support values ≥50% as well as Bayesian PP ≥0.5 are indicated on the branches. Clades with ML support <50% are collapsed. The subgenera of *Austrostipa* are marked by different colors. The asterisked clade in the ITS tree was recovered also in the *Acc1* gene tree of Fig. 3 (Copy types A and B) as well as the clade with diamond (Copy type A). Abbreviation: *A*. = *Austrostipa*.

#### Plastid DNA analysis – 3’trnK region

The plastid 3’*trnK* region DNA sequence dataset (sequence lengths 579*–*798 bp) for 63 taxa of the reduced dataset (each species or subspecies represented by only one accession) included 832 aligned positions, of which 139 were variable (17%) and 59 parsimony-informative (7.0%).

After branching of the outgroup taxa (*Bromus erectus* Huds., *Hordeum vulgare*, *Secale sylvestre* Host and *Anisopogon avenaceus* R.Br. next to the stipoid taxa), a clade formed by *Neotrinia splendens* (Trin.) M.Nobis, P.D.Gudkova & A.Nowak, *Stipa pennata* L. and *S. tirsa* was sister to all other Stipeae sampled (100/100/1.00) (Fig. 2). *Anemanthele lessoniana* (Steud.) Veldkamp, *Celtica gigantea* (Link) F.M.Vázquez & Barkworth, *Nassella neesiana* (Trin. & Rupr.) Barkworth, and *N. trichotoma* stood more or less in a polytomy with *Oloptum miliaceum* (L.) Röser & Hamasha, *Stipellula capensis* (Thunb.) Röser & Hamasha, a clade of four *Achnatherum* species (*A. bromoides* (L.) P.Beauv., *A. calamagrostis* (L.) P.Beauv., *A. paradoxum*, *A. sibiricum*), three *Austrostipa* species (*A. macalpinei* (Reader) S.W.L.Jacobs & J.Everett, *A. ramosissima* (Trin.) S.W.L.Jacobs & J.Everett, *A. verticillata* (Nees ex Spreng.) S.W.L.Jacobs & J.Everett) and the remainder of the latter genus (Fig. 2). *Austrostipa drummondii* (Steud.) S.W.L.Jacobs & J.Everett, *A. muelleri* (Tate) S.W.L.Jacobs & J.Everett, *A. nitida* (Summerh. & C.E.Hubb.) S.W.L.Jacobs & J.Everett and *A. pilata* (S.W.L.Jacobs & J.Everett) S.W.L.Jacobs & J.Everett formed a supported clade (97/95/1.00), which was opposed to a large polytomy of all other species studied (77/69/1.00), here termed ‘core *Austrostipa*’ clade. Among them, *A. oligostachya* (Hughes) S.W.L.Jacobs & J.Everett and *A. petraea* (Vickery) S.W.L.Jacobs & J.Everett formed a moderately supported species pair (72/63/0.95). Groups of varying size and mostly low support were formed by (1) *A. nodosa* (S.T.Blake) S.W.L.Jacobs & J.Everett, *A. scabra*, *A. scabra* subsp. *falcata* (Hughes) S.W.L.Jacobs & J.Everett, *A. trichophylla* (Benth.) S.W.L.Jacobs & J.Everett (74/60/0.98), and (2)*A. campylachne* (Nees) S.W.L.Jacobs & J.Everett, *A. densiflora* (Hughes) S.W.L.Jacobs & J.Everett, *A. hemipogon* (Benth.) S.W.L.Jacobs & J.Everett, *A. juncifolia* (Hughes) S.W.L.Jacobs & J.Everett, *A. mollis* (R.Br.) S.W.L.Jacobs & J.Everett, *A. pubinodis* (Trin. & Rupr.) S.W.L.Jacobs & J.Everett, *A. rudis* (Spreng.) S.W.L.Jacobs & J.Everett subsp. *rudis*, *A. semibarbata* (R.Br.) S.W.L.Jacobs & J.Everett and *A. stipoides* (55/‒/0.92) (Fig. 2).

#### Nuclear DNA – ITS

The reduced nr ITS DNA sequence dataset for 68 taxa (each species or subspecies represented by only one accession) included 644 aligned positions (sequence lengths 500*–*627 bp), of which 263 (41%) were variable and 185 (29%) parsimony-informative.

Following the outgroup taxa (*Bromus erectus*, *Hordeum vulgare*, *Secale sylvestre* and *Anisopogon avenaceus* next to the stipoid taxa), representatives from several Stipeae genera including *Achnatherum*, *Celtica*, *Nassella*, *Neotrinia*, *Oloptum*, *Stipa* and *Stipellula* were next to a clade of *Anemanthele* and *Austrostipa* (51/‒/1.00; Fig. 2). Overall resolution within this clade was low, however, several supported groups of species or species pairs could be discerned, for example, that of (1) *A. compressa* (R.Br.) S.W.L.Jacobs & J.Everett, *A. macalpinei* (100/100/1.00), (2) *A. ramosissima*, *A. verticillata* (87/85/1.00), (3) *A. acrociliata* (Reader) S.W.L.Jacobs & J.Everett, *A. breviglumis* (J.M.Black) S.W.L.Jacobs & J.Everett, *A. platychaeta* (Hughes) S.W.L.Jacobs & J.Everett (67/‒/0.58), (4) *A. elegantissima* (Labill.) S.W.L.Jacobs & J.Everett, *A. tuckeri* (F.Muell.) S.W.L.Jacobs & J.Everett (87/‒/0.57), (5) a larger clade encompassing *A. drummondii*, *A. nitida*, *A. nodosa*, *A. pilata*, *A. pycnostachya* (Benth.) S.W.L.Jacobs & J.Everett, *A. scabra*, *A. scabra* subsp. *falcata, A. trichophylla*, *A. tenuifolia* (Steud.) S.W.L.Jacobs & J.Everett, *A. variabilis* (Hughes) S.W.L.Jacobs & J.Everett (99/99/1.00), (6) *A. feresetacea* (Vickery, S.W.L.Jacobs & J.Everett) S.W.L.Jacobs & J.Everett, *A. setacea* (R.Br.) S.W.L.Jacobs & J.Everett (100/100/1.00) and (7) *A. geoffreyi* S.W.L.Jacobs & J.Everett, *A. juncifolia*, *A. stipoides* (100/100/1.00). The remaining species formed a larger clade asterisked in Fig. 2 (80/71/1.00). More or less supported internal clades consisted of (1) *A. exilis* (Vickery) S.W.L.Jacobs & J.Everett and *A. flavescens* (Labill.) S.W.L.Jacobs & J.Everett (56/57/1.00), (2) *A. puberula* (Steud.) S.W.L.Jacobs & J.Everett and *A. velutina* (Vickery, S.W.L.Jacobs & J.Everett) S.W.L.Jacobs & J.Everett (97/77/0.71) together with *A*. *blackii* (C.E.Hubb.) S.W.L.Jacobs & J.Everett and *A*. *mundula* (J.M.Black) S.W.L.Jacobs & J.Everett (64/‒/0.85), (3) *A*. *aristiglumis* (F.Muell.) S.W.L.Jacobs & J.Everett and *A*. *gibbosa* (Vickery) S.W.L.Jacobs & J.Everett (88/76/0.97), *A*. *curticoma* (Vickery) S.W.L.Jacobs & J.Everett and *A. stuposa* (Hughes) S.W.L.Jacobs & J.Everett (99/98/1.00) together with *A. bigeniculata* (Hughes) S.W.L.Jacobs & J.Everett (68/76/1.00), *A. petraea* and A. *pubescens* (R.Br.) S.W.L.Jacobs & J.Everett (74/68/0.98) and (6) *A*. *hemipogon*, *A*. *mollis*, *A*. *oligostachya*, *A*. *semibarbata* (81/79/0.99).

#### Nuclear DNA – single-copy locus *Acc1*

The *Acc1* DNA dataset of 73 sequences from 33 species and subspecies included 1512 aligned positions (sequence lengths 1329–1458 bp), of which 568 were variable (38%) and 348 parsimony-informative (23%).

The phylogram of the single-copy region *Acc1* with *Bromus inermis*, *Henrardia persica*, *Hordeum chilense* and *H. vulgare* as outgroup showed species of *Stipa* s.str. basal to a clade comprising all other stipoid taxa (*Achnatherum*, *Anemanthele*, *Austrostipa, Nassella*; Fig. 3). We identified two different copy types of *Acc1* for The tetraploids *S. capillata* and *S. tirsa* (Fig. 3) had two copies of *Acc1*,which resulted in formation of two separate clade that were not sister(both 100/100/1.00). One of these *Stipa* copy type clades was sister to the strongly supported clade with all *Acc1* copy types of *Achnatherum*, *Anemanthele* and *Austrostipa* (100/100/1.00). The *Acc1* copy types of *Anemanthele* and *Austrostipa* segregated into two lineages, Copy type A and B in Fig. 3. The diploids of *Achnatherum* (*A. paradoxum*, *A. sibiricum*) as well as polyploid *Nassella trichotoma* (see Appendix 2) had only a single copy type of *Acc1*. They formed a basal grade to the well-supported Copy type A Australasian clade of *Anemanthele* and *Austrostipa* (87/74/1.00).

**Fig. 3.**
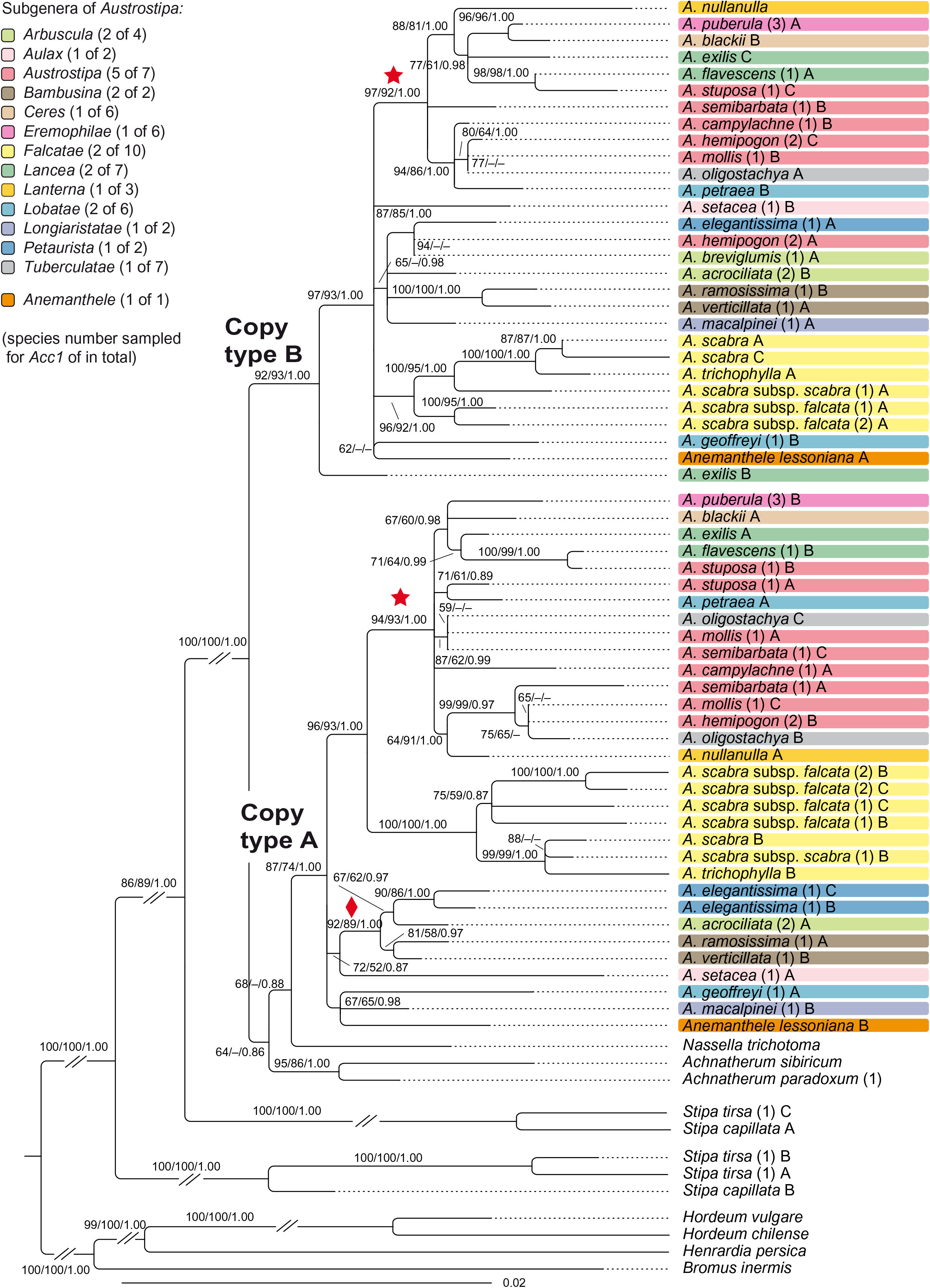
Maximum likelihood phylogram of *Austrostipa* species, *Anemanthele lessoniana* and exemplary other taxa of tribe Stipeae (*Achnatherum* spp., *Nassella trichotoma*, *Stipa* spp.) inferred from DNA sequences of the nuclear single-copy locus *Acc*1 (exon 6–13) with *Bromus inermis* (Bromeae), *Henrardia persica* and *Hordeum* spp. (both Triticeae) used as outgroup. ML and MP bootstrap support values ≥50% as well as Bayesian PP ≥0.5 are indicated on the branches. Clades with ML support <50% are collapsed. The subgenera of *Austrostipa* are marked by different colors. The asterisked clade within the Copy type A and B clades was recovered also in the ITS tree of Fig. 2 as well as the clade with diamond. Abbreviation: *A*. = *Austrostipa*.

Copy type A clade comprised three subclades in a polytomy, namely (1) *Anemanthele lessoniana*, *Austrostipa geoffreyi* and *A. macalpinei* (67/65/0.98), (2) *A. acrociliata*, *A. elegantissima*, *A. ramosissima*, *A. setacea* and *A. verticillata* (92/89/1.00) and (3) a clade (96/93/1.00) with all six *A. scabra* accessions plus *A. trichophylla* (100/100/1.00) and another clade (94/93/100) with some well-supported minor lineages. Copy type B clade showed *A. exilis* sister to a larger polytomy encompassing *Anemanthele lessoniana* and the remaining species of *Austrostipa*, organized in several minor lineages.

### Morphological analyses

In all but 6 of the 34 taxa of *Austrostipa* examined, the lemma epidermal patterns (LEP) were relatively uniform (Figs. 4a‒u, 5a‒d, f; Appendix 1; *Austrostipa acrociliata*, *A. mundula* and *A. rudis* subsp. *rudis* not shown) and typical of achnatheroid grasses as seen, for example, in *Achnatherum*, *Anemanthele, Jarava* Ruiz & Pav. or *Stipellula* (Fig. 5l‒o) This so-called maize-like LEP is characterized by wider than long, short or square to rectangular fundamental cells, with undulate to almost straight side walls. Silica bodies were very frequent, ovate to elongate, densely packed and regularly alternate with fundamental cells. In the remaining six species, namely *A. stipoides* (subg. *Lobatae* S.W.L.Jacobs & J.Everett), *A. elegantissima* and *A. tuckeri* (subg. *Petaurista* S.W.L.Jacobs & J.Everett); *A. ramosissima* and *A. verticillata* (subg. *Bambusina*); and *A. pubescens* (subg. *Tuberculatae*) (Fig. 5e, g‒k), the LEP was distinctively different and more similar to the so-called saw-like LEP found in such Old World genera as *Stipa, Neotrinia*, *Orthoraphium* Nees and *Ptilagrostis* (Fig. 5s‒y). The *Stipa*-like LEP was observed in species from subgenera *Petaurista* and *Bambusina* as well as in *Austrostipa pubescens* from subg. *Tuberculatae* (Fig. 5e, g‒j). It is characterized by having short cells with hooks that alternate with square or rectangular fundamental cells and ovate silica bodies. The *Ptilagrostis*-like LEP observed in *A. stipoides* (Fig. 5k) is characterized by its rectangular to elongated fundamental cells (1.5‒4 times as long as wide), which often alternate with silica cells and cork cells as well as sometimes scattered hooks. The elongated silica cells were often associated with cork cells and frequently had 1‒4 constrictions.

**Fig. 4.**
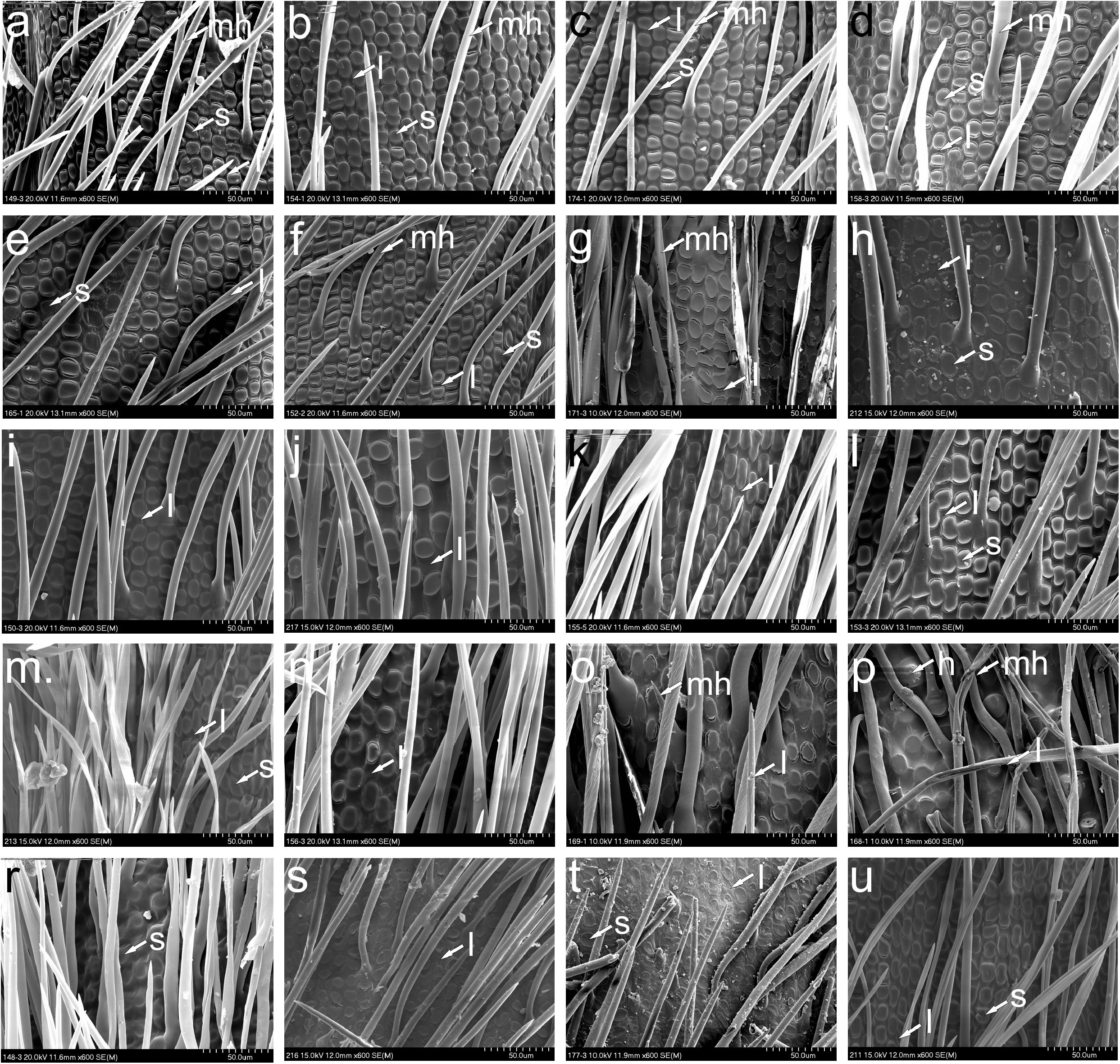
SEM morphology of lemma epidermal patterns in *Austrostipa*. **a,** Austrostipa nitida; b, *A. nodosa*; **c,** A. scabra subsp*. scabra*; **d,** A. scabra subsp*. falcata*; **e,** A. variabilis; **f,** A. drummondii; **g,** A. stuposa; **h,** A. campylachne; **i,** A. mollis; **j,** A. densiflora; **k,** A. hemipogogon; **l,** A. semibarbata; **m,** A. eremophila; **n,** A. setacea; **o,** A. bigeniculata; **p,** A. aristiglumis; **r,** A. gibbosa; **s,** A. compressa; **t,** A. macalpinei; **u,** A. flavescens. Abbreviations: l = long cell (fundamental cell); s = silica cell (silica body); h = hook; mh = macrohair. The list of specimens studied is presented in Appendix 1.

**Fig. 5.**
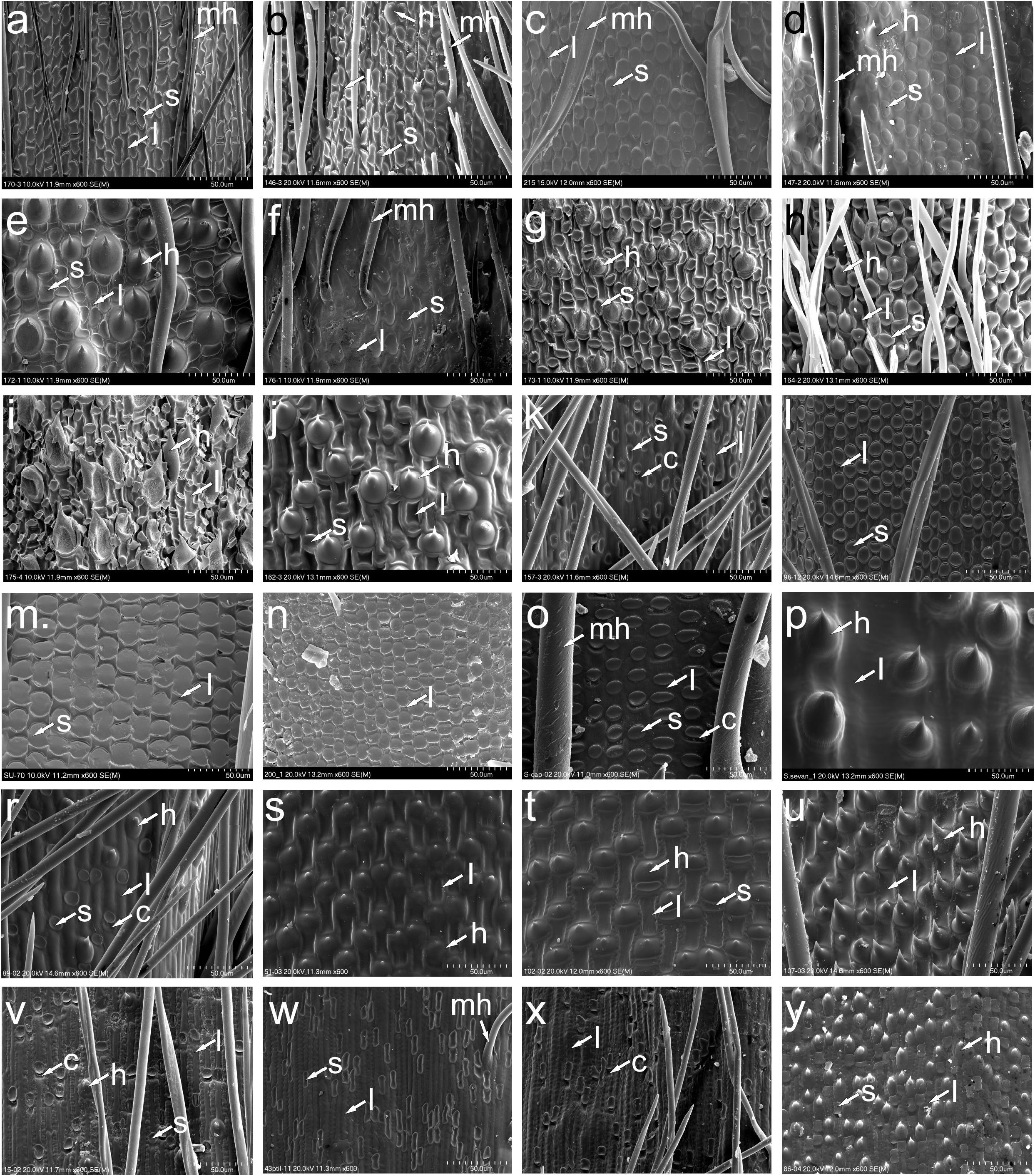
SEM morphology of lemma epidermal patterns of *Austrostipa*. **a,** Austrostipa breviglumis; **b,** A. platychaeta; **c,** A. rudis subsp*. australis*; **d,** A. pubinodis; **e,** A. pubescens; **f,** A. muelleri; **g,** A. ramosissima; **h,** A. verticillata; **i,** A. tuckeri; **j,** A. elegantissima; **k,** A. stipoides; **l,** Achnatherum calamagrostis; **m,** A. bromoides; **n,** A. paradoxum; **o,** Stipellula capensis; **p,** Nassella neesiana; **r,** Macrochloa tenacissima; **s,** Stipa tirsa; **t,** S. kirghisorum; **u,** S. drobovii; **v,** Neotrinia splendens; **w,** Ptilagrostis mongholica; **x,** P. concinna; **y,** Orthoraphium roylei. Abbreviations: l = long cell (fundamental cell); s = silica cell (silica body); c = cork cell; h = hook; mh = macrohair. The list of specimens studied is presented in Appendix 1.

The species representing subgenera *Bambusina, Lobatae* and *Petaurista* were well separated from all other species in the cluster analysis (UPGMA), which was performed on a combined macro- and micromorphological 34-taxa dataset (Fig. 6; Table S2). Similar results were obtained when the micro- and macromorphological characters were analyzed separately (34- and 65-taxa set, respectively; suppl. Figs. S3, S4; Tables S1, S2). A unique LEP (*A. stipoides*) and presence of distinct lobes on the top of the lemma (*A. juncifolia*, *A. geoffreyi, A. petraea*) separated these four species of subg. *Lobatae* from the remaining subgenera of *Austrostipa* (Fig. 6, suppl. Figs. S3, S4). A further cluster was formed by representatives of subgenera *Petaurista* and *Bambusina*, which had saw-like LEP (Fig. 6, suppl. Fig. S3). Additionally, a unique morphology of the inflorescences characterized by long pilose branches, which occurred exclusively in *A. elegantissima* and *A. tuckeri* (subg. *Bambusina*), resulted in a clear separation from the other subgenera of *Austrostipa* (Fig. 6, suppl. Fig. S4). Due to its long apical lemma lobes also *A. muelleri* was well-distinguished not only from the other representatives of subg. *Tuberculatae* but from all other species with achnatheroid (maize-like) LEP, whereas the species of the remaining subgenera of *Austrostipa* with achnatheroid LEP were grouped in several clusters in accordance to each of the three performed analyses, with rather weekly noticeable subgeneric ordination (Fig. 6, suppl. Figs. S3, S4).

**Fig. 6.**
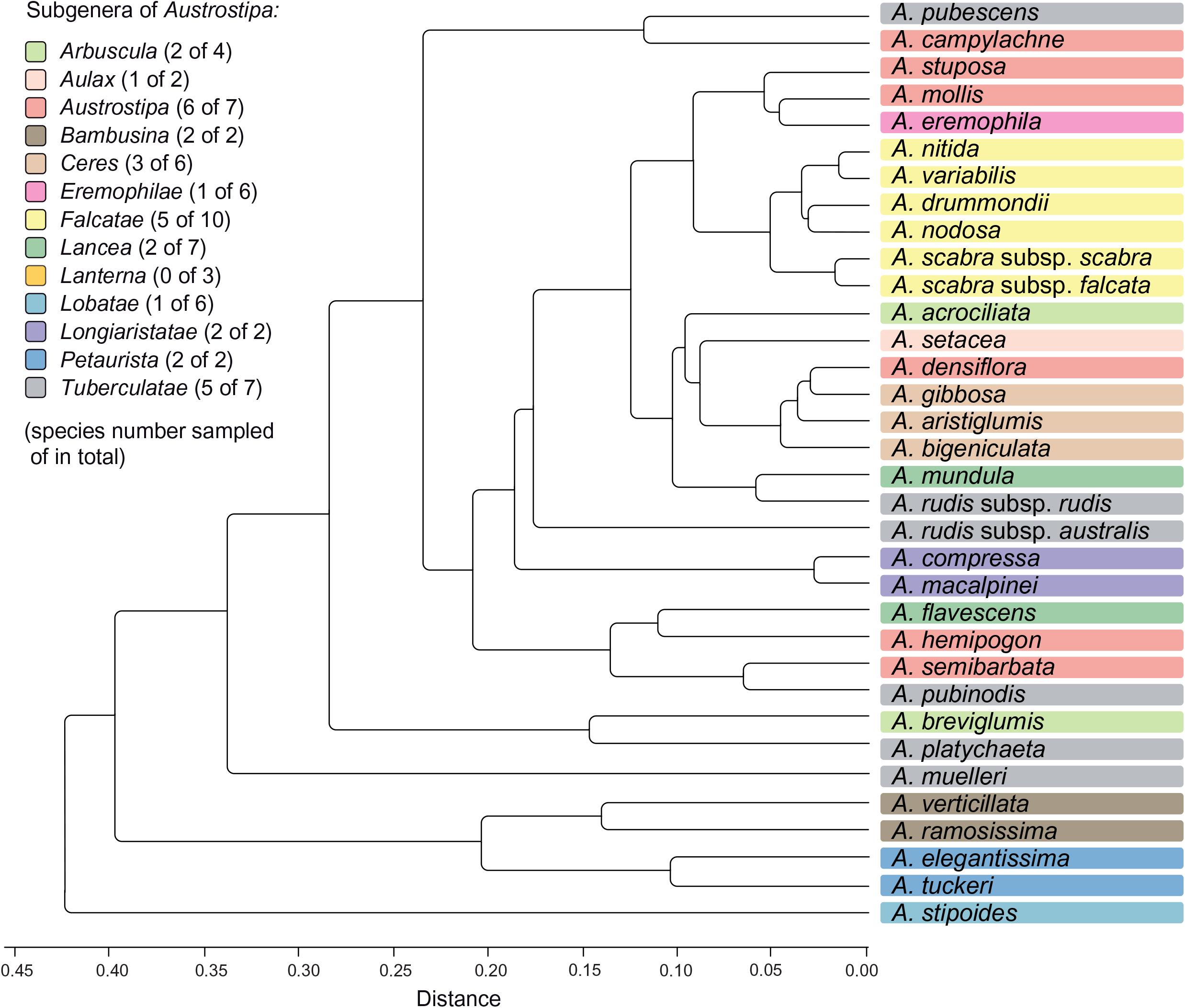
Cluster analysis (UPGMA) performed on eight macro- and four micromorphological characters for 34 *Austrostipa* taxa. See suppl. Table S2 for the data matrix evaluated. The subgenera of *Austrostipa* are marked by different colors. Abbreviation: *A*. = *Austrostipa*.

## DISCUSSION

### Molecular phylogenetic delineation of *Austrostipa*

Monophyly of *Austrostipa* was not clearly supported by any of the three DNA regions we investigated (plastid 3’*trnK* region: Fig. 2, suppl. Fig. S1; nr ITS region: Fig. 2, suppl. Fig. S2 and the single-copy locus *Acc1*: Fig. 3).

The plastid DNA trees showed *Austrostipa* paraphyletic. Most species (36/43) grouped in a single clade (consisting of supported ‘core *Austrostipa*’ and a supported clade of *A. drummondii*, *A. muelleri*, *A. nitida*, *A. pilata*), which stood in a polytomy that included the three remaining species of *Austrostipa* (*A. macalpinei*, *A. ramosissima* and *A. verticillata*) and three Eurasian stipoid genera (*Achnatherum*, *Oloptum* and *Stipellula*). *Anemanthele* was not part of this polytomy, but of the next lower one which also included the two representatives of the primarily South American genus *Nassella* and the western Mediterranean species *Celtica gigantea*. Possible explanations for the failure of the plastid data to support, even weakly, the monophyly of *Austrostipa* include incomplete lineage sorting (ILS) affecting the inheritance of plastids or genetic introgression from the Eurasian species into the three *Austrostipa* species in the lowest *Austrostipa-*containing clade (*A. macalpinei*, *A. ramosissima, A. verticillata*). This last seems unlikely, given the present day distribution of the species involved.

Nuclear ITS grouped *Austrostipa* (all species) and *Anemanthele* in a single clade but support was minimal for the relationship (see reduced dataset with each species represented only by a single accession in Fig. 2 and suppl. Fig. S2 with all accessions studied).

*Austrostipa* and *Anemanthele* were alike in having two different copies of the nuclear gene *Acc1* (Copy types A and B). These resolved together into two separate clades (Fig. 3). The copies obtained for *Achnatherum paradoxum*, *A. sibiricum* and *Nassella trichotoma* were close to the Copy type A clade; those for the two species of *Stipa* included (*S. capillata*, *S. tirsa*) divided likewise into two copy types, both of which were outside the two *Austrostipa* clades (Fig. 3).

Our results, in failing to contradict or providing only weak support for the monophyly of *Austrostipa* and the closer relationship of *Austrostipa* to *Anemanthele* rather than non-Australasian stipoids, basically agrees with the findings of several previous studies regardless of taxon sampling (Jacobs & al., 2000, 2007; Barkworth & al., 2008; Romaschenko & al., 2010, 2012; Syme, 2011; Syme & al., 2012; Hamasha & al., 2012). The odd results for *Austrostipa stipoides* reported in two studies (Jacobs & al. 2007; Barkworth & al. 2008), which placed the species distant from other species of the genus, were from duplicate collections (Barkworth & al. 2007: 725) and were not corroborated by this study, in which three different collections were used (see Appendix 1). Their plastid and nuclear DNA sequences clustered with those of other *Austrostipa* species (Figs. 2, 3, suppl. Figs. S1, S2), as did sequences from the specimens of *A. stipoides* studied by Syme (2011) and Syme & al. (2012).

### Phylogenetic differentiation in *Austrostipa* and taxonomy

All but one of the subgenera of *Austrostipa* were represented by two representatives for at least one of the sequences we examined (Table 1). The exception was (1) subg.

*Lanterna*, which means we cannot comment on its monophyly.

#### Comparison of plastid and nr ITS tree

The peculiar position of *A. ramosissima* and *A. verticillata*, the two species of (2) subg. *Bambusina* and *A. macalpinei* of (3) subg. *Longiaristatae*( altogether 2 species) in the plastid DNA tree was not reflected in the ITS tree, however, both subgenera were supported as monophyletic considering *A. compressa* (subg. *Longiaristatae*), which was sampled only for nr ITS DNA.

The smaller clade of the remainder of *Austrostipa* species in the plastid DNA tree comprised species of two different subgenera, namely three species of (4) subg. *Falcatae* (*A. drummondii*, *A. nitida*, *A. pilata*), the largest subgenus of *Austrostipa* with altogether 10 species, and *A. muelleri* of (5) subg. *Tuberculatae* (see also below). Both subgenera were represented also in the ‘core *Austrostipa*’ clade of the plastid DNA tree (subg. *Falcatae*: *A. nodosa*, *A. scabra*, *A. scabra* subsp. *falcata, A. trichophylla*, *A. variabilis*; subg.

*Tuberculatae*: *A. pubescens*, *A. pubinodis*, *A. rudis* subsp. *rudis* and subsp. *nervosa* (Vickery) S.W.L.Jacobs & J.Everett; Fig. 2, suppl. Fig. S1). *Falcatae*, however, were resolved in the ITS tree as monophyletic, whereas *Tuberculatae* were highly polyphyletic. In other words, subg. *Falcatae* is characterized by remarkable cytonuclear discordance, having at least two different chloroplast ‘types’.

The weakly supported clade marked by a diamond in the ITS tree of Fig. 2 united three subgenera. It comprised both species of monophyletic (2) subg. *Bambusina* (*A. ramosissima*, *A. verticillata*), the three sampled species of monophyletic (6) subg. *Arbuscula* (*A. acrociliata*, *A. breviglumis*, *A. platychaeta*; altogether 4 species) and both species of (7) subg. *Petaurista* (*A. tuckeri*, *A. elegantissima*). This clade was not recovered in the plastid DNA tree. In this instance, the occurrence of altogether three different plastid types has to be noted, one of which is shared with that of Stipeae outgroup genera (see above).

Except for small (8) subg. *Aulax* with both of its species sampled for ITS (*A. feresetacea*, *A. setacea*), none of the remaining *Austrostipa* subgenera encompassing several species resolved in our plastid and ITS DNA analyses as monophyletic. They were para- or polyphyletic or their species were placed in polytomies (Fig. 2, suppl. Figs. S1, S2) as seen in (6) subg. *Arbuscula* (3 of altogether 4 species sampled: *A. acrociliata*, *A. breviglumis*, *A. platychaeta*), (9) subg. *Austrostipa* (6 of 7 species sampled: *A. campylachne*, *A. densiflora*, *A. hemipogon*, *A. mollis*, *A. semibarbata*, *A. stuposa*), (10) subg. *Ceres* S.W.L.Jacobs & J.Everett (5 of 6 species sampled*: A. aristiglumis*, *A. bigeniculata*, *A. blackii*, *A. curticoma*, *A. gibbosa*), (11) subg. *Eremophilae* S.W.L.Jacobs & J.Everett (3 of 6 species sampled: *A. eremophila* (Reader) S.W.L.Jacobs & J.Everett, *A. puberula*, *A. wakoolica* (Vickery, S.W.L.Jacobs & J.Everett) S.W.L.Jacobs & J.Everett), (12) subg. *Lancea* S.W.L.Jacobs & J.Everett (4 of 7 species sampled: *A. exilis*, *A. flavescens*, *A.* aff. *flavescens*, *A. mundula*, *A. velutina*), (13) subg. *Lobatae* (4 of 6 species sampled: *A. geoffreyi*, *A. juncifolia*, *A. petraea*, *A.stipoides*) and (5) subg. *Tuberculatae* (5 of 7 species sampled: *A. muelleri* mentioned above, *A. oligostachya*, *A. pubescens*, *A. pubinodis*, *A. rudis* subsp. *rudis* and subsp. *nervosa*).

#### Single-copy locus *Acc1*

The analyses of the *Acc1* sequences corroborated monophyly of subgenera *Bambusina* and *Falcatae*, whereas subgenera *Arbuscula*, *Austrostipa*, *Lancea* and *Lobatae* were non-monophyletic (Fig. 3). Within Copy type A clade, the asterisked clade supported by 94/93/1.00 (Fig. 3) comprised species belonging to subg. *Austrostipa* (*A. campylachne*, *A. hemipogon*, *A. mollis*, *A. semibarbata*, *A. stuposa*), subg. *Ceres* (*A. blackii*), subg. *Eremophilae* (*A. puberula*), subg. *Lancea* (*A. exilis*, *A. flavescens*), subg. *Lanterna* (*A. nullanulla* (J.Everett & S.W.L.Jacobs) S.W.L.Jacobs & J.Everett), subg. *Lobatae* (*A. petraea*) and subg. *Tuberculatae* (*A. oligostachya*). This clade was largely reflected also in the Copy type B topology (asterisked; 97/92/1.00). *Austrostipa exilis* (accession shown to be tetraploid with 2*n* = 44; Appendix 2; Winterfeld & al., 2015) and *A. hemipogon* (accession shown to be hexaploid with 2*n* = 66; Appendix 2; Winterfeld & al., 2015) have additional copies of *Acc1* gene Copy type B. That for *A. exilis* was placed external to all other *Australasian* stipoids in the tree (Fig. 3). The asterisked clades in *Acc1* Copy type A and B clades corresponded well with the asterisked clade supported by 80/71/1.00 in the ITS tree (Fig. 2), thus there is consistent phylogenetic signal in both nuclear markers studied.

#### Delineation and relationship of subgenera

Despite limited phylogenetic resolution recovered from the sequenced plastid and nuclear DNA loci as well as the combined macro- and micromorphological analysis, some conclusions can be drawn with respect to the infrageneric taxonomy of *Austrostipa* and the validity of the altogether 13 subgenera presented in Vickery & al. (1986), Jacobs & Everett (1996) and Everett & al. (2009), all of which were included in this study.

(1) The small subgenera *Longiaristatae* S.W.L.Jacobs & J.Everett (both species sampled, plastid DNA data missing for *A. compressa*) and *Bambusina* (both species sampled) belong to the early-branching lineages within *Austrostipa* considering the plastid DNA tree. Subg. *Bambusina* assembled together with subgenera *Petaurista* (both species sampled) and *Arbuscula* (three of four species sampled) in the same ITS and in Copy type A clades of the *Acc1* gene analyses marked by diamonds (Figs. 2, 3). The latter subgenera were placed in the ‘core *Austrostipa*’ clade of the plastid DNA tree distantly to the species of subg. *Bambusina*. Maintenance of subgenera *Petaurista* and *Arbuscula* is neither explicitly supported nor contradicted by our data. Thus, we argue that these four subgenera should remain unchanged.

(2) *Austrostipa muelleri* was placed distantly from all other taxa of subg. *Tuberculatae* (see below), in which it was accommodated (Jacobs & Everett, 1996; Everett & al., 2009). This deviating position was noted already previously (Jacobs & al., 2007: Fig. 4; Syme & al., 2012). We propose placing *A. muelleri* by itself in a new subgenus (see below New names and combinations).

(3) Subg. *Falcatae* (9 of 10 species sampled, plastid DNA data missing for *A. pycnostachya* and *A. tenuifolia*) was supported because of the ITS and *Acc1* DNA data (Copies A and B) but it disintegrated into two lineages of the plastid DNA phylogeny. One group of species possessed the ‘core *Austrostipa*’ plastid, the other shared a deviant plastid type with *A. muelleri* (Fig. 2; suppl. Fig. S1). The placement of *A. pycnostachya* in the ITS clade of subg. *Falcatae* (Fig. 2) supports the transfer of this species from subg. *Arbuscula*, in which it was placed by Jacobs & Everett (1996), to subg. *Falcatae* (Everett & al., 2009)

(4) Subg. *Aulax* S.W.L.Jacobs & J.Everett (both species sampled, plastid DNA data missing for *A. feresetacea*) and subg. *Lobatae* (4 of 6 species sampled) could be maintained after excluding *A. petraea* from the latter (Fig. 2). Segregation of *A. petraea* from the other species of subg. *Lobatae* was noted also by Syme & al. (2012). We found no support, however, for a placement of this species in subg. *Aulax* as suggested by the latter study (see Syme & al., 2012: Fig. 1).

(5) The high-support clades asterisked in the ITS and *Acc1* phylograms (Figs. 2, 3) encompass, apart from *A. petraea*, the species of subgenera *Austrostipa* (6 of 7 species sampled), *Ceres* (5 of 6 species sampled), *Eremophilae* (5 of 6 species sampled), *Lancea* (six of seven species sampled), *Lanterna* (1 of 3 species sampled) and *Tuberculatae* (5 of 7 species sampled). The asterisked clades showed several sister species relationships and minor lineages within and between subgenera (see above), but none of the subgenera mentioned was resolved as separate lineage, which is in agreement with the trees presented by Syme & al. (2012). For the time being it seems best to assign all these subtribes to a single, expanded and most likely monophyletic subtribe *Austrostipa*. This suggestion, however, should not be interpreted as attempt to supersede traditional morphology- by molecular phylogenetics-based taxonomic concepts. It is rather a contribution to obtain monophyletic taxa, which can serve as reliable units to address the interesting questions of character evolution or biogeography in *Austrostipa*, which have been barely touched upon to date.

Some of our suggestions for classification are not new, having been made in previous molecular phylogenetic studies of *Austrostipa*, for example, the maintenance of subgenera *Falcatae* (Jacobs & al., 2007; Bustam, 2010; Syme & al., 2012), *Longiaristatae* and *Lobatae* (Jacobs & al., 2007; Syme & al., 2012), the broadening of subg. *Austrostipa* to include also subgenera *Tuberculatae* (Jacobs & al., 2007; Syme & al., 2012) and *Eremophilae* (Syme & al., 2012), but our data do not support combining subgenera *Arbuscula* and *Bambusina*, a suggestion based on their similar habit (Jacobs & al., 2007).

In summary, we suggest to divide *Austrostipa* into the following nine subgenera (with number of species): *Arbuscula* (4), *Aulax* (2), *Austrostipa* (36), *Bambusina* (2), *Falcatae* (10), *Lobatae* (5), *Longiaristatae* (2), *Petaurista* (2) and *Paucispiculatae*, subg. nov., with *A. muelleri* (1).

### Basic chromosome numbers and whole-genome duplications

#### Austrostipa and Anemanthele

Our previous study on chromosome numbers and karyotypes in *Austrostipa* and *Anemanthele* (Winterfeld & al., 2015) established the somatic chromosome numbers of 2*n* = 44 in 18 and 2*n* = 66 in seven *Austrostipa* species as well as 2*n* = 44 in *Anemanthele lessoniana*, corroborating the available earlier chromosome counts in *Austrostipa stipoides* (2*n* = 44; Murray & al., 2005) and *Anemanthele lessoniana* (2*n* = 40–44; Dawson & Beuzenberg, 2000; Edgar & Connor, 2000). In some accessions a certain degree of aneusomaty was noted, for example, 2*n* = 65, 66, 68, 70 in *Austrostipa semibarbata*, but usually the chromosome number showed less variation or was uniform in the metaphase plates of each accession studied (Winterfeld & al., 2015). *Austrostipa* and *Anemanthele* thus encompass consistently polyploids with a basic chromosome number of *x* = 11. Apart from the overall similarity of their karyotypes (Winterfeld & al., 2015), this common basic number supports a close relationship of both genera and makes a common ancestry of *Austrostipa* and *Anemanthele* likely, in addition to the relationship shown by the molecular phylogenetic data (Figs. 2, 3) (Jacobs & al., 2007; Romaschenko & al., 2012).

#### Monoploid chromosome number variation in Stipeae

*Austrostipa* and *Anemanthele* were placed in a clade, in which otherwise the basic chromosome number of *x* = 12 prevails (Clade A in Fig. 7), which supports recognizing *x* = 11 as a synapomorphic character of both genera. The basic number of *x* = 12 was found in the likely sister of *Austrostipa* and *Anemanthele*, namely a lineage formed by *Achnatherum* (2*n* = 2*x* = 24; rarely 2*n* = 28 and few polyploids; see Appendix 2) and *Oloptum* (usually 2*n* = 2*x* = 24), whereas *Stipellula* most likely deviates from *x* = 12. Various somatic chromosome numbers have been reported for *S. capensis* (2*n* = 18, ca. 34, 36; Appendix 2), 2*n* = 36 being the most frequent in the whole Mediterranean (Appendix 2). 2*n* = 18 appears to be trustworthy for an accession from Gran Canaria, Canary Islands (Borgen, 1970 using the synonym *Stipa retorta* Cav.), making a derived monoploid chromosome number of *x* = 9 strongly conceivable for this species with annual life form, which is unusual in Stipeae. Moreover, 2*n* = 28, possibly pointing to *x* = 7, was reported in its congener *S. parviflora* (Appendix 2). The clade of *Austrostipa*, *Anemanthele*, *Achnatherum*, *Oloptum* and *Stipellula* has highly polyploid, monospecific *Celtica* (usually 2*n* = 8*x* = 96; *x* = 12) as sister. Australian/New Zealand *Austrostipa* and *Anemanthele* therefore are related with a group of genera distributed in Eurasia, the Mediterranean and with few outliers in Tropical East and South Africa (Clayton, 1970, 1972; Freitag, 1989; Fish & al., 2015).

**Fig. 7.**
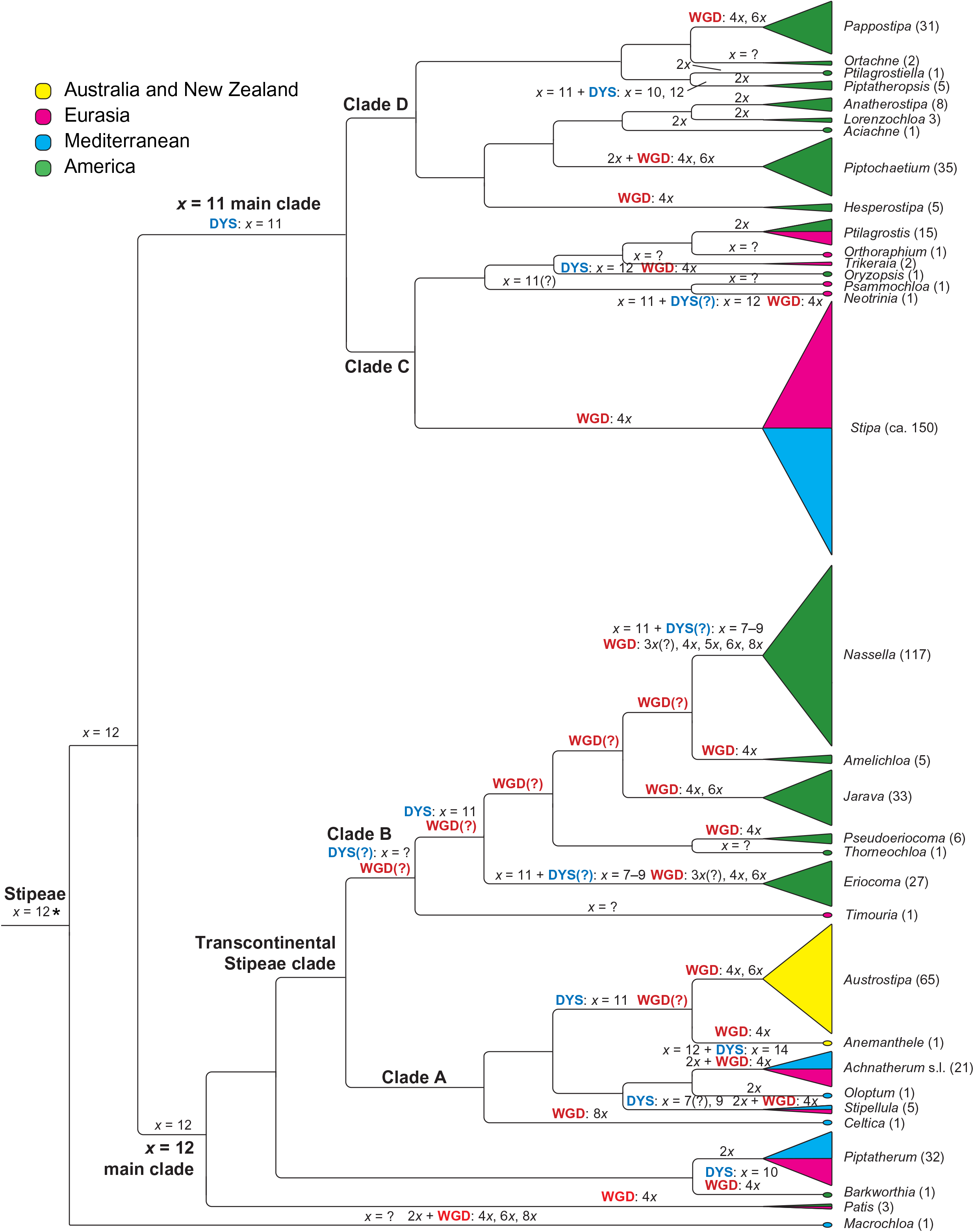
Basic chromosome numbers, dysploidy and whole-genome duplications in the evolution of tribe Stipeae mapped on a simplified molecular phylogenetic tree with the genera of Stipeae and their approximate sizes. The cladogram is modified from the plastid DNA tree of Romaschenko & al. (2012) and the plastid/nuclear DNA tree (concatenated data of congruent taxa) of Hamasha & al. (2012). The treatment of genera and estimated number of species largely follows Peterson & al. (2019). For chromosome data see Appendix 2. The species number of a genus is indicated after the genus name. Due to the unknown chromosome numbers in *Thorneochloa* and *Timouria* it is unclear at which split in Clade B whole-genome duplications occurred, potentially already in the common ancestor of all Clade B taxa. Abbreviations: DYS = dysploid chromosome number change, WGD = whole-genome duplication. The asterisk denotes our suggested most likely basic chromosome number (see Discussion).

Sister to all these Clade A genera is an almost exclusively and comparatively large New World lineage (Clade B) with a wide range of chromosome numbers (Appendix 2, Fig. 7). Chromosome numbers of *Eriocoma* (2*n* = 32, 34, 36, 40, 44, 48, 64, 66, 68, 70) and very speciose *Nassella* (2*n* = 26, 28, 32, 34, 36, 38, 42, 56, 58, 60, 64, 66, 70, 82, 88) seem to be based prevailingly on *x* = 11, implying the occurrence of 4*x*, 6*x*, 8*x* and possibly also 3*x* and 5*x* ploidy levels and a certain degree of aneusomatic variation. Assuming that chromosome numbers of 2*n* = 32‒34 are triploid numbers based on *x* = 11, the occurrence of triploids and pentaploids points towards heteroploid diploid-tetraploid and tetraploid-hexaploid hybridizations.

Also lower monoploid numbers such as *x* = 6 suggested by Stebbins & Love (1941: 379) for *Eriocoma*, and *x* = 7, 8 suggested by Barkworth (2007) for *Nassella* or *x* = 9 might occur in both genera, which means that accessions with 2*n* = 28, 32, 36, 38 would represent tetraploids or hexaploids. Given the branching order in the phylogenetic scheme of Fig. 7, such hypothetical monoploid chromosome sets of *Ericoma* and *Nassella* with x = 7‒9 have originated secondarily from *x* =11, the most likely original number of Clade B. In this phylogenetic context they do not give evidence of a sometimes suggested low ‘original’ base chromosome number of Stipeae (see Stebbins & Love, 1941, Tzvelev, 1977). Numbers reported in *Amelichloa* (2*n* = 40, 44, 46), *Jarava* (2*n* = 40, 44, 66) and *Pseudoeriocoma* Romasch., P.M.Peterson & Soreng (2*n* = 44, 46) seem to be based most likely on *x* = 11 if aneusomaty also plays some role here to explain the slightly varying chromosome numbers (Fig. 7). Chromosome numbers are unknown in monospecific North American *Thorneochloa* Romasch., P.M.Peterson & Soreng and in *Timouria* Roshev., a Central to East Asian outlier of this otherwise American clade. In summary, we suggest a secondary reduction of chromosome numbers in *Eriocoma* and *Nassella* via aneusomaty, whereas the basic chromosome number originally was *x* = 11 in Clade B and not lower (Fig. 7).

Both lineages of prevailingly *x* = 12 (Clade A) and *x* = 11 (Clade B), though with exceptions in *Stipellula* and species of *Eriocoma* and *Nassella*, constitute one of the major clades in Stipeae, which was named ‘Transcontinental Stipeae clade’ in Hamasha & al. (2012) to denote its representation on all continents including Australia and New Zealand (Fig. 7), and it is congruent with the ‘achnatheroid clade’ of Romaschenko & al. (2012).

The ‘Transcontinental Stipeae clade’ is allied with further genera of prevailingly *x* = 12, altogether forming the ‘*x* = 12 main clade’ of Stipeae (Fig. 7), namely comparatively species-rich and consistently diploid *Piptatherum* from the Mediterranean and Eurasia (32 species; 2*n* = 2*x* = 24; Appendix 2), East Asian/North American tetraploid *Patis* (3 species; 2*n* = 4*x* = 46, 48) and monospecific North American *Barkworthia* Romasch., P.M.Peterson & Soreng, in which 2*n* = 4*x* = 40 was reported, implying a monoploid set of *x* = 10 (Appendix 2) (Myers, 1947 citing an unpublished count of G.L. Stebbins).

The ‘*x* = 11 main clade’ (Fig. 7) represents the second main clade of Stipeae and includes *Stipa* s.str., by far the largest genus of this tribe. Exceptions from *x* = 11 are seemingly scarce in this clade but were noted for North American monospecific *Oryzopsis* Michx. (only *O. asperifolia* Michx. with probably *x* = 12), monotypic Asian *Neotrinia* (unclear whether x = 11 or 12) and some species of the North American genus *Piptatheropsis* Romasch., P.M.Peterson & Soreng (5 species; Appendix 1). In *P. pungens* (Torr. ex Spreng.) Romasch., P.M.Peterson & Soreng 2*n* = 22 and 24 were found, the latter number possibly caused by aneusomaty, whereas 2*n* = 2*x* = 20 was counted in mitotic and meiotic stages of two different accessions in *P. shoshoneana* (Curto & Douglass M.Hend.) Romasch., P.M.Peterson & Soreng (Curto & Henderson, 1998), which implies *x* = 10, and represents the lowest chromosome number of Stipeae in the New World as noted already by these authors.

The ‘*x* = 11 main clade’ is geographically clearly structured because it is divided into the Eurasian/Mediterranean Clade C (*Neotrinia*, *Orthoraphium*, *Psammochloa* Hitchc., *Stipa*, *Trikeraia* Bor) and the New World Clade D (*Aciachne* Benth., *Anatherostipa* (Hack. ex Kuntze) Peñail., *Hesperostipa* (M.K.Elias) Barkworth, *Lorenzochloa* Reeder & C.Reeder, *Ortachne* Nees, *Pappostipa*, *Piptatheropsis*, *Ptilagrostiella* Romasch., P.M.Peterson & Soreng; Fig. 7). There are only few exceptions since monospecific *Oryzopsis*, widely distributed in woodland of North America, is nested in Eurasian Clade C and *Ptilagrostis* occurs in mountainous to alpine landscapes of both Central Asia and western North America (Fig. 7).

The occurrence of two main clades in Stipeae, one clade primarily with *x* = 12 harboring also *Austrostipa* and *Anemanthele*, which are characterized by a derived number of *x* = 11, the other with primary *x* = 11 and few exceptions (see above *Stipellula*, *Oryzopsis*, species of *Piptatheropsis* and possibly also of *Eriocoma* and *Nassella*; Fig. 7), raises the question which one was the ‘original’ basic chromosome number of the whole tribe Stipeae. Due to the tree topology with monospecific *Macrochloa* Kunth, in which chromosome counts are equivocal, some suggesting *x* = 12 and others *x* = 11 or *x* = 10 (Appendix 2), as sister to the remainder of the tribe (Fig. 7), it cannot be reliably answered. We regard *x* = 12, as firstly proposed by Avdulov (1931: 130) and accepted also by Romaschenko & al. (2012), a bit more probable as original basic chromosome number of Stipeae than *x* = 11. Interestingly, this is supported mainly by chromosome numbers represented in the presumably closely related tribes of Stipeae (see below).

Further research into chromosome numbers of Stipeae, especially the re-examination of questionable counts contained in the older karyological literature as cited in reference works of Darlington & Wiley (1956) and Fedorov (1969), particularly the reported low numbers, seems worthwhile. This problem is frequently encountered with older chromosome counts, because counting was made using tissue sections, in which single chromosomes could easily become lost, instead of the nowadays employed and more reliable squashing technique.

#### The lowest chromosome number in Stipeae

2*n* = 18 from a diploid accession of *Stipellula capensis* from the Canary Islands (Borgen, 1970) seems to represent the lowest reliably known chromosome number of the whole tribe Stipeae (Appendix 2). The chromosome number so far considered as lowest in Stipeae (Curto & Henderson, 1998; Barkworth, 2007) refers to a Crimean accession of *Achnatherum bromoides* with 2*n* = 18 (Petrova, 1968), which was cited also in the reference works of Prokudin & al. (1977) and Agapova & al. (1993). This report appears to be questionable in view of the other chromosome counts available for *A. bromoides*, namely 2*n* = 24 (Ghukasyan, 2004) and repeatedly reported 2*n* = 28 (Vázquez & Devesa, 1996 and references therein). *Stipellula capensis* is otherwise known from many tetraploid populations (2*n* = 36; Appendix 2), which are widespread in the Mediterranean, and further numbers (see above and Appendix 2), which are currently difficult to interpret (eventually triploid hybrids, aneusomatic specimens and partly probably also erroneous counting).

#### Neighbor tribes of Stipeae also have *x* = 12

It is interesting to note that presumably close relatives of grass tribe Stipeae, which likewise belong to the rather early-diverging lineages of grass subfamily Pooideae (Schneider & al., 2009, 2011; Romaschenko & al., 2012; Saarela & al., 2015) seem to share the basic chromosome number of *x* = 12 with Stipeae (see above), even though only comparatively few counts are available. (1) monospecific *Ampelodesmos* Link (*A. mauritanicus* (Poir.) T.Durand & Schinz), regarded as either sole member of tribe Ampelodesmeae (GPWG, 2001; Soreng & al., 2017) or as morphologically anomalous genus of Stipeae (Decker, 1964; Barkworth, 2007; Schneider & al., 2009, 2011; Winterfeld & al., 2015) has 2*n* = 4*x* = 48 (Nilsson & Lassen, 1971; Schneider & al., 2011) or 2*n* = 8*x* = 96 (Myers, 1947 citing an unpublished count of G.L. Stebbins); (2) *Danthoniastrum compactum* (Boiss. & Heldr.) Holub, *Duthiea brachypodium* (P.Candargy) Keng & Keng f., *Sinochasea trigyna* Keng, *Stephanachne monandra* (P.C.Kuo & S.L.Lu) P.C.Kuo & S.L.Lu and *S. pappophorea* (Hack.) Keng (all tribe Duthieeae) all have 2*n* = 2*x* = 24 (Fedorov, 1969: 565 citing an unpublished count of L.A. Alexandrova; Winterfeld, 2006; Zhang & al., 2018; Schneider & al., 2011 with a discussion of a seemingly wrong previous chromosome counts of 2*n* = 14 in *Danthoniastrum compactum* of Kožuharov & Petrova, 1991), while *n* = 14 in was reported for *Duthiea bromoides* Hack. (Mehra & Sharma, 1975, 1977); and (3) monospecific *Phaenosperma* Munro ex Benth. (*P. globosum* Munro ex Benth.) of monogeneric tribe Phaenospermateae has 2*n* = 2*x* = 24 (Avdulov, 1931: 92, Tateoka, 1954, 1955, 1956; Schneider & al., 2011; Winterfeld & al., 2015; Zhang & al., 2018).

#### Whole-genome duplications in Stipeae

Although chromosome numbers are still unknown in a number of genera (*Orthoraphium*, *Psammochloa*, *Ptilagrostiella*, *Thorneochloa*, *Timouria*, *Trikeraia*), the enormous significance of whole-genome duplications is clearly obvious in Stipeae. This was pointed out already by Tzvelev (1977), who noted that most extant Stipeae are polyploids and have hybrid origin as corroborated by our exemplary findings on the single-copy gene *Acc1* in *Austrostipa* and *Stipa* (Fig. 3). Although no correlation between whole-genome duplication and diversification could be found in many tested clades of angiosperms (Clark & Donoghue, 2018), all of the larger, speciose genera of Stipeae have consistently polyploid species as far as known, for example, *Stipa* (>150 species) as delineated in the present (*Stipa* s.str.), *Nassella* (117), *Austrostipa* (64), *Eriocoma* (27), *Pappostipa* (31), etc. *Piptatherum* (32 species) and *Ptilagrostis* (15) are the largest genera with only diploids known (Appendix 2). Only two further genera have only diploids, namely *Ortachne* with two and *Oloptum* with a single species. *Piptochaetium* (35) and *Achnatherum* (21) have prevailingly diploids but also polyploids (Appendix 2).

### Biogeographic relations and origin of *Austrostipa* and *Anemanthele*

*Austrostipa* and *Anemanthele* were previously considered as overall rather derived members of Stipeae (ITS analyses of Jacobs & al., 2000, 2007), having groups of American Stipeae like *Amelichloa*, *Jarava*, *Nassella* and American *Achnatherum*, whose species have meanwhile been transferred to *Eriocoma* and *Pseudoeriocoma* (Peterson & al., 2019) as phylogenetically close relatives. On the other hand, considerable differences in lemma epidermal cell patterns between these genera were noted, for example, the occurrence of specialized elongated fundamental cells with thin, slightly sinuate sidewalls in *Austrostipa* (Barkworth & Everett, 1987; Jacobs & Everett, 1996), whereas *Jarava* and *Nassella* have characteristically shortened fundamental cells (Barkworth & Everett, 1987). Considering *Austrostipa*, Tzvelev (1977) discussed a migration of Stipeae from South America via Antarctica as more likely than migration of Stipeae from north to south from Eurasia to Australia. Nevertheless, he argued that the Australian feathergrasses were morphologically closer to Eurasian sections of *Stipa* than to sections of South American feathergrasses and cited among the examples also section *Stipelulla* which encompassed only *Stipa capensis* Thunb. (≡ *Stipellula capensis*) in his view (Tzvelev, 2011). Within the ‘Transcontinental Stipeae clade’, *Austrostipa* and *Anemanthele* are closely affiliated with Eurasian to Mediterranean genera, namely *Achnatherum* (including few eastern to South Africa species; see above), *Celtica*, *Oloptum* and also *Stipellula*, whereas they are distant to the American members of this clade (Fig. 7). This implies that the ancestors of *Austrostipa* and *Anemanthele* came from the lineage with *x* = 12 (Clade A) and not the mainly American lineage with *x* = 11 (Clade B), and therefore acquainted *x* = 11 in parallel to the American representatives of the ‘Transcontinental Stipeae clade’. Secondly, the ancestor was most likely diploid as most if not all extant species of *Achnatherum* except for South African *A. dregeanum*, comb. nov., with 2*n* = 48 (Appendix 2), and whole-genome duplication followed later but probably preceded the evolutionary radiation of *Austrostipa*. Finally, the colonization of Australia started likely from Central to eastern Asia, where *Achnatherum* species, for example, are well represented in the present and the precursor of *Austrostipa* might have occurred as well. Judging from the current distribution, with no occurrences of *Achnatherum* in subtropical and tropical southeastern mainland Asia and Indonesia/New Guinea, long-distance dispersal from Central to eastern Asia to Australia seems plausible. It would be not without parallel considering, for example, the sister relation between *Duthiea*, a genus of China and the Himalayas, and *Anisopogon* from southeastern Australia in the neighboring tribe Duthieeae, as supported by plastid DNA data (Schneider & al., 2011). *Anemanthele* could have reached New Zealand from Australia, and acquainted and established its main apomorphic character, the occurrence of only a single stamen per floret, due to isolation and initial small population size.

This scenario as pictured in Fig. 8, is by no means warranted because, as noted by Jacobs & Everett (1996: 582), *Anemanthele* bears resemblance with *Achnatherum* rather than with *Austrostipa* and it is not sure that the precursor of *Anemanthele* came from Australia and had something to do with *Austrostipa*. *Achnatherum* is not represented in Australia by any autochthonous species, whereas, New Zealand harbors an endemic species of *Achnatherum*, *A. petriei* (Fig. 8) (Edgar & Connor, 2000), the presumably only autochthonous representative of this genus in whole Australasia. Chromosome counts reported 2*n* = 42 for this species (Appendix 2), which is remarkable because otherwise 2*n* = 24 is characteristic of *Achnatherum* (Appendix 2), seemingly except for *A. bromoides*, in which 2*n* = 28 was repeatedly reported (see above). The hexaploid number of *A. petriei* is most likely based on *x* = 11, if aneusomatic change from 44 to 42 is assumed, rather than on *x* = 12. *Achnatherum petriei* thus seems to share with *Austrostipa* and *Anemanthele* the apomorphic basic chromosome number of *x* = 11 in the ‘Transcontinental Stipeae clade’ and has the same ploidy level of 4*x*, both in marked contrast to typical *Achnatherum*. It is reasonable to assume therefore that *Achnatherum petriei* might be a close relative of *Anemanthele* and *Austrostipa* pending further investigation.

**Fig. 8.**
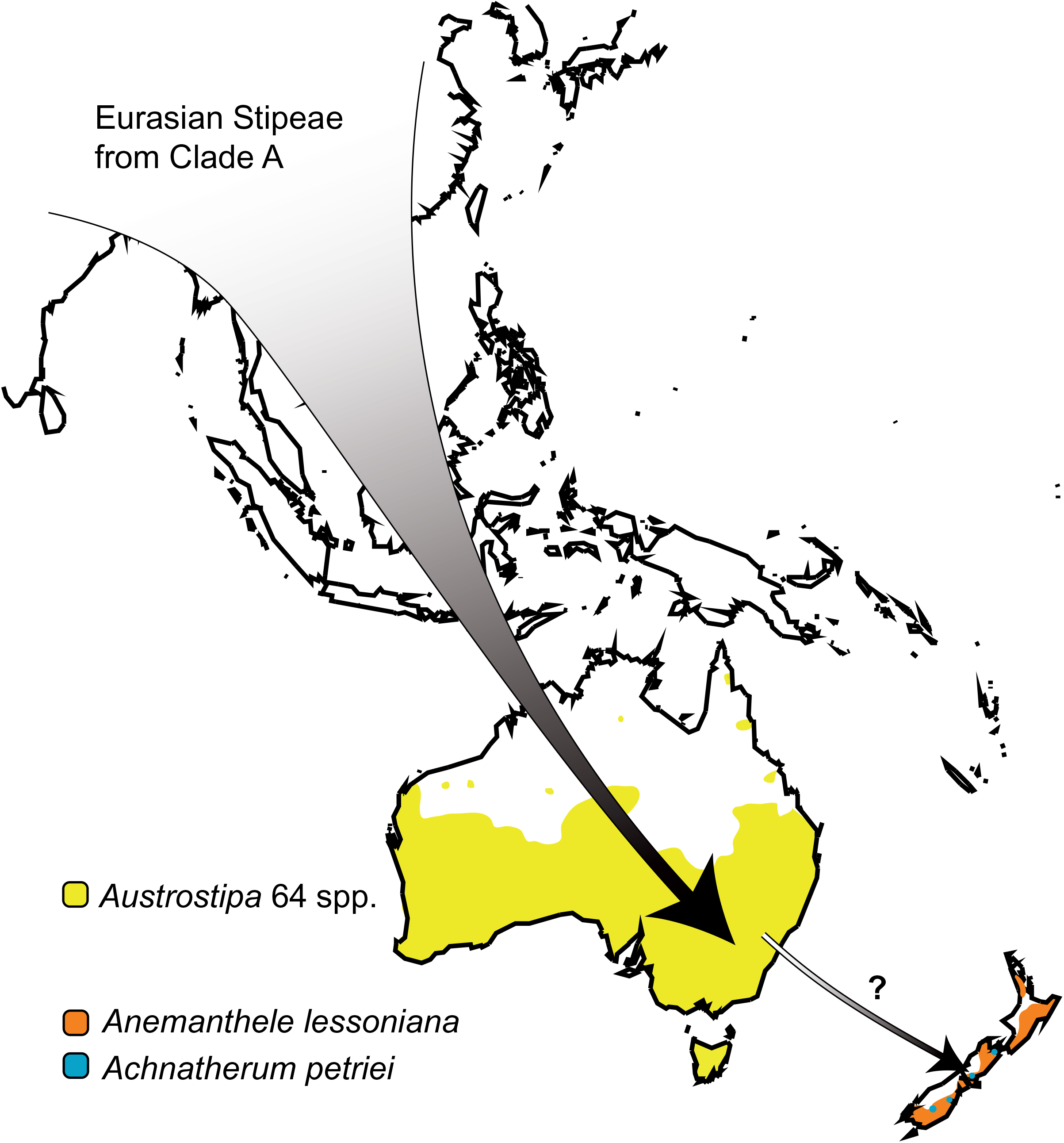
Simplified distributions ranges of *Austrostipa* (ca. 64 species in Australia and Tasmania), monogeneric *Anemanthele* (*A. lessoniana*; endemic New Zealand) and *Achnatherum petriei* (endemic New Zealand), the presumably sole autochthonous representative of this genus in Australasia. The large arrow points to a hypothesized colonization of Australia by Central to east Asian ancestors via long-distance dispersal. The small arrow points to a possible colonization of New Zealand from Australia rather than directly from Asia. See text for discussion of phylogenetic relationships of the three genera and their distribution.

## NEW NAMES AND COMBINATIONS

***Achnatherum dregeanum*** Röser, Tkach & M.Nobis, **comb. nov.** ≡ *Stipa dregeana* Steud., Syn. Pl. Glumac. 1(2): 132. 1854.

***Austrostipa*** subg. ***Paucispiculatae*** Röser, Tkach & M.Nobis, **subg. nov.** – Type: *A. muelleri* (Tate) S.W.L.Jacobs & J.Everett

### Description

Inflorescence reduced to 1‒3 spikelets, lemma with maize-like epidermal pattern, with short fundamental cells and frequent ovate silica bodies, apex of lemma with two distinct, ca. 3 mm long lobes; spreading plants without basal tuft of leaves.

### Included species

*A. muelleri*.

### Distribution

SE South Australia and S Victoria.

### Etymology

The epithet is derived from Latin ‘paucus’ (few) and ‘spicula, spiculatus’ (spikelet, with spikelets).

## AUTHORS CONTRIBUTION

Research conception and design: MEB, MR, NT, SWLJ, MN; field work: MEB, SWLJ; acquisition of molecular data and phylogenetic analysis: JS, NT, HB; acquisition and analysis of micromorphological and morphological data, scanning electron microscopy (SEM): MN; acquisition and analysis of cytogenetic data: MR, GW, NT; drafting of the manuscript: MR, NT, MN; final critical revision: all authors. — NT, https://orcid.org/0000-0002-4627-0706; MN, https://orcid.org/0000-0002-1594-2418; HB, https://orcid.org/0000-0003-3700-2942; GW, https://orcid.org/0000-0002-9866-335X; MEB, https://orcid.org/0000-0001-9785-1538; MR, https://orcid.org/0000-0001-5111-0945.

## ACKNOWLEDGEMENTS

We are very grateful to Bärbel Hildebrandt and Laura Freisleben (Halle) for technical support in the molecular laboratory work. We wish to thank the Millennium Seedbank, Wakehurst Place, Royal Botanic Gardens, Kew and the herbaria listed in Appendix 1 for providing plant material to our study. Dale Dixon (Sydney) provided voucher information for some accessions of NSW. A research grant of the German Research Foundation to MR (DFG Ro 865/8) is gratefully acknowledged.

## Supplementary Figures

**Fig. S1.**
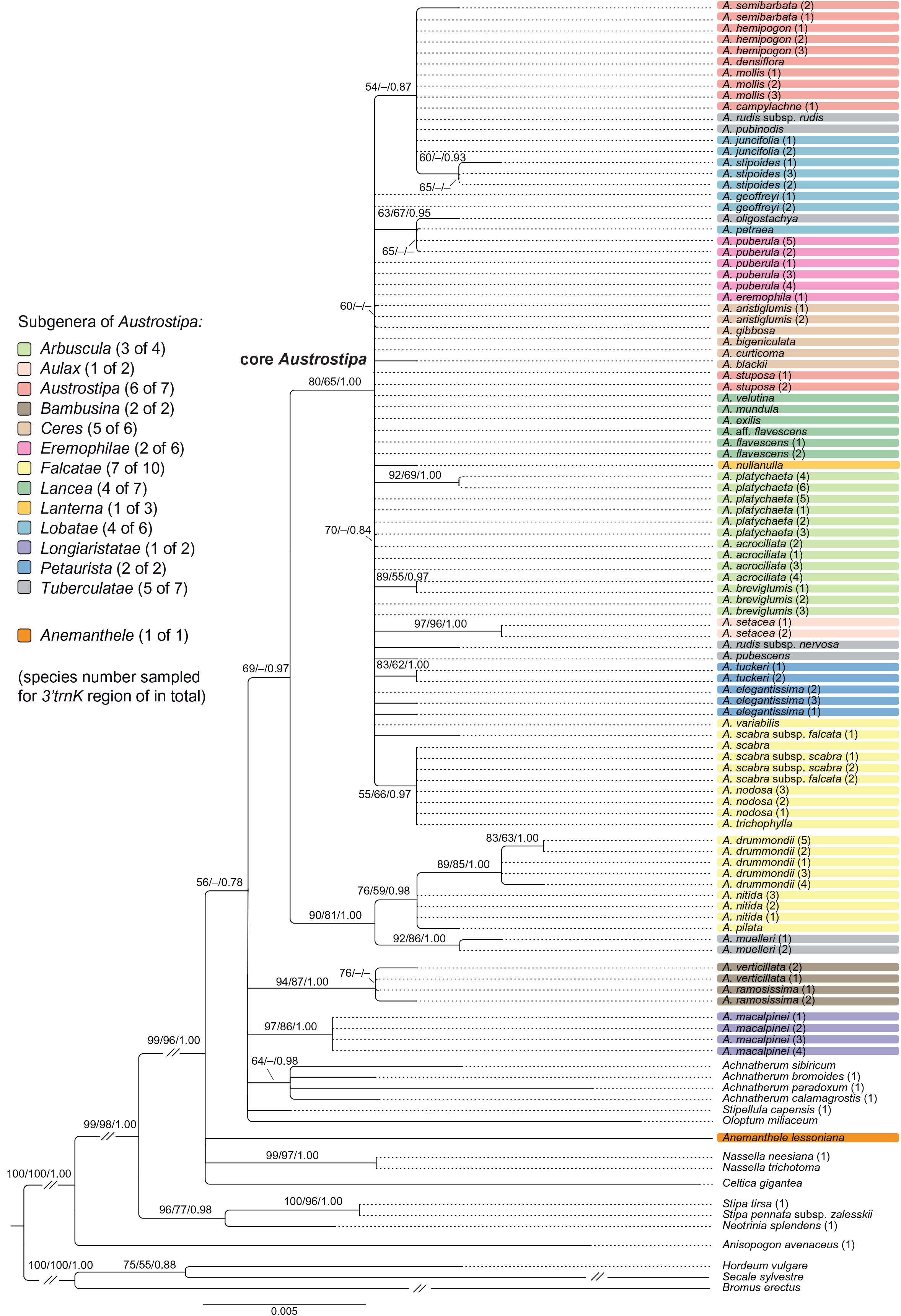
Maximum likelihood phylograms of all studied accessions of *Austrostipa* species inferred from plastid 3’*trnK* region DNA sequences with *Anisopogon avenaceus* (Duthieeae), *Bromus erectus* (Bromeae), *Hordeum vulgare* and *Secale sylvestre* (both Triticeae) used as outgroup. ML and MP bootstrap support values ≥50% as well as Bayesian PP ≥0.5 are indicated on the branches. Clades with ML support <50% are collapsed. The subgenera of *Austrostipa* are marked by different colors. Abbreviation: *A*. = *Austrostipa*.

**Fig. S2.**
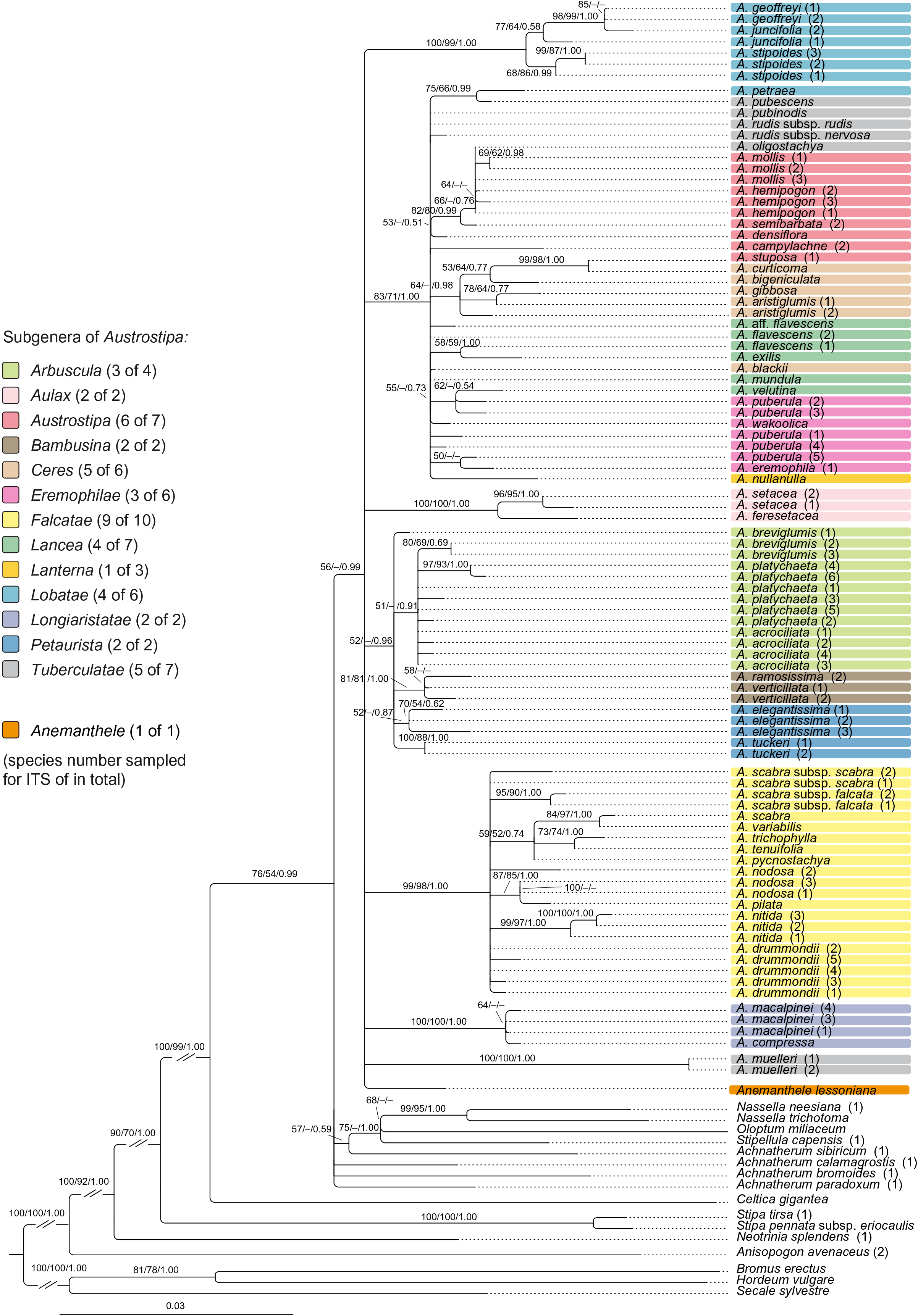
Maximum likelihood phylograms of all studied accessions of *Austrostipa* species inferred from nuclear ribosomal ITS1–5.8S gene–ITS2 DNA sequences with *Anisopogon avenaceus* (Duthieeae), *Bromus erectus* (Bromeae), *Hordeum vulgare* and *Secale sylvestre* (both Triticeae) used as outgroup. ML and MP bootstrap support values ≥50% as well as Bayesian PP ≥0.5 are indicated on the branches. Clades with ML support <50% are collapsed. The subgenera of *Austrostipa* are marked by different colors. Abbreviation: *A*. = *Austrostipa*.

**Fig. S3.**
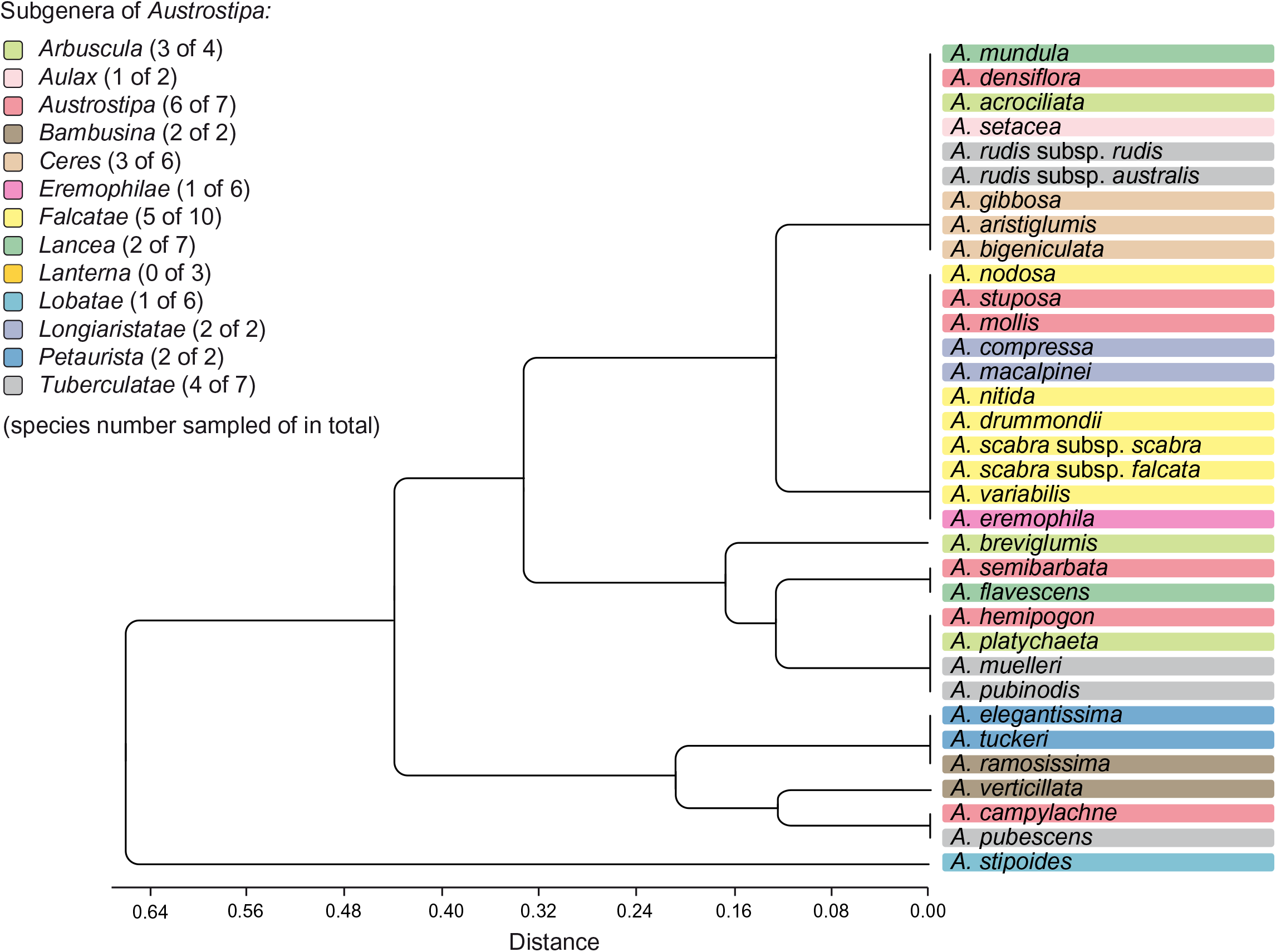
Cluster analysis (UPGMA) performed on four micromorphological characters of 34 *Austrostipa* taxa. See suppl. Table S2 for the data matrix evaluated. The subgenera of *Austrostipa* are marked by different colors. Abbreviation: *A*. = *Austrostipa*.

**Fig. S4.**
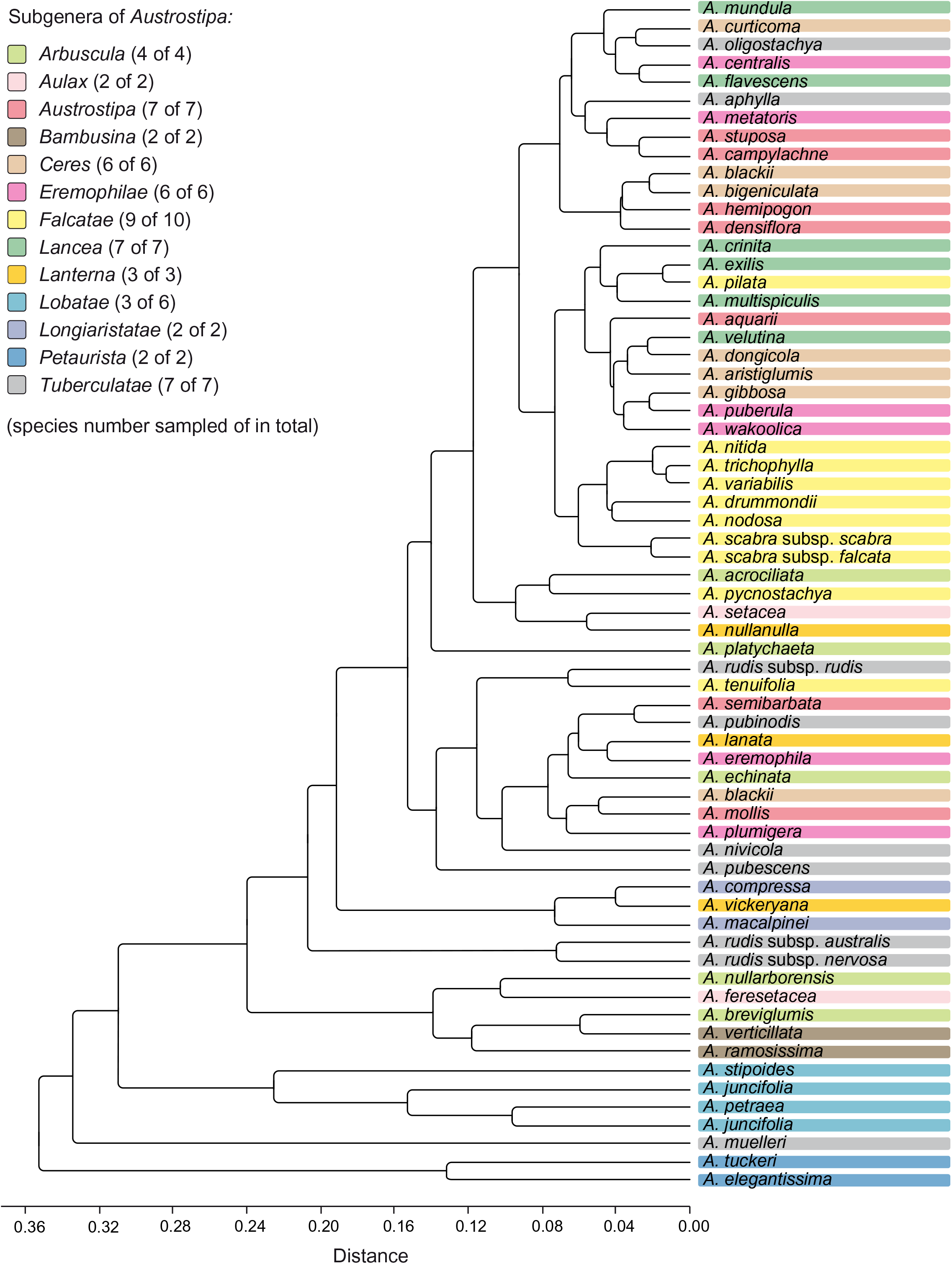
Cluster analysis (UPGMA) performed on nine macromorphological characters of 65 *Austrostipa* taxa. See suppl. Table S1 for the data matrix evaluated. The subgenera of *Austrostipa* are marked by different colors. Abbreviation: *A*. = *Austrostipa*.

**Supplementary Appendices S1, S2, S3**. FASTA files sequence alignments *matK*, ITS, *Acc1*.

## Appendix 1 Taxa studied for DNA sequences, morphology and lemma micromorphology

Taxon, geographical origin, voucher information with collectors and herbarium code and ENA/GenBank accession numbers for plastid matK gene–3′*trnK* exon region; nuclear ribosomal ITS1–5.8S gene–ITS2 and nuclear single-copy gene Acc1. Sequences #####–###### were newly generated for this study. A single asterisk (*) indicates sequences previously generated in our lab, two asterisks (**) indicate sequences taken from ENA/GenBank, a dash (–) missing data. BG = Botanical Garden; LEP = lemma epidermal pattern studied; LEP ill. = LEP illustrated in Figs. 4, 5; morph = morphological data for suppl. Table S2 taken from Everett & al. (2009); MSB = Millennium Seed Bank, Kew, Wakehurst Place; NSW = New South Wales; NT = Northern Territory; SA = South Australia; TAS = Tasmania; VIC = Victoria, WA = Western Australia.

***Achnatherum bromoides*** (L.) P.Beauv.: **(1)** FN434250*; FN434517*; ‒; **(2)** Iran, *T. Alexeenko* 316 (LE); LEP ill. ***A. calamagrostis*** (L.) P.Beauv.: **(1)** FN434255*; FN434520*; ‒; **(2**) Hungary, *H. Petry s.n.* (KRA); LEP ill. ***A. paradoxum*** (L.) Banfi, Galasso & Bartolucci: **(1)** grown in BG Halle, *M. Röser 11092*, 30 May 2012 (HAL); #matK#; #ITS#; #Acc1#; **(2)** France, between Garabella and Montpellier, *J. Kornaś s.n.* (KRA); LEP ill. ***A. sibiricum*** (L.) Keng ex Tzvelev: Mongolia, Töv ajmag SE Ulaanbaatar; *H. Heklau s.n.,* 2006. Grown from seed in BG Halle no. R503 (no voucher); FN434339*; FN434595*; #Acc1#. ***Anemanthele lessoniana*** (Steud.) Veldkamp: Grown from seed obtained in 2007 from Royal Horticultural Society Garden, Wisley, U.K. in BG Halle no. R610 (no voucher); LN832437*; LN832437*; #Acc1#. ***Anisopogon avenaceus*** R.Br.: AM234575*; FM179386*; ‒. ***Austrostipa acrociliata*** (Reader) S.W.L.Jacobs & J.Everett: **(1)** Australia, SA, 33°51’50.19”S, 136°8’55.88”E, *T.S. Te & P.J. Lang s.n.*, 26 Nov. 2007, collected seed at MSB no. 426442. Plants grown from seed at BG Halle by G. Winterfeld GW92 (HAL0137631); LN832440*; LN832412*; ‒; LEP; **(2)** Australia, VIC, 34°50’26”S, 142°17’53”E, *D. Roberts, N.G. Walsh & M.J. Hirst s.n.*, 23 Oct. 2007, collected seed at MSB no. 460352 (K). Plants grown from seed at BG Halle by G. Winterfeld GW97 (HAL0141317); LN832441*; LN832413*; #Acc1#; LEP; **(3)** Australia, SA, Eyre Peninsula, c. 54 km SE of Streaky Bay, Port Kenny road, 33°7’11”S, 134°37’9”E, *S.W.L. Jacobs 9760 & M.E. Barkworth*, 06 Dec. 2007 (AD229036, NSW832532); #matK#; #ITS#; ‒; **(4)** Australia, WA, Eyre, Hopetoun, 33°56’44”S, 120°07’34”E, *S.W.L. Jacobs 9746 & M.E. Barkworth*, 02 Dec. 2007 (HAL0110716, duplicates: NSW832487, PERTH8048193, UTC00252175); #matK#; #ITS#; ‒; LEP. ***A. aphylla*** (Rodway) S.W.L.Jacobs & J.Everett: morph. ***A. aquarii*** (Vickery, S.W.L.Jacobs & J.Everett) S.W.L.Jacobs & J.Everett: morph. ***A. aristiglumis*** (F.Muell.) S.W.L.Jacobs & J.Everett **(1)** Australia, VIC, Murray Mallee, c. 11 km S of Hopetoun, Warracknabeal road, 35°48’45”S, 142°25’22”E, *S.W.L. Jacobs 9777 & M.E. Barkworth*, 09 Dec. 2007 (HAL0110797, duplicates: B, BA, K, MEL2330408A, NSW594293, TCD, US, UTC); #matK#; #ITS#; ‒; LEP ill.; **(2)** Australia, NSW, South Western Plains, c. 1 km S of Darlington Point, 34°35’04”S, 145°59’40”E, *S.W.L. Jacobs 9795 & M.E. Barkworth*, 12 Dec. 2007 (HAL0107516, duplicates: BA, NSW594352, UTC); #matK#; #ITS#; ‒; LEP. ***A. bigeniculata*** (Hughes) S.W.L.Jacobs & J.Everett: Australia, NSW, Southern Tablelands, c. 17 km E of Bungonia, Goulburn road, 34°48’45”S, 149°47’03”E, *S.W.L. Jacobs 9728 & M.E. Barkworth*, 23 Nov. 2007 (HAL0110805, duplicates: B, BA, K, NSW832478, TCD, UTC00252175); #matK#; #ITS#; ‒; LEP ill. ***A. blackii*** (C.E.Hubb.) S.W.L.Jacobs & J.Everett: Australia, SA, *D.J. & M.K. Jones s.n.*, 22 Nov. 2005, collected seed at MSB no. 297934 (K). Plants grown from seed at BG Halle by G. Winterfeld GW53 (HAL0140625, HAL0137622); LN832453*; LN832421*; #Acc1#. ***A. breviglumis*** (J.M.Black) S.W.L.Jacobs & J.Everett: **(1)** Australia, SA, 34°52’30.4”S, 138°42’49.46”E, 20 Dec. 2005, collected seed at MSB no. 302966 (K). Plants grown from seed at BG Halle by G. Winterfeld GW56 (HAL0141315, HAL0137637); LN832442*; LN832414*; #Acc1#; **(2)** Australia, SA, Eyre Peninsula, c. 135 km SE of Elliston, Port Lincoln rd, 34°35’04”S, 135°35’06”E, *S.W.L. Jacobs 9766 & M.E. Barkworth*, 06 Dec. 2007 (HAL0110713, duplicates: AD228918, UTC00252101, NSW832490); #matK#; #ITS#; ‒; LEP ill.; **(3)** same collection as (2); #matK#; #ITS#; ‒. ***A. campylachne*** (Nees) S.W.L.Jacobs & J.Everett**: (1)** Australia, WA, 12.5 km E of Catchment Road on Qualen Road, 32°5’16.7”S, 116°40’14.2”E, *A. Crawford ADC 987*, 26 Nov. 2005, collected seed at MSB no. 376226 (K, PERTH7346883). Plants grown from seed at BG Halle by G. Winterfeld GW89 (HAL0137630); LN832445*; ‒; #Acc1#; **(2)** ‒; GU254627; ‒; **(3)** Australia, WA, Smith’s Mill, *A. Morrison s.n.*, 13 Nov. 1897 (E690354); LEP ill. ***A. centralis*** (Vickery, S.W.L.Jacobs & J.Everett) S.W.L.Jacobs & J.Everett: morph. ***A. compressa*** (R.Br.) S.W.L.Jacobs & J.Everett: **(1)** ‒; EU489100**; ‒; **(2)** Australia, WA, Cottesloe, *A. Morrison s.n.*, 21.10.1899 (E690358); LEP ill. ***A. crinita*** (Gaudich.) S.W.L.Jacobs & J.Everett: morph. ***A. curticoma*** (Vickery) S.W.L.Jacobs & J.Everett: LN832454*; LN832422*; ‒. ***A. densiflora*** (Hughes) S.W.L.Jacobs & J.Everett: Australia, NSW, Southern Tablelands, Bungonia turnoff, S of Marulan, Hume Highway, 34°43’59”S, 149°58’47”E, *S.W.L. Jacobs 9725*, 23 Nov. 2007 (HAL0110782, duplicates: NSW832480, UTC00252106); #matK#; #ITS#; ‒; LEP ill. ***A. dongicola*** (Vickery, S.W.L.Jacobs & J.Everett) S.W.L.Jacobs & J.Everett: morph. ***A. drummondii*** (Steud.) S.W.L.Jacobs & J.Everett: **(1)** LN832455*; LN832423*; ‒; **(2)** Australia, WA, Dundas (S) Eucla, base of Eucla Pass, 31°40’52”S, 128°52’30”E., *S.W.L. Jacobs 9752 & M.E. Barkworth*, 04 Dec. 2007 (NSW832530, PERTH8500231); #matK#; #ITS#; ‒; **(3)** Australia, SA, South-eastern, c. 28 km E of Woods Well, Tintinara road, 35°58’25”S, 139°50’25”E, *S.W.L. Jacobs 9776 & M.E. Barkworth*, 09 Dec. 2007 (HAL0110798, duplicates: B, BA, MEL2330245A, NSW594288, TCD, UTC); #matK#; #ITS#; ‒; LEP ill.; **(4)** Australia, SA, Nullarbor, c. 76 km E of Border Village, Nullarbor road, 31°36’04”S, 129°46’30”E, *S.W.L. Jacobs 9757 & M.E. Barkworth*, 05 Dec. 2007 (HAL064847, duplicates: AD229038, BA, NSW832524, TCD, UTC); #matK#; #ITS#; ‒; LEP; **(5)** Australia, WA, Eucla, base of Eucla Pass, 31°40’52”S, 128°52’30”E, *S.W.L. Jacobs 9754 & M.E. Barkworth*, 04 Dec. 2007 (HAL0110717, duplicates: NSW594291, PERTH8003912, UTC00252318); #matK#; #ITS#; ‒; LEP. ***A. echinata*** (Vickery, S.W.L.Jacobs & J.Everett) S.W.L.Jacobs & J.Everett: morph. ***A. elegantissima*** (Labill.) S.W.L.Jacobs & Everett: **(1)** Australia, SA, 34°52’26.15”S,138°42’44.73”E, 20 Dec. 2005, collected seed at MSB no. 302977 (K). Plants grown from seed at BG Halle by G. Winterfeld GW57 (HAL0137633); LN832467*; #ITS#; #Acc1#; **(2)** Australia, WA, Roe, c. 68 km S of Norseman, Salmon Gums road, 32°44’31”S, 121°32’08”, *S.W.L. Jacobs 9736 & M.E. Barkworth*, 01 Dec. 2007 (HAL0111057, duplicates: NSW832555, PERTH8256861, UTC); #matK#; #ITS#; ‒; LEP; **(3)** Australia, WA, Eyre, c. 50 km E of Ravensthorpe, Esperance Road, 33°40’17”S, 120°39’34”E, *S.W.L. Jacobs 9749 & M.E. Barkworth*, 03 Dec. 2007 (HAL0110796, duplicates: NSW594284, PERTH8003912, TCD, UTC00252324); #matK#; #ITS#; ‒; LEP ill.; **(4)** Australia, SA, Murray Mallee, between Kingston and Waikerie, *D.J.E. Whibley 2651*, 28 Sep. 1968 (E690373); LEP. ***A. eremophila*** (Reader) S.W.L.Jacobs & J.Everett: **(1)** Australia, WA, Eyre, c. 4 km N of Esperance, Ravensthorpe road, 33°50’8”S, 121°51’40”E, *S.W.L. Jacobs 9739 & M.E. Barkworth*, 01 Dec. 2007 (NSW832514, PERTH8003882, UTC00252213); #matK#; #ITS#; ‒; **(2)** Australia, SA, Redcliff Survey Area, 33°43’S, 137°51’E, *R.J. Chinnock 1594* (E690379); LEP ill. ***A. exilis*** (Vickery) S.W.L.Jacobs & J.Everett: Australia, WA, Esperance (S), Mondrain Island, off Esperance, NE slope of N part of Island, 34°7’18”S, 122°14’38”E, *J.A. Cochrane 4286*, 20 Nov. 2002, collected seed at MSB no. 359476 (K, PERTH6476562). Plants grown from seed at BG Halle by G. Winterfeld GW84 (HAL0140634, HAL0137636); LN832462*; LN832430*; #Acc1#. ***A. feresetacea*** (Vickery, S.W.L.Jacobs & J.Everett) S.W.L.Jacobs & J.Everett: ‒; JF769061**; ‒. ***A. flavescens*** (Labill.) S.W.L.Jacobs & J.Everett: **(1)** Australia, TAS, 41°24’29.8”S, 148°16’54.5”E, 13 Dec. 2007, collected seed at MSB no. 442387 (K). Plants grown from seed at BG Halle by G. Winterfeld GW98 (no voucher); LN832463*; LN832431*; #Acc1#; **(2)** Australia, SA, South-eastern, c. 18 km S of Rode, Beachport Road, 37°14’38”S, 139°54’22”E, *S.W.L. Jacobs 9684*, 24 Oct. 2007 (HAL0110799, duplicates: AD228901, NSW831786, UTC00252189); #matK#; #ITS#; ‒; LEP ill. ***A.* aff. *flavescens*** (Labill.) S.W.L.Jacobs & J.Everett: Australia, SA, c. 7 km NW Port Lincoln, Elliston Road, along powerline track near road, 34°44’1”S, 135°8’49”E, *S.W.L. Jacobs 9767, 06 Dec. 2007* (NSW860187); #matK#; #ITS#; ‒. ***A. geoffreyi*** S.W.L.Jacobs & J.Everett: **(1)** Australia, WA, Lake Grace (S), 33°5’23”S, 119°33’45”E, *J.A. Cochrane 4243*, 06 Nov. 2002, collected seed at MSB no. 213064 (K, PERTH6476740). Plants grown from seed at BG Halle by G. Winterfeld GW44 (no voucher); LN832465*; LN832433*; #Acc1#; **(2)** Australia, WA, Roe, Lake King, c. 11.5 km W of Lake King township, 33°05’23”S, 119°33’20”E, *S.W.L. Jacobs 9743 & M.E. Barkworth*, 02 Dec. 2007 (HAL0110783, UTC00252284, NSW832518); #matK#; #ITS#; ‒. ***A. gibbosa*** (Vickery) S.W.L.Jacobs & J.Everett: Australia, SA, Northern Lofty, c. 19 km SE of Crystal Brook, Clare road, 33°26’16”S, 138°22’11”E, *S.W.L. Jacobs 9773 & M.E. Barkworth*, 08 Dec. 2007 (HAL0110795, duplicates: AD228915, BA, NSW594286, TCD, UTC00252323); #matK#; #ITS#; ‒; LEP ill. ***A. hemipogon*** (Benth.) S.W.L.Jacobs & J.Everett: **(1)** LN832446*; LN832417*; ‒; **(2)** Australia, WA, Albany (C), Barrs Road Reserve, 34°18’24”S, 118°26’49”E, *J.A. Cochrane 5022*, 01 Nov. 2004, collected seed at MSB no. 359502 (K). Plants grown from seed at BG Halle by G. Winterfeld GW85 (HAL0140623, HAL0137629); LN832447*; LN832418*; #Acc1#; **(3)** Australia, WA, Roe, Lake King, island c. 8 km W of Lake King township, 33°05’26”S, 119°35’32”E, *S.W.L. Jacobs 9744 & M.E. Barkworth*, 02 Dec. 2007 (HAL0110789, duplicates: NSW832519, PERTH8003904, UTC00252265); #matK#; #ITS#; ‒; LEP ill.; **(4)** Australia, WA, Roe, c. 13 km S of Grass patch, Esperance Road, 33°20’48”S, 121°42’48”E, *S.W.L. Jacobs 9737a & M.E. Barkworth*, 01 Dec. 2007 (HAL0110788, duplicates: NSW832517, PERTH5038308, UTC00252282, UTC); LEP. ***A. juncifolia*** (Hughes) S.W.L.Jacobs & J.Everett: **(1)** Australia, SA, Eyre, 58 km S of Grass Patch, Esperance road, 33°42’31”S, 121°50’21”E, *S.W.L. Jacobs 9738 & M.E. Barkworth*, 01 Dec. 2007 (HAL0111022, duplicates: NSW832548, PERTH5038308, UTC00259369); #matK#; #ITS#; ‒; **(2)** Australia, WA, Roe, Lake King, Western shore. W of Lake King township, 33°05’24”S, 119°32’42”E, *S.W.L. Jacobs 9742 & M.E. Barkworth*, 02 Dec. 2007 (HAL0107543, duplicates: NSW832527, UTC); #matK#; #ITS#; ‒. ***A. lanata*** (Vickery, S.W.L.Jacobs & J.Everett) S.W.L.Jacobs & J.Everett: morph. ***A. macalpinei*** (Reader) S.W.L.Jacobs & Everett: **(1)** Australia, WA, 31°2’22”S, 116°2’37”E, *L. Lumbus s.n.*, 12 Nov. 2003, collected seed at MSB no. 230847 (K). Plants grown from seed at BG Halle by G. Winterfeld GW47 (no voucher); #matK#; #ITS#; #Acc1#; **(2)** Australia, WA, Eyre, c. 70 km E of Ravensthorpe, Esperance Road, 33°40’47”S, 120°45’38”E, *S.W.L. Jacobs 9740 & M.E. Barkworth*, 01 Dec. 2007 (HAL0110785, duplicates: NSW832512, PERTH8003955, UTC00252316, UTC); #matK#; ‒; ‒; LEP ill.; **(3)** Australia, WA, Eyre, c. 40 km E of Ravensthorpe, Esperance road, 33°38’53”S, 120°26’58”E, *S.W.L. Jacobs 9748 & M.E. Barkworth*, 03 Dec. 2007 (NSW832528, PERTH8500223); #matK#; #ITS#; ‒; **(4)** same collection as (2); #matK#; #ITS#; ‒; LEP. ***A. metatoris*** (J.Everett & S.W.L.Jacobs) S.W.L.Jacobs & J.Everett: morph. ***A. mollis*** (R.Br.) S.W.L.Jacobs & J.Everett: **(1)** Australia, SA, 35°10’0.09”S, 138°50’44.27”E, *D.J. Duval s.n.*, 26 Nov. 2006, collected seed at MSB no. 359904 (K). Plants grown from seed at BG Halle by G. Winterfeld GW79 (HAL0141318, HAL0137620); LN832448*; LN832419*; #Acc1#; **(2)** Australia, VIC, Grampians, Rose Creek road, c. 55 km SE of Horsham. Grampians National Park, 37°07’33”S, 142°24’46”E, *S.W.L. Jacobs 9787 & M.E. Barkworth*, 11 Dec. 2007 (HAL0107541, duplicates: MEL2340051A, NSW594349, UTC); #matK#; #ITS#; ‒; LEP ill.; **(3)** Australia, SA, South-eastern, c. 18 km S of Pinaroo, Bordertown Road, 35°23’44”S, 140°49’14”E, *S.W.L. Jacobs 9701*, 29 Oct. 2007 (HAL0110800, duplicates: AD228900, BA, NSW831784, UTC00252178); #matK#; #ITS#; ‒; LEP. ***A. muelleri*** (Tate) S.W.L.Jacobs & Everett: **(1)** Australia, VIC, Grampians, Harrops Track, c. 1 km N of Grahams Creek (c. 9 km N Glenelg River Road.), 37°18’14”S, 142°15’04”E, *S.W.L. Jacobs 9790 & M.E. Barkworth*, 11 Dec. 2007 (HAL0110712, duplicates: BA, MEL2330272A, NSW832495, UTC00252111); #matK#; #ITS#; ‒; LEP ill.; **(2)** Australia, VIC, Grampians, Grampians National Park, junction of Glenelg River Road and Harrop Track, 37°22’7”S, 142°11’56”E, *S.W.L. Jacobs 9791 & M.E. Barkworth*, 11 Dec. 2007 (MEL2340048A, NSW594311); #matK#; #ITS#; ‒. ***A. multispiculis*** (J.M.Black) S.W.L.Jacobs & J.Everett: morph. ***A. mundula*** (J.M.Black) S.W.L.Jacobs & J.Everett: Australia, SA, Eyre Peninsula, Tooligie, c. 45 km N of Cummins, 33°51’42”S, 135°42’20”E, *S.W.L. Jacobs 9768 & M.E. Barkworth*, 06 Dec. 2007 (HAL0110714, duplicates: AD228922, NSW832492, UTC00252183); #matK#; #ITS#; ‒; LEP. ***A. nitida*** (Summerh. & C.E.Hubb.) S.W.L.Jacobs & J.Everett: **(1)** Australia, NSW, South Western Plains, c. 40 km N of Goolgowi, Hillston road, 33°39’43”S, 145°32’33”E, *S.W.L. Jacobs 9803 & M.E. Barkworth*, 12 Dec. 2007 (HAL0107550, duplicates: B, BA, G-DC, NSW594317, TCD, US, UTC); #matK#; #ITS#; ‒; LEP; **(2)** Australia, SA, Eyre Peninsula, c. 40 km E of Kimba, Port Augusta road, 32°55’53”S, 136°55’34”E, *S.W.L. Jacobs 9769 & M.E. Barkworth*, 07 Dec. 2007 (HAL0111023, duplicates: AD238077, B, G, K, NSW832547, TCD, UTC); #matK#; #ITS#; ‒; LEP ill.; **(3)** Australia, WA, Roe, c. 40 km S of Norseman, Salmon Gums road, 32°19’09”S, 121°45’33”E, *S.W.L. Jacobs 9731 & M.E. Barkworth*, 01 Dec. 2007 (HAL0111024, duplicates: NSW832550, PERTH8256837, UTC); #matK#; #ITS#; ‒; LEP. ***A. nivicola*** (J.H.Willis) S.W.L.Jacobs & J.Everett: morph. ***A. nodosa*** (S.T.Blake) S.W.L.Jacobs & J.Everett: **(1)** Australia, VIC, Wimmera, c. 23 km SW of Warracknabeal, Dimboola Road, 36°22’05”S, 142°10’55”E, *S.W.L. Jacobs 9780 & M.E. Barkworth*, 10 Dec. 2007 (HAL0110711, duplicates: B, BA, G, K, MEL2331037A, NSW832494, UTC00252262); #matK#; #ITS#; ‒; LEP ill.; **(2)** Australia, SA, Eyre Peninsula, c. 3 km SE of Elliston, Port Lincoln road, 33°39’37”S, 134°54’53”E, *S.W.L. Jacobs 9762 & M.E. Barkworth*, 06 Dec. 2007 (HAL0110784, duplicates: AD228913, BA, NSW832520, UTC00252263); #matK#; #ITS#; ‒; LEP; **(3)** LN832456*; LN832424*; ‒. ***A. nullanulla*** (J.Everett & S.W.L.Jacobs) S.W.L.Jacobs & J.Everett: Australia, SA, 31°39’47.25”S, 135°15’22.34”E, *D.J. Duval, B. Haas & P. Swanson s.n.*, 09 Nov. 2005, collected seed at MSB no. 291460 (K). Plants grown from seed at BG Halle by G. Winterfeld GW66 (no voucher); LN832464*; LN832432*; #Acc1#. ***A. nullarborensis*** (Vickery, S.W.L.Jacobs & J.Everett) S.W.L.Jacobs & J.Everett: morph. ***A. oligostachya*** (Hughes) S.W.L.Jacobs & J.Everett: Australia, SA, 35°35’24.29”S, 138°33’29.35”E, *T.S. Te*, *D.J. Duval & M.J. Thorpe s.n.*, 07 Nov. 2006, collected seed at MSB no. 359889 (K). Plants grown from seed at BG Halle by G. Winterfeld GW78 (HAL0141319, HAL0141389, HAL0137624); LN832468*; LN832436*; #Acc1#. ***A. petraea*** (Vickery) S.W.L.Jacobs & J.Everett: Australia, SA, Flinders Ranges National Park, Bunyeroo Gorge, South Flinders Ranges, 31°25’9”S, 138°33’48”E., *S.W.L. Jacobs 9698*, 28 Oct. 2007 (NSW831779); #matK#; #ITS#; #Acc1#. ***A. pilata*** (S.W.L.Jacobs & J.Everett) S.W.L.Jacobs & J.Everett: LN832461*; LN832429*; ‒. ***A. platychaeta*** (Hughes) S.W.L.Jacobs & J.Everett: **(1)** Australia, SA, Murray, c. 3 km N of Cambrai, Sedan Road, 34°37’39”S, 139°16’55”E, *S.W.L. Jacobs 9700*, 29 Oct. 2007 (HAL0110801, duplicates: AD229030, NSW831785, UTC00252179); #matK#; #ITS#; ‒; LEP; **(2)** Australia, WA, Coolgardie, c. 47 km S of Norseman, Salmon Gums road, 32°34’2”S, 121°35’27”E, *S.W.L. Jacobs 9735 & M.E. Barkworth*, 01 Dec. 2007 (NSW832549, PERTH8256845); #matK#; #ITS#; ‒; **(3)** Australia, VIC, Wimmera, c. 21 km SW of Warracknabeal, Dimboola road, 36°21’23”S, 142°11’56”E, *S.W.L. Jacobs 9778 & M.E. Barkworth*, 10 Dec. 2007 (HAL0110787, duplicates: B, MEL2330210A, NSW832516, TCD, UTC00252268); #matK#; #ITS#; ‒; LEP; **(4)** Australia, SA, Nullarbor, c. 10 km N of Nullarbor Roadhouse, Murrawinje Cave Road, 31°21’60”S, 130°52’17”E, *S.W.L. Jacobs 9759 & M.E. Barkworth*, 05 Dec. 2007 (NSW832531); #matK#; #ITS#; ‒; **(5)** Australia, WA, Eucla, c. 2 km E of Eucla, Border Village, 31°40’15”S, 128°54’24”E, *S.W.L. Jacobs 9755 & M.E. Barkworth*, 04 Dec. 2007 (HAL0110718, duplicates: BA, NSW594292, PERTH8003920, UTC00252319); #matK#; #ITS#; ‒; LEP ill.; **(6)** Australia, WA, Roe, c. 4 km S of Norseman, Salmon Gums road, 32°19’09”S, 121°45’33”E, *S.W.L. Jacobs 9730 & M.E. Barkworth*, 01 Dec. 2007 (HAL0110801, duplicates: AD, NSW832551, PERTH8256918, UTC); #matK#; #ITS#; ‒; LEP. ***A. plumigera*** (Hughes) S.W.L.Jacobs & J.Everett; morph. ***A. puberula*** (Steud.) S.W.L.Jacobs & J.Everett: **(1)** Australia, WA, Coolgardie, c. 47 km S of Norseman, Salmon Gums rd, 32°34’02”S, 121°35’28”E, *S.W.L. Jacobs 9734 & M.E. Barkworth*, 01 Dec. 2007 (HAL0110709, duplicates: BA, NSW832491, PERTH8048177, UTC00252184); #matK#; #ITS#; ‒; **(2)** Australia, VIC, Wimmera, c. 21 km SW of Warracknabeal, Dimboola road, 36°21’23”S, 142°11’56”E, *S.W.L. Jacobs 9779*, 10 Dec. 2007 (HAL0110786, duplicates: BA, MEL2330238A, NSW832515, TCD, UTC00252266); #matK#; #ITS#; ‒; **(3)** Australia, SA, Nullarbor, c. 42 km E of Border Village, Nullarbor road, 31°37’57”S, 129°25’47”E, *S.W.L. Jacobs 9756 & M.E. Barkworth*, 05 Dec. 2007 (HAL0107544, duplicates: AD229037, BA, NSW832525, UTC); #matK#; #ITS#; #Acc1#; **(4)** Australia, SA, South-eastern, c. 6 km E of Woods Well, Tintinara Road, 35°59’11”S, 139°36’22”E, *S.W.L. Jacobs 9775 & M.E. Barkworth*, 09 Dec. 2007 (HAL0110710, duplicates: AD228898, B, BA, CAN, K, NSW832493, PE, TCD, US, UTC00252185); #matK#; #ITS#; ‒; **(5)** Australia, SA, Eyre Peninsula, c. 15 km SE of Port Kenny, Elliston road, 33°16’19”S, 134°47’15”E, *S.W.L. Jacobs 9761 & M.E. Barkworth*, 06 Dec. 2007 (HAL0111026, duplicates: AD238076, B, BA, G, GRA, K, NSW832554, TCD, UTC); #matK#; #ITS#; ‒. ***A. pubescens*** (R.Br.) S.W.L.Jacobs & J.Everett: Australia, NSW, Central Coast, Kentlyn, 34°03’21”S, 150°52’32”E, S*.W.L. Jacobs 9724 & M.E. Barkworth*, 20 Nov. 2007 (HAL0110778, duplicates: NSW832477, UTC00252176); #matK#; #ITS#; ‒; LEP ill. ***A. pubinodis*** (Trin & Rupr.) S.W.L.Jacobs & J.Everett: Australia, NSW, Grampians, Rose Creek road, c. 55 km SE of Horsham, Grampians National Park, 37°07’33”S, 142°24’46”E, *S.W.L. Jacobs 9788 & M.E. Barkworth*, 11 Dec. 2007 (HAL0107511, duplicates: MEL2340073A, NSW594312, UTC); #matK#; #ITS#; ‒; LEP ill. ***A. pycnostachya*** (Benth.) S.W.L.Jacobs & J.Everett; ‒; EU489101**; ‒. ***A. ramosissima*** (Trin.) S.W.L.Jacobs & J.Everett: **(1)** Australia, QLD, 27°28’25.8”S, 149°16’52.4”E, *T. Tyson-Doneley s.n.*, 10 May 2005, collected seed at MSB no. 273060 (K). Plants grown from seed at BG Halle by G. Winterfeld GW65 (HAL0137634); LN832451*; ‒; #Acc1#; **(2)** Australia, NSW, Central Coast, Mount Annan BG, 34°04’17”S, 150°46’03”E, *S.W.L. Jacobs 9831 & M.E. Barkworth*, 08 Jan. 2008 (HAL0107512, duplicates: BA, NSW594356, UTC); #matK#; #ITS#; ‒; LEP ill. ***A. rudis*** (Spreng.) S.W.L.Jacobs & Everett subsp. ***rudis***: Australia, VIC, Gippsland Plain, c. 3 km N of Sale, Bairnsdale Road, 38°04’45”S, 147°04’18”E, *S.W.L. Jacobs 9710*, 31 Oct. 2007 (HAL0110802, duplicates: MEL2330239A, NSW831783, UTC00252181); #matK#; #ITS#; ‒; LEP. ***A. rudis*** subsp. ***australis*** (J.Everett & S.W.L.Jacobs) S.W.L.Jacobs & J.Everett: Australia, VIC, Ringwood, *A. Morrison s.n.*, 21.10.1891 (E692076); LEP ill. ***A. rudis*** subsp. ***nervosa*** (Vickery) S.W.L.Jacobs & J.Everett: Australia, NSW,Central Coast, Georges River, Appin, 34°12’35”S, 150°47’46”E, *S.W.L. Jacobs 9722 & M.E. Barkworth*, 20 Nov. 2007 (HAL0110779, duplicates: BA, NSW832476, UTC00252104); #matK#; #ITS#; ‒; LEP. ***A. scabra*** (Lindl.) S.W.L.Jacobs & J.Everett: Australia, SA, 35°10’0.09”S, 138°50’44.27”E, *D.J. Duval s.n.*, 01 Dec. 2006, collected seed at MSB no. 359926 (K). Plants grown from seed at BG Halle by G. Winterfeld GW80 (no voucher); LN832458*; LN832426*; #Acc1#. ***A. scabra*** subsp. ***scabra*: (1)** Australia, TAS, 42°41’33.7”S, 147°16’0”E, *J.A. Wood s.n.*, 03 Dec. 2007, collected seed at MSB no. 463744 (K). Plants grown from seed at BG Halle by G. Winterfeld GW112 (HAL0140650, HAL0137627); LN832459*; LN832427*; #Acc1#; **(2)** Australia, NSW, South Western Plains, c. 82 km N of Hillston, Mount Hope road, 32°55’52”S, 145°52’38”E, *S.W.L. Jacobs 9804 & M.E. Barkworth*, 13 Dec. 2007 (HAL0110793, duplicates: BA, NSW594289, UTC00252321); #matK#; #ITS#; ‒; LEP ill. ***A. scabra*** subsp. ***falcata*** (Hughes) S.W.L.Jacobs & J.Everett: **(1)** Australia, QLD, 26°29’1.68”S, 151’20’2.4”E, *D. Bowen & P. Boyle s.n.*, 25 Nov. 2005, collected seed at MSB no. 366454 (K). Plants grown from seed at BG Halle by G. Winterfeld GW63 (HAL0137626); LN832457*; #ITS#; #Acc1#; **(2)** Australia, NSW, Southern Tablelands, Bungonia turnoff, S of Marulan, Hume Highway, 34°43’59”S, 149°58’47”E, *S.W.L. Jacobs 9726 & M.E. Barkworth*, 23 Nov. 2007 (HAL0110781, duplicates: BA, NSW832479, UTC00252107, UTC); #matK#; #ITS#; #Acc1#; LEP ill. ***A. semibarbata*** (R.Br.) S.W.L.Jacobs & J.Everett: **(1)** Australia, SA, 34°57’51.26”S, 138°42’39.79”E, *R.J. Bates s.n.*, 15 Nov. 2006, collected seed at MSB no. 368768 (K). Plants grown from seed at BG Halle by G. Winterfeld GW64 (HAL0137628); #matK#; ‒; #Acc1#; **(2)** Australia, VIC, Gippsland Plain, 33 km E of Longford, Loch Sport Road, 38°08’26”S, 147°24’23”E, *S.W.L. Jacobs 9709*, 31 Oct. 2007 (HAL0110777, duplicates: MEL2330348A, NSW831776, UTC00252208); #matK#; #ITS#; ‒; LEP ill. ***A. setacea*** (R.Br.) S.W.L.Jacobs & J.Everett: **(1)** Australia, NSW, 34°4’4”S, 150°45’57”E, 18 Nov. 2003, collected seed at MSB no. 225014 (K). Plants grown from seed at BG Halle by G. Winterfeld GW55 (no voucher); LN832444*; LN832416*; #Acc1#; **(2)** Australia, VIC, Grampians, c. 11 km W of Natimuk, Mount Arapiles-Tooan State Park, 36°45’39”S, 141°50’13”E, *S.W.L. Jacobs 9781 & M.E. Barkworth*, 10 Dec. 2007 (HAL0110806, duplicates: MEL2330274A, NSW832509, UTC); #matK#; #ITS#; ‒; LEP ill.; **(3)** Australia, VIC, Grampians, c. 11 km W of Natimuk; Mount Arapiles-Tooan State Park, 36°45’39”S, 141°50’13”E, *S.W.L. Jacobs 9785 & M.E. Barkworth*, 10 Dec. 2007 (HAL0110715, duplicates: MEL2330242A, NSW832488, UTC); LEP. ***A. stipoides*** (Hook.f.) S.W.L. Jacobs & Everett: **(1)** Australia, NSW, South Coast, Bermagui, 36°25’39”S, 150°4’29”E, *S.W.L. Jacobs 9715 & M.E. Barkworth*, 01 Nov. 2007 (NSW832485, UTC00252109); #matK#; #ITS#; ‒; **(2)** Australia, SA, South-eastern, c. 6 km S of Meningie, Coorong road, 35°44’38”S, 139°20’03”E, *S.W.L. Jacobs 9774 & M.E. Barkworth*, 09 Dec. 2007 (HAL0110794, duplicates: AD228914, BA, NSW594287, UTC00252322); #matK#; #ITS#; ‒; LEP ill.; **(3)** Australia, SA, South-eastern, c. 40 km N of Kingston SE, Salt Creek Road, 36°29’33”S, 139°49’29”E, *S.W.L. Jacobs 9689*, 25 Oct. 2007 (HAL0110804, duplicates: AD229032, B, NSW831774, TCD, UTC00252157); #matK#; #ITS#; ‒; LEP. ***A. stuposa*** (Hughes) S.W.L.Jacobs & J.Everett: **(1)** Australia, VIC, East Gippsland, Swan Reach, Mettung Road, 37°49’29”S, 147°51’48”E, *S.W.L. Jacobs 9711*, 31 Oct. 2007 (HAL0110803, duplicates: MEL2330236A, NSW831782, UTC00252156); #matK#; #ITS#; #Acc1#; LEP ill.; **(2)** LN832450*; ‒; ‒. ***A. tenuifolia*** (Steud.) S.W.L.Jacobs & J.Everett: ‒; EU489104**; ‒. ***A. trichophylla*** (Benth.) S.W.L.Jacobs & J.Everett: Australia, WA, 31°37’59”S, 117°42’23”E, 01 Oct. 2004, collected seed at MSB no. 363604 (K). Plants grown from seed at BG Halle by G. Winterfeld GW81 (HAL0137635); LN832460*; LN832428*; #Acc1#. ***A. tuckeri*** (F.Muell) S.W.L.Jacobs & Everett: **(1)** Australia, NSW, North Western Plains, Hermidale, to SE of village, 31°32’46”S, 146°43’43”E, *S.W.L. Jacobs 9810 & M.E. Barkworth*, 13 Dec. 2007 (HAL0107514, duplicates: NSW594354, UTC); #matK#; #ITS#; ‒; LEP ill.; **(2)** Australia, NSW, South Western Plains, c. 3 km NE of Gilgunnia, Nymagee road, 32°23’51”S, 146°2’48”E, *S.W.L. Jacobs 9807 & M.E. Barkworth*, 13 Dec. 2007 (NSW594348); #matK#; #ITS#; ‒; LEP. ***A. variabilis*** (Hughes) S.W.L.Jacobs & J.Everett: Australia, WA, Eyre, c. 40 km S of Ravensthorpe, Hopetoun road, 33°52’26”S, 120°09’47”E, *S.W.L. Jacobs 9745 & M.E. Barkworth*, 02 Dec. 2007 (HAL0107542, duplicates: B, NSW832526, PERTH8500258, TCD, UTC); #matK#; #ITS#; ‒; LEP ill. ***A. velutina*** (Vickery, S.W.L.Jacobs & J.Everett) S.W.L.Jacobs & J.Everett: Australia, SA, Nullarbor, c. 135 km E of Border Village, Nullarbor road, 31°35’2”S, 130°22’21”E, *S.W.L. Jacobs 9758 & M.E. Barkworth*, 05 Dec. 2007 (AD229035, NSW832523); #matK#; #ITS#; ‒. ***A. verticillata*** (Nees ex Spreng.) S.W.L.Jacobs & J.Everett: **(1)** Australia, QLD, 28°50’7”S, 151°38’57”E, *T. Tyson-Doneley s.n.*, 23 Mar. 2006, collected seed at MSB no. 332235 (K). Plants grown from seed at BG Halle by G. Winterfeld GW71 (HAL0137625); LN832452*; LN832420*; #Acc1#; **(2)** Australia, NSW, Central Western Slopes, c, 36 km of Wellington, Molong road, 32°50’57”S, 148°54’53”E, *S.W.L. Jacobs 9811 & M.E. Barkworth*, 13 Dec. 2007 (HAL0107513, duplicates: BA, NSW594355, UTC); #matK#; #ITS#; ‒; LEP ill. ***A. vickeryana*** (J.Everett & S.W.L.Jacobs) S.W.L.Jacobs & J.Everett; morph. ***A. wakoolica*** (Vickery, S.W.L.Jacobs & J.Everett) S.W.L.Jacobs & J.Everett: ‒; JF769087**; ‒. ***Bromus erectus*** Huds.: AM234570*; FM179394*; ‒. ***B. inermis*** Leyss.: ‒; ‒; EU366392**. ***Celtica gigantea*** (Link) F.M.Vázquez & Barkworth: FN434281*; FN434544*; ‒. ***Henrardia persica*** C.E.Hubb.: ‒; ‒; GQ228396**. ***Hordeum chilense*** Roem. & Schult.: ‒; ‒; DQ497805**; ‒. ***H. vulgare*** L.: KC912687**; FN556796**; AF343509**. ***Macrochloa tenacissima*** (L.) Kunth: Spain, *R. Piwowarczyk s.n.* (KRA); LEP ill. ***Nassella neesiana*** (Trin. & Rupr.) Barkworth: **(1)** Australia, NSW, Southern Tablelands, c. 18 km E of Bungonia, Goulburn Road, 34°48’28”S, 149°46’41”E, *S.W.L. Jacobs 9729 & M.E. Barkworth*, 23 Nov. 2007 (HAL0110790, duplicates: NSW832484, TCD, UTC); #matK#; #ITS#; ‒; **(2)** Italy, Roma, *Moraldo s.n.* (KRA); LEP ill. ***N. trichotoma*** (Nees) Hack. & Arechav.: Australia, NSW, Southern Tablelands, c. 5 km S of Marulan, Prarie Oak Road on Bungonia-Marulan Road, 34°45’17”S, 149°58’17”E, *S.W.L. Jacobs 9727 & M.E. Barkworth*, 23 Nov. 2007 (HAL0110791, duplicates: B, BA, K, NSW832483, TCD, UTC); #matK#; #ITS#; #Acc1#. ***Neotrinia splendens*** (Trin.) M.Nobis, P.D.Gudkova & A.Nowak: **(1)** FN434208*; FN434477*; ‒; **(2)** Tajikistan, *Yu. Gusev s.n.*, s.d. (LE); LEP ill. ***Oloptum miliaceum*** (L.) Röser & Hamasha: LN832471*, AM234597*; LN832439*, FM179427*; ‒. ***Orthoraphium roylei*** Nees: *J.F. Duthie 3558* (LE); LEP ill. ***Ptilagrostis concinna*** (Hook.f.) Roshev.: India, Ladakh, *L. Klimes s.n.* (KRA); LEP ill. ***P. mongholica*** (Turcz. ex Trin) Griseb.: Mongolia, *A. Pacyna s.n.* (KRA); LEP ill. ***Secale sylvestre*** Host: AM234581*; FM179434*; ‒. ***Stipa capillata*** L.: Germany,-Saxony-Anhalt, plants grown in BG Halle, *M. Röser 11083*, 16 June 2011 (HAL); ‒; ‒; #Acc1#. ***S. drobovii*** (Tzvel.) Czerep.: Tajikistan, *M. Nobis s.n.* (KRA); LEP ill. ***S. kirghisorum*** P.A.Smirn.: Tajikistan, *M. Nobis s.n.* (KRA); LEP ill. ***S. pennata*** subsp. ***eriocaulis*** (Borbás) Martinovský & Skalický: ‒; FM179438*; ‒. ***S. pennata*** L. subsp. ***zalesskii*** (Wilensky) Freitag: FN434317*; ‒; ‒. ***S. tirsa*** Steven: **(1)** Germany, Saxony-Anhalt, plants grown in BG Halle, *M. Röser 11084*, 16 June 2011 (HAL): FN434350*; FN434605*; #Acc1#; **(2)** Georgia, *M. Nobis s.n.* (KRA); LEP ill. ***Stipellula capensis*** (Thunb.) Röser & Hamasha: **(1)** FN434257*; FN434522*; ‒; **(2)** Spain, *R. Piwowarczyk s.n.* (KRA); LEP ill.

## Appendix 2. Annotated list of chromosome number reports for taxa of Stipeae

Underlined are the most frequent chromosome numbers in case of different countings, if this was stated in the original publications or if concluded from this survey (mostly for the genera). Original publications we did not examine are identified as such in the list of references below.

***Achnatherum*** P.Beauv.: **2*n* = 18**, **20**, **22**, **24**, **28**, **42**, **44**, **46**, **48**. — ***A***. ***brandisii*** (Mez) Z.L.Wu: ***n* = 12**: Kumari & al. (2019) as *Stipa brandisii* Mez. ***A***. ***bromoides*** (L.) P.Beauv.: **2*n* = 18**: Petrova (1968); **2*n* = 24**: Strid & Andersson (1985) as *Stipa bromoides* (L.) Dörfl.; Ghukasyan (2004); **2*n* = 28**: Löve & Löve (1961) reporting an unpubl. count of Uihelyi; Vázquez & Devesa (1996) as *S. bromoides* (L.) Dörfl. ***A***. ***calamagrostis*** (L.) P.Beauv.: **2*n* = 22 + 0‒2B**: Strid (1983); **2*n* = 24**: Gervais (1966); Löve & Löve (1974); Strobl & Wittmann (1985); Kožuharov & Petrova (1991). ***A***. ***caragana*** (Trin.) Nevski: **2*n* = 24**: Sokolovskaya & Probatova (1978). ***A***. ***confusum*** (Litv.) Tzvelev: **2*n* = 24**: Sokolovskaya & Probatova (1977); Probatova & Seledets (2008); Probatova & al. (2008, 2015). ***A***. ***dregeanum*** (Steud.) Röser, Tkach & M.Nobis: **2*n* = 48**: de Winter (1965) as *Stipa dregeana* Steud. ***A***. ***jacquemontii*** (Jaub. & Spach) P.C.Kuo & S.L.Lu: ***n* = 10**: Kumar & al. (2018) as *Stipa jacquemontii* Jaub. & Spach; ***n* = 12**: Mehra & Sharma (1975a) as *S. jacquemontii*; ***n* = 21**, **22**: Gupta & al. (2014) as *S. jacquemontii*. ***A***. ***paradoxum*** (L.) Banfi, Galasso & Bartolucci: **2*n* = 24**: Litardière (1950) as *Oryzopsis paradoxa* (L.) Nutt.; Löve & Kjellqvist (1973) as *O. paradoxa*. ***A***. ***pekinense*** (Hance) Ohwi: **2*n* = 24**: Ono & Tateoka (1953) as *Stipa extremiorientalis* Hara; Tateoka (1954, 1967, 1986); Sokolovskaya & Probatova (1977) as *Achnatherum extremiorientale* (Hara) Keng; Korobkov & al. (2013) as *A. extremiorientale*. ***A***. ***petriei*** (Buchanan) S.W.L.Jacobs & J.Everett: **2*n* = 42**: Murray & al. (2005). ***A***. ***sibiricum*** (L.) Keng ex Tzvelev: ***n* = 12**, **2*n* = 24:** Johnson (1945) as *Stipa sibirica* (L.) Lam.; **2*n* = 22**, **23**: Měsíček & Soják (1972) as *S. sibirica*; **2*n* = 22**: Stepanov (1994) as *S. sibirica*; **2*n* = 24**: Avdulov (1928, 1931: 129) as *S. sibirica*; Myers (1947) citing an unpubl. count of R.M. Love as *S. sibirica*; Belyaeva & Siplivinsky (1977) as *S. sibirica*; Sokolovskaya & Probatova (1978); Gusik & Levkovskiy (1979) as *S. sibirica*; Probatova & Sokolovskaya (1980); Rudyka (1990); Probatova & al. (2009, 2012, 2015, 2016a); Chepinoga & al. (2012); Winterfeld & al. (2015). ***A***. ***turcomanicum*** (Roshev.) Tzvelev: **2*n* = 24**: Chopanov & Yurtzev (1976) as *Stipa litwinowiana* P.A.Smirn. ex Pavlov & Lipsch. ***A***. ***virescens*** (Trin.) Banfi, Galasso & Bartolucci: **2*n* = 24**: Avdulov (1931: 131) as *Oryzopsis virescens* (Trin.) Beck; Májovský 1974 as *O. virescens*.

***Aciachne*** Benth.: **2*n* = 22**. — ***A***. ***pulvinata*** Benth.: **2*n* = 22**: Reeder & Reeder (1968).

***Amelichloa*** Arriaga & Barkworth: **2*n* = 40**, **44**, **44‒46**. — ***A***. ***brachychaeta*** (Godr.) Arriaga & Barkworth: **2*n* = 40**: Saura (1943) as *Stipa brachychaeta* Godr.; **2*n* = 40**, **44**: Saura (1948) as *S. brachychaeta*; **2*n* = 44**: Parodi (1946) as *S. brachychaeta*; Bowden & Senn (1962) as *S. brachychaeta*; **2*n* = 44‒46**: Myers (1947) citing an unpubl. count of G.L. Stebbins as *S. brachychaeta*.

***Anatherostipa*** (Hack. ex Kuntze) Peñail.: **2*n* = 22**. — ***A***. ***hans-meyeri*** (Pilg.) Peñail.: ***n* = 11** Davidse & Pohl (1978) as *Stipa hans-meyeri* Pilg.

***Anemanthele*** Veldkamp: **2*n* = 40**, **42**, **<u>44</u>**, **46**. — ***A***. ***lessoniana*** (Steud.) Veldkamp: **2*n* = 40‒ 44**: Dawson & Beuzenberg (2000); Edgar & Connor (2000) citing an unpubl. count of M.I. Dawson; **2*n* = 42**, **<u>44</u>**, **46**: Winterfeld & al. (2015).

***Austrostipa* S**.**W**.**L**.**Jacobs & J**.**Everett**: **2*n* = 42**, **<u>44</u>**, **64**, **<u>66</u>**, **68**, **70**. — ***A***. ***acrociliata*** (Reader) S.W.L.Jacobs & J.Everett: **2*n* = 44**: Winterfeld & al. (2015). ***A***. ***blackii*** (C.E.Hubb.) S.W.L.Jacobs & J.Everett: **2*n* = 44**: Winterfeld & al. (2015). ***A***. ***breviglumis*** (J.M.Black) S.W.L.Jacobs & J.Everett: **2*n* = 44**: Winterfeld & al. (2015). ***A***. ***campylachne*** (Nees) S.W.L.Jacobs & J.Everett: **2*n* = 66**: Winterfeld & al. (2015). ***A***. ***curticoma*** (Vickery) S.W.L.Jacobs & J.Everett: **2*n* = 66**: Winterfeld & al. (2015). ***A***. ***drummondii*** (Steud.) S.W.L.Jacobs & J.Everett: **2*n* = 44**: Winterfeld & al. (2015). ***A***. ***elegantissima*** (Labill.) S.W.L.Jacobs & J.Everett: **2*n* = 66**: Winterfeld & al. (2015). ***A***. ***exilis*** (Vickery) S.W.L.Jacobs & J.Everett: **2*n* = 44**: Winterfeld & al. (2015). ***A***. ***flavescens*** (Labill.) S.W.L.Jacobs & J.Everett: **2*n* = 44**: Winterfeld & al. (2015). ***A***. ***geoffreyi*** S.W.L.Jacobs & J.Everett: **2*n* = 44**: Winterfeld & al. (2015). ***A***. ***hemipogon*** (Benth.) S.W.L.Jacobs & J.Everett. **2*n* = 66**, **70**: Winterfeld & al. (2015). ***A***. ***mollis*** (R.Br.) S.W.L.Jacobs & J.Everett: **2*n* = 64**, **66**: Winterfeld & al. (2015). ***A***. ***nodosa*** (S.T.Blake) S.W.L.Jacobs & J.Everett: **2*n* = 44**: Winterfeld & al. (2015). ***A***. ***nullanulla*** (S.W.L.Jacobs & J.Everett) S.W.L.Jacobs & J.Everett: **2*n* = 44**: Winterfeld & al. (2015). ***A***. ***oligostachya*** (Hughes) S.W.L.Jacobs & J.Everett: **2*n* = 44**: Winterfeld & al. (2015). ***A***. ***pilata*** (S.W.L.Jacobs & J.Everett) S.W.L.Jacobs & J.Everett: **2*n* = 44**: Winterfeld & al. (2015). ***A***. ***platychaeta*** (Hughes) S.W.L.Jacobs & J.Everett: **2*n* = 44**: Winterfeld & al. (2015). ***A***. ***ramosissima*** (Trin.) S.W.L.Jacobs & J.Everett: **2*n* = 44**: Winterfeld & al. (2015). ***A***. ***scabra*** (Lindl.) S.W.L.Jacobs & J.Everett: **2*n* = 42**, **44**: Winterfeld & al. (2015). ***A***. ***scabra*** (Lindl.) S.W.L.Jacobs & J.Everett subsp. ***falcata*** (Hughes) S.W.L.Jacobs & J.Everett: **2*n* = 44**: Winterfeld & al. (2015). ***A***. ***scabra*** (Lindl.) S.W.L.Jacobs & J.Everett subsp. ***scabra***: **2*n* = 60**, **<u>62</u>**: Winterfeld & al. (2015). ***A***. ***semibarbata*** (R.Br.) S.W.L.Jacobs & J.Everett: **2*n* = 65**, **66**, **<u>68</u>**, **<u>70</u>**: Winterfeld & al. (2015). ***A***. ***stipoides*** (Hook.f.) S.W.L.Jacobs & J.Everett: **2*n* = 44**: Murray & al. (2005); Winterfeld & al. (2015). ***A***. ***stuposa*** (Hughes) S.W.L.Jacobs & J.Everett: **2*n* = 66**: Winterfeld & al. (2015). ***A***. ***trichophylla*** (Benth.) S.W.L.Jacobs & J.Everett: **2*n* = 44**: Winterfeld & al. (2015). ***A***. ***verticillata*** (Nees ex Spreng.) S.W.L.Jacobs & J.Everett: **2*n* = 44**: Winterfeld & al. (2015).

***Barkworthia*** Romasch., P.M.Peterson & Soreng: **2*n* = 40**. — ***B***. ***stillmanii*** (Bol.) Romasch., P.M.Peterson & Soreng: **2*n* = 40**: Myers (1947) citing an unpubl. count of G.L. Stebbins as *Stipa stillmanii* Bol.

***Celtica*** F.M.Vázquez & Barkworth: **2*n* = 84**, **<u>96</u>**. — ***C***. ***gigantea*** (Link) F.M.Vázquez & Barkworth: ***n* = 48**: Devesa & al. (1991) as *Stipa gigantea* Link; Gallego & Talavera (1994) as *S. gigantea*; **2*n* = 84**: Winterfeld (2006) as *S. gigantea*; **2*n* = 96**: Fernandes & Queirós (1969) as *S. gigantea*; Queirós (1973) as *S. gigantea*; Queirós (1974) as *S. gigantea*. ***C***. ***gigantea*** subsp. ***gigantea***: **2*n* = 96**: Vázquez & Devesa (1996) as *S. gigantea* subsp. *gigantea*. ***C***. ***gigantea*** subsp. ***donyanae*** (F.M.Vázquez & Devesa) F.M.Vázquez & Barkworth: **2*n* = 96**: Vázquez & Devesa (1996) as *S. gigantea* subsp. *donyanae* F.M.Vázquez & Devesa.

***Eriocoma*** Nutt.: **2*n* = <u>32</u>**, **<u>34</u>**, **<u>36</u>**, **40**, **<u>44</u>**, **48**, **64**, **66**, **<u>68</u>**, **70**. — ***E***. ***contracta*** (B.L.Johnson) Romasch.: ***n* = 24**: Reeder (1977) as *Oryzopsis contracta* (B.L.Johnson) Schechter; **2*n* = 48**: Shechter & Johnson (1966) as *O. contracta*. ***E***. ***coronata*** (Thurb.) Romasch.: **2*n* = 40**: Stebbins & Love (1941) as *Stipa coronata* Thurb.; Myers (1947) citing an unpubl. count of R.M. Love as *S. coronata*; Reeder (1984) as *S. coronata*. ***E***. ***curvifolia*** (Swallen) Romasch.: ***n* = 22**: Hatch & Bearden (1983) as *Stipa curvifolia* Swallen. ***E***. ***hendersonii*** (Vasey) Romasch.: ***n* = 17**: Hitchcock & Spellenberg (1968) as *Oryzopsis hendersonii* Vasey; ***n* = 17**, **2*n* = 34**: Spellenberg (1968) as *O. hendersonii*. ***E***. ***hymenoides*** (Roem. & Schult.) Rydb.: ***n* = 24**: Reeder (1977) as *Oryzopsis hymenoides* (Roem. & Schult.) Ricker ex Piper; ***n* = 24**, **2*n* = 48**: Johnson (1945, 1960, 1962a) as *O. hymenoides*; **2*n* = 48**: Stebbins & Love (1941) as *O. hymenoides*; Johnson & Rogler (1943) as *O. hymenoides*; Johnson (1960) as *O. hymenoides*; Löve & Löve (1981) as *O. hymenoides*; **2*n* = 48 + 0‒5B**: Johnson (1963) as *O. hymenoides*. ***E***. ***latiglumis*** (Swallen) Romasch.: ***n* = 35**: Pohl (1954) as *Stipa latiglumis* Swallen representing an allopolyploid hybrid between *S. lettermanii* and *S. elmeri*. ***E***. ***lemmonii*** (Vasey) Romasch.: ***n* = 17**: Pohl (1954) as *Stipa lemmonii* (Vasey) Scribn.; Reeder (1977) as *S. lemmonii*; **2*n* = 34**: Myers (1947) citing an unpubl. count of R.M. Love as *S. lemmonii*; ***n* = 18**, **2*n* = 36**: Stebbins & Love (1941) as *S. lemmonii*; Johnson (1962a) as *S. columbiana* Macoun; **2*n* = 44**: Nielsen (1939) as *S. columbiana*. ***E***. ***lemmonii*** subsp. ***pubescens*** (Crampton) Romasch.: **2*n* = 34**: Stebbins & Love (1941) as *Stipa lemmonii* var. *jonesii* Scribn. ***E***. ***lettermanii*** (Vasey) Romasch.: ***n* = 16**, **2*n* = 32**: Johnson (1962a) as *S. lettermanii*; **2*n* = 32**: Dedecca (1954) as *S. lettermanii* and commenting that this plant with 2*n* = 32 was true *S. lettermanii*, whereas the plant of Stebbins & Love (1941) with 2*n* = 66 belonged to an undescribed polyploid; **2*n* = 66**: Stebbins & Love (1941) as *Stipa lettermanii* Vasey var.; **2*n* = 68**: Myers (1947) citing an unpubl. count of R.M. Love as *S. lettermanii*. ***E***. ***nevadensis*** (B.L.Johnson) Romasch.: ***n* = 34**, **2*n* = 68**: Johnson (1962a) as *Stipa nevadensis* B.L.Johnson and representing an amphidiploid hybrid between *S. lettermanii* and *S. occidentalis* or *S. elmeri*; **2*n* = 68**: Johnson (1962b) as *S. nevadensis*. ***E***. ***occidentalis*** (Thurb. ex S.Watson) Romasch.: **2*n* = 36**: Stebbins & Love (1941) as *Stipa occidentalis* Thurb. ***E***. ***occidentalis*** subsp. ***californica*** (Merr. & Burtt Davy) Romasch.: ***n* = 18**: Reeder (1977) as *Stipa californica* Merr. & Burtt Davy; ***n* = 18**, **2*n* = 36**: Stebbins & Love (1941) as *S. californica*; Johnson (1962a) as *S. occidentalis* Thurb. ***E***. ***occidentalis*** subsp. ***pubescens*** (Vasey) Romasch.: ***n* = 18**: Pohl (1954) as *Stipa elmeri* Piper & Brodie ex Scribn.; ***n* = 18**, **2*n* = 36**: Stebbins & Love (1941) as *S. elmeri*; Johnson (1945, 1962a) as *S. elmeri*. ***E***. ***pinetorum*** (M.E.Jones) Romasch.: ***n* = 16**, **2*n* = 32**: Johnson (1963) as *Stipa pinetorum* M.E.Jones; **2*n* = 32**: both Johnson (1945) and Myers (1947) and citing an unpubl. count of G.L. Stebbins as *S. pinetorum*. ***E***. ***richardsonii*** (Link) Romasch.: **2*n* = 44**: Löve & Löve (1981) as *Stipa richardsonii* Link. ***E***. ***robusta*** (Vasey) Romasch.: **2*n* = 64**: Myers (1947) citing an unpubl. count of R.M. Love as *Stipa robusta* (Vasey) Scribn. ***E***. ***scribneri*** (Vasey) Romasch.: **2*n* = 40**: Tateoka (1955) as *Stipa scribneri* Vasey. ***E***. ***swallenii*** (C.L.Hitchc. & Spellenb.) Romasch.: ***n* = 17**, **2*n* = 34**: Hitchcock & Spellenberg (1968) as *Oryzopsis swallenii* C.L.Hitchc. & Spellenb. ***E***. ***thurberiana*** (Piper) Romasch.: ***n* = 17**, **2*n* = 34**: Stebbins & Love (1941) as *Stipa thurberiana* Piper. ***E***. ***webberi*** Thurb.: ***n* = 16**, **2*n* = 32**: Johnson (1945) as *Stipa webberi* (Thurb.) B.L.Johnson.

***Hesperostipa*** (M.K.Elias) Barkworth: **2*n* = 38**, **<u>44</u>**, **46**. — ***H***. ***comata*** (Trin. & Rupr.) Barkworth: ***n* = 19**, **22**: Reeder (1977) as *Stipa comata* Trin. & Rupr.; **2*n* = 46**: Myers (1947) citing an unpubl. count of R.M. Love as *S. comata*. ***H***. ***comata*** subsp. ***comata***: **2*n* = 44**: Bowden (1960) as *S. comata*; Löve & Löve (1981) as *S. comata*. ***H***. ***comata*** subsp. ***intermedia*** (Scribn. & Tweedy) Barkworth: **2*n* = 44‒46**: Stebbins & Love (1941) as *Stipa comata* var. *intermedia* Scribn. & Tweedy. ***H***. ***curtiseta*** (Hitchc.) Barkworth: **2*n* = 44**: Löve & Löve (1981) as *Stipa curtiseta* (Hitchc.) Barkworth. ***H***. ***neomexicana*** (Thurb.) Barkworth: ***n* = 22**: Gould (1968) as *Stipa neomexicana* (Thurb.) Scribn.; **2*n* = 44**: Myers (1947) citing an unpubl. count of R.M. Love as *S. neomexicana*. ***H***. ***spartea*** (Trin.) Barkworth: **2*n* = 44**: Tateoka (1955) as *Stipa spartea* Trin.; Löve & Löve (1981) as *S. spartea*; **2*n* = 46**: Myers (1947) citing an unpubl. count of R.M. Love as *S. spartea*.

***Jarava*** Ruiz & Pav.: **2*n* = 40**, **42**, **<u>44</u>**, **66**. — ***J***. ***ichu*** Ruiz & Pav.: ***n* = 20**: Pohl & Davidse (1971) as *Stipa ichu* (Ruiz & Pav.) Kunth; Davidse & Pohl (1974) as *S. ichu*; ***n* = 20**, **ca**. **20**: Gould (1966) as *S. ichu*; **2*n* = 42**: Bowden & Senn (1962) as *S. ichu*; **2*n* = 44**: Covas (1945) as *Stipa gynerioides* Phil.; Saura (1948) as *S. ichu*; Tateoka (1962) as *S. ichu. **J***. ***neaei*** (Nees ex Steud.) Peñail.: **2*n* = 66**: Covas & Bocklet (1945) as *Stipa neaei* Nees ex Steud. ***J***. ***plumosa*** (Spreng.) S.W.L.Jacobs & J.Everett: **2*n* = 40**: Bowden & Senn (1962) as *Stipa papposa* Nees; **2*n* = 44**: Avdulov (1928, 1931: 129) as *S. papposa* Nees; Lorenzo-Andreu (1953) as *S. papposa* Nees. ***J***. ***plumosula*** (Nees ex Steud.) F.Rojas: **2*n* = 44**: Covas & Bocklet (1945) as *Stipa plumosa* Trin. & Rupr.

***Lorenzochloa*** Reeder & C.Reeder: **2*n* = 22**. — ***L***. ***erectifolia*** (Swallen) Reeder & C.Reeder: **2*n* = 22**: Reeder & Reeder (1968) as *Parodiella erectifolia* (Swallen) Reeder & C.Reeder.

***Macrochloa*** Kunth; **2*n* = 24**, **40**, **<u>66</u>**, **72**. — ***M***. ***tenacissima*** (L.) Kunth: ***n* = 33**: Ferchichi & al. (1994) as *Stipa tenacissima* L.; **2*n* = 24**: Labadie (1979a,b) as *S. tenacissima*; **2*n* = 40**: Fernandes & Queirós (1969) as *S. tenacissima*; **2*n* = 66**: Lungeanu (1980) as *S. tenacissima*; ***n* = 36**: Faruqi & al. (1987) as *S. tenacissima*.

***Nassella*** (Trin.) É.Desv.: **2*n* = 26**, **<u>28</u>**, **32**, **<u>34</u>**, **<u>36</u>**, **38**, **42**, **56**, **58**, **60**, **64**, **<u>66</u>**, **70**, **82**, **88**. — ***N cernua*** (Stebbins & Love) Barkworth: ***n* = 35**: Love (1954) as *Stipa cernua* Stebbins & Love; ***n* = 35**, **2*n* = 70**: Stebbins & Love (1941) as *S. cernua*; **2*n* = 70**: Love (1946) as *S. cernua*. ***N***. ***charruana*** (Arechav.) Barkworth: **2*n* = 36**: Bowden & Senn (1962) as *Stipa charruana* Arechav. ***N***. ***chilensis*** (Trin.) É.Desv.: **2*n* = 42**: Bowden & Senn (1962). ***N***. ***gigantea*** (Steud.) Muñoz-Schick: **2*n* = 42**: Bowden & Senn (1962) as *N. exserta* Phil. ***N***. ***hyalina*** (Nees) Barkworth: **2*n* = 34**: Bowden & Senn (1962) as *Stipa hyalina* Nees. ***N***. ***lachnophylla*** (Trin.) Barkworth: **2*n* = 66**: Bowden & Senn (1962) as *Stipa laxa* É.Desv. ***N***. ***lepida*** (Hitchc.) Barkworth: ***n* = 17**: Love (1954) as *Stipa lepida* Hitchc.; ***n* = 17**, **2*n* = 34**: Stebbins & Love (1941) as *S. lepida*; **2*n* = 34**: Love (1946) as *S. lepida*; **2*n* = 46**: Reeder (1967) as *S. lepida*. ***N***. ***leucotricha*** (Trin. & Rupr.) R.W.Pohl: ***n* = 13**: Brown (1949) as *Stipa leucotricha* Trin. & Rupr.; ***n* = 14**: Gould (1958) as *S. leucotricha*; Hatch (1980) as *S. leucotricha*; **2*n* = 26**: Brown (1951) as *S. leucotricha*; **2*n* = 28**: Myers (1947) citing an unpubl. count of R.M. Love as *S. leucotricha*. ***N***. ***megapotamia*** (Spreng. ex Trin.) Barkworth: **2*n* = 34**: Myers (1947) citing an unpubl. count of G.L. Stebbins as *Stipa megapotamica* Spreng. ex Trin. & Rupr. ***N***. ***mucronata*** (Kunth) R.W.Pohl: **2*n* = 36**: Myers (1947) citing an unpubl. count of R.M. Love as *Stipa mucronata* Kunth; **2*n* = 64**: Bowden & Senn (1962) as *S. mucronata*. ***N***. ***neesiana*** (Trin. & Rupr.) Barkworth: **2*n* = 28**: Myers (1947) citing an unpubl. count of G.L. Stebbins as *Stipa neesiana* Trin. & Rupr.; Brown (1949) as *S. neesiana*; Bowden & Senn (1962) as *S. neesiana*. ***N***. ***philippii*** (Steud.) Barkworth: **2*n* = 36**: Myers (1947) citing an unpubl. count of G.L. Stebbins as *Stipa philippii* Steud. ***N***. ***pulchra*** (Hitchc.) Barkworth: **2*n* = 55**, **56**, **58**, **<u>60</u>**: Winterfeld & al. (2015); ***n* = 32**: Love (1954) as *Stipa pulchra* Hitchc.; ***n* = 32**, **2*n* = 64**: Stebbins & Love (1941) as *S. pulchra*; **2*n* = 64**: Love (1946) as *S. pulchra*; **2*n* = 66**: Nielsen (1939) as *S. pulchra*. ***N***. ***tenuissima*** (Trin.) Barkworth: **2*n* = 32**: Brown (1951) as *Stipa tenuissima* Trin.; **2*n* = 40**: Bowden & Senn (1962) as *S. tenuissima*. ***N***. ***trichotoma*** (Nees) Hack. ex Arechav.: **2*n* = 36**: Bowden & Senn (1962); **2*n* = 38**: Avdulov (1928) as *Urachne trichotoma* (Nees) Trin. fide Fedorov (1969). ***N***. ***viridula*** (Trin.) Barkworth: ***n* = 41**, **2*n* = 82**: Johnson & Rogler (1943) as *Stipa viridula* Trin.; **2*n* = 88**: Löve & Löve (1981) as *S. viridula*.

***Neotrinia*** (Tzvelev) M.Nobis, P.D.Gudkova & A.Nowak. **2*n* = <u>44</u>**, **46**, **<u>48</u>**. — *N. splendens* (Trin.) M.Nobis, P.D.Gudkova & A.Nowak: **2*n* = 42**: Sokolovskaya & Strelkova (1948) as *Lasiagrostis splendens* (Trin.) Kunth fide Probatova & Sokolovskaya (1980) regarding it as miscount; ***n* = 22**: Gupta & al. (2014) as *Stipa splendens* Trin.; **2*n* = 44**: Winterfeld & al. (2015) as *Achnatherum splendens* (Trin.) Nevski; ***n* = 23**: Gohil & Koul (1986) as *S. splendens*; **2*n* = 48**: Myers (1947) citing an unpubl. count of R.M. Love as *S. splendens* Trin.; **2*n* = 48**: Probatova & Sokolovskaya (1980) as *A. splendens*; **2*n* = ca**. **48**: Probatova & Seledets (2008) as *A. splendens*.

***Oloptum*** Röser & Hamasha: **2*n* = <u>24</u>**, **36**. — ***O***. ***miliaceum*** (L.) Röser & Hamasha: ***n* = 12**: Faruqi & al. (1987) as *Pipthaterum miliaceum* (L.) Cosson; Devesa & al. (1991) as *P. miliaceum*; ***n* = 12**, **2*n* = 24**: Johnson (1945) as *Oryzopsis miliacea* (L.) Benth. & Hook.f. ex Asch. & Schweinf.; **2*n* = 24**: Avdulov (1928, 1931: 131) as *O. miliacea*; Tateoka (1957) as *O. miliacea*; Fernandes & Queirós (1969) as *O. miliacea*; Luque & Diaz Lifante (1991) as *P. miliaceum*; Verlaque & al. (1997) as *P. miliaceum*; Winterfeld & al. (2015); ***n* = 18**: Ferchichi & al. (1994) as *O. miliacea* and considering it a triploid plant.

***Ortachne*** Nees: **no chromosome counts available**.

***Orthoraphium*** Nees: **no chromosome counts available**.

***Oryzopsis*** Michx.: **2*n* = 46**, **48**: — ***O***. ***asperifolia*** Michx.: **2*n* = 46**: Johnson (1945); **2*n* = 48**: Bowden (1960); Löve & Löve (1981).

***Pappostipa*** (Speg.) Romasch., P.M.Peterson & Soreng: **2*n* = 42**, **60**, **64**, **<u>66</u>**, **68**, **ca**. **74**. — ***P***. ***humilis*** (Cav.) Romasch.: **2*n* = 42(‒44)**: Myers (1947) citing an unpubl. count of G.L. Stebbins as *Stipa humilis* Cav.; **2*n* = 66**: Covas & Bocklet (1945) as *S. humilis*.

***P***. ***speciosa*** (Trin. & Rupr.) Romasch: **2*n* = 60**: Delay (1950) as *Stipa speciosa* Trin. & Rupr.; Stebbins & Love (1941) as *S. speciosa*; ***n* = 32**, **2*n* = 64**: Johnson (1960, 1962a) as *S. speciosa*; **2*n* = 66**: Covas & Bocklet (1945) as *S. speciosa* var. *macrochaeta* Parodi; **2*n* = 68**: Myers (1947) citing an unpubl. count of R.M. Love as *S. speciosa*; ***n* = ca**. **37**: Gould (1968) as *S. speciosa*.

***Patis*** Ohwi: **2*n* = <u>46</u>**, **48**. — ***P***. ***coreana*** (Honda) Ohwi: **2*n* = 46**: Tateoka (1986) as *Orthoraphium coreanum* var. *kengii* (Ohwi) Ohwi. ***P***. ***racemosa*** (Sm.) Romasch., P.M.Peterson & Soreng: **2*n* = 46**: Johnson (1945) as *Oryzopsis racemosa* (Sm.) Ricker ex Hitchc.; **2*n* = 48**: Bowden (1960) as *O. racemosa*.

***Piptatheropsis*** Romasch., P.M.Peterson & Soreng: **2*n* = 20**, **<u>22</u>**, **24**. — ***P***. ***canadensis*** (Poir.) Romasch., P.M.Peterson & Soreng: **2*n* = 22**: Spellenberg (1970) as *Oryzopsis canadensis* (Poir.) Torr., not Johnson (1945), erroneously cited by Curto & Henderson (1998). ***P***. ***exigua*** (Thurb.) Romasch., P.M.Peterson & Soreng: **2*n* = 22**: Hitchcock & Spellenberg (1968) as *Oryzopsis exigua* Thurb. ***P. micrantha*** (Trin. & Rupr.) Romasch., P.M.Peterson & Soreng: **2*n* = 22**: Johnson (1945) as *Oryzopsis micrantha* (Trin. & Rupr.) Thurb.; **2*n* = 24**: Löve & Löve (1981) as *O. micrantha*. ***P***. ***pungens*** (Torr. ex Spreng.) Romasch., P.M.Peterson & Soreng: **2*n* = 22**: Johnson (1945) as *Oryzopsis pungens* (Torr. ex Spreng.) Hitchc.; **2*n* = 24**: Bowden (1960) as *O. pungens*; Löve & Löve (1981) as *O. pungens*. ***P***. ***shoshoneana*** (Curto & Douglass M.Hend.) Romasch., P.M.Peterson & Soreng: ***n* = 10**, **2*n* = 20**: Curto & Henderson (1998) as *Stipa shoshoneana* Curto & Douglass M.Hend.

***Piptatherum*** P.Beauv.: **2*n* = 20**, **22**, **<u>24</u>**, **26**. — ***P***. ***aequiglume*** (Duthie ex Hook.f.) Roshev.: ***n* = 12**: Mehra & Sharma (1975a) as *Oryzopsis aequiglumis* Duthie ex Hook.f. ***P***. ***coerulescens*** (Desf.) P.Beauv.: **2*n* = 24**: Kerguélen (1975) as *Oryzopsis coerulescens* (Desf.) Hack. ***P***. ***holciforme*** (M.Bieb.) Roem. & Schult.: **2*n* = 24**: Johnson (1945) as *Oryzopsis holciformis* (M.Bieb.) Hack.; Petrova (1968). ***P***. ***gracile*** Mez: ***n* = 12**, **2*n* = 24**: Moinuddin & al. (1994). ***P***. ***laterale*** (Munro ex Regel) Munro ex Nevski: ***n* = 12**: Mehra & Sharma (1975b, 1977) as *Oryzopsis lateralis* (Munro ex Regel) Stapf ex Hook.f.; Gupta & al. (2014) as *O. lateralis*; Singhal & al. (2014) as *O. lateralis*. ***P***. ***microcarpum*** (Pilg.) Tzvelev: ***n* = 12**: Kaur & al. (2011a,b); **2*n* = 24**: Podlech & Dieterle (1969) as *Oryzopsis microcarpa* Pilg. ***P. molinioides*** Boiss.: ***n* = 11 + 1B**: Mehra & Sunder (1969) as *Oryzopsis molinioides* (Boiss.) Hack. ex Paulsen; ***P***. ***munroi*** (Stapf) Mez: **2*n* = 22**: Mehra & Sunder (1969) as *Oryzopsis munroi* Stapf; ***n* = 12**: Sharma & Sharma (1978, 1979) as *O. munroi*; Moinuddin & al. (1994); Singhal & al. (2014) as *O. munroi*. ***P***. ***songaricum*** (Trin. & Rupr.) Roshev. ex Nikitina subsp. ***songaricum***: **2*n* = 24**: Sokolovskaya & Probatova (1978). ***P***. ***vicarium*** (Grig.) Roshev. ex Nikitina: **2*n* = 20**: Chopanov & Yurtzev (1976); **2*n* = <u>24</u>**, **26**: Sokolovskaya & Probatova (1978). ***P***. ***virescens*** (Trin.) Boiss.: **2*n* = 24**: Avdulov (1928, 1931: 131) as *Oryzopsis virescens* (Trin.) Beck; Májovský (1974) as *O. virescens*; Sokolovskaya & Probatova (1978); Kožuharov & Petrova (1991); Probatova & Seledets (2008); Marhold & al. (2007) as *O. virescens*.

***Piptochaetium*** J.Presl: **2*n* = <u>22</u>**, **42**, **44**, **ca**. **60**. — ***P***. ***avenaceum*** (L.) Parodi: ***n* = 11**: Gould (1958) as *Stipa avenacea* L.; Valencia & Costas (1968). ***P***. ***bicolor*** (Vahl) É.Desv.: **2*n* = 22**: Covas & Bocklet (1945); Valencia & Costas (1968) as *P. bicolor* var. *minor* (Speg.) Parodi; Valencia & Costas (1968) as *P. bicolor* var. *typicum* (Speg.) Parodi. ***P***. ***brevicalyx*** (E.Fourn.) Ricker: ***n* = 11**: Gould (1965). ***P. confusum*** Parodi: **2*n* = 22**: Valencia & Costas (1968). ***P***. ***fimbriatum*** (Kunth) Hitchc.: ***n* = 21**: Gould (1965); Reeder (1968); **2*n* = 44**: Brown (1951: Fig. 11 and Table 1), on p. 296 apparently erroneously 2*n* = 22; Valencia & Costas (1968). ***P. hackelii*** (Arechav.) Parodi: **2*n* = 22**: Valencia & Costas (1968). ***P***. ***lasianthum*** Griseb.: **2*n* = 22**: Parodi (1946); Valencia & Costas (1968). ***P***. ***montevidense*** (Spreng.) Parodi: **2*n* = 22**: Parodi (1946); Valencia & Costas (1968). ***P***. ***napostaense*** (Speg.) Hack.: **2*n* = 22**: Covas & Bocklet (1945). ***P***. ***pringlei*** (Beal) Parodi: ***n* = 21**: Reeder (1977) as *Stipa pringlei* Scribn.; **2*n* = 42**: Myers (1947) citing an unpubl. count of R.M. Love as *S. pringlei*. ***P. ruprechtianum*** É. Desv.: **2*n* = 22**: Valencia & Costas (1968). ***P***. ***stipoides*** (Trin. & Rupr.) Hack. ex Arechav.: **2*n* = 22**: Parodi (1946); Valencia & Costas (1968) as *P. stipoides* var. *chaetophorum* (Griseb.) Parodi; Valencia & Costas (1968) as *P. stipoides* var. *genuinum* Parodi; Valencia & Costas (1968) as *P. stipoides* var. *purpurascens* (Hack.) Parodi; Valencia & Costas (1968) as *P. stipoides* var. *verruculosum* (Mez) Parodi. ***P***. ***uruguense*** Griseb.: **2*n* = 22**: Parodi (1946); Valencia & Costas (1968). ***P***. ***virescens*** (Kunth) Parodi: ***n* = ca**.**30**: Gould (1966) as *Stipa virescens* Kunth and reporting disturbance of meiosis.

***Psammochloa*** Hitchc.: **no chromosome counts available**.

***Pseudoeriocoma*** Romasch., P.M.Peterson & Soreng: **2*n* = 44**, **46**. — ***P***. ***eminens*** (Cav.) Romasch.: ***n* = 22**: Gould (1965) as *Stipa eminens* Cav.; **2*n* = 44**: Reeder (1984) as *S. eminens*; ***n* = 23**: Gould (1966) as *S. eminens*; **2*n* = 46**: Myers (1947) citing an unpubl. count of R.M. Love as *S. eminens*; Brown (1951) as *S. eminens* Cav..

***Ptilagrostiella*** Romasch., P.M.Peterson & Soreng: **2*n* = 22**. — ***P***. ***kingii*** (Bol.) Romasch.: **2*n* = 22**: Johnson (1945) as *Oryzopsis kingii* (Bol.) Beal.

***Ptilagrostis*** Griseb.: **2*n* = 22**. — ***P***. ***contracta*** Z.S.Zhang & W.L.Chen: **2*n* = 22**: Zhang & al. (2017). ***P***. ***junatovii*** Grubov: **2*n* = 22**: Krogulevich (1971, 1972). ***P***. ***mongholica*** (Turcz. ex Trin.) Griseb.: **2*n* = 22**: Sokolovskaya & Strelkova (1948); Krogulevich (1971); Murin & al. (1984) as *Achnatherum mongholicum* (Turcz. ex Trin.) Ohwi; Tateoka (1986). ***P***. ***mongholica*** subsp. ***mongholica***: **2*n* = 22**: Sokolovskaya & Probatova (1978); Probatova & Sokolovskaya (1980).

***Stipa*** L.: **2*n* = 20**, **32**, **40**, **<u>44</u>**, **48**, **66**. — ***S***. ***aktauensis*** Roshev.: **2*n* = 44**: Sokolovskaya & Probatova (1978). ***S***. ***apertifolia*** Martinovský: **2*n* = 44**: Moraldo (1986) as *S. dasyvaginata* subsp. *apenninicola* Martinovský & Moraldo. ***S***. ***arabica*** Trin. & Rupr.: ***n* = 22**: Sheidai & Attaei (2005). ***S***. ***arabica*** subsp. ***caspia*** (K.Koch) Tzvelev: **2*n* = 44**: Sokolovskaya & Probatova (1978). ***S***. ***baicalensis*** Roshev.: **2*n* = 48**: Gusik (1973); Gusik & Levkovskiy (1979); **2*n* = 44**: Probatova & al. (2006); Korobkov & al. (2013). ***S***. ***balcanica*** (Martinovský) Kožuharov: **2*n* = 44**: Kožuharov & Petrova (1991). ***S***. ***barbata*** Desf.: **2*n* = 44**: Vázquez & Devesa (1996). ***S***. ***capillacea*** Keng: **2*n* = 44**: Gupta & al. (2014: Table 1) as *S. koelzii* R.R.Stewart but *n* = 11 on p. 1245 and in Fig. 36. ***S***. ***capillata*** L.: **2*n* = 44**: Avdulov (1928, 1931: 129); Delay (1947); Sokolovskaya & Strelkova (1948); Tarnavschi (1948); Lorenzo-Andreu (1951: 201) and erroneously as “*S. ciliata* L.” on p. 202; Lorenzo-Andreu (1953); Kožuharov & Kuzmanov (1968); Petrova (1968); Skalińska & al. (1968); Májovský (1974); Chopanov & Yurtzev (1976); Sokolovskaya & Probatova (1978); Gusik & Levkovskiy (1979); Měsíček & Javůrcová-Jarolímová (1992); Vázquez & Devesa (1996); Ghukasyan (2004); Probatova & al. (2009, 2011). ***S***. ***caucasica*** Schmalh.: ***n* = 22**: Sheidai & Attaei (2005); **2*n* = 44**: Sokolovskaya & Strelkova (1939); Probatova & Seledets (2008); Sokolovskaya & Strelkova (1948) as *S. glareosa* P.A.Smirn.; Probatova & Sokolovskaya (1980) as *S. glareosa*; Erst & al. (2019) as *S. glareosa*. ***S***. ***caucasica*** subsp. ***desertorum*** (Roshev.) TzveIev; **2*n* = 44**: Sokolovskaya & Probatova (1978). ***S***. ***ehrenbergiana*** Trin. & Rupr.; ***n* = 22**: Sheidai & Attaei (2005). ***S***. ***×gegarkunii*** P.A.Smirn.: **2*n* = 44**: Ghukasyan (2004). ***S***. ***hohenackeriana*** Trin. & Rupr.: ***n* = 22**: Sheidai & Attaei (2005). ***S***. ***holosericea*** Trin.: ***n* = 22**: Sheidai & Attaei (2005); **2*n* = 44**: Chopanov & Yurtzev (1976), ascribed to subsp. *transcaucasica* (Grossh.) Tzvelev by Tzvelev (1976); Strid & Andersson (1985) as *S. fontanesii* Parl.; Strid (1987). ***S***. ***iberica*** Martinovský: **2*n* = 44**: Vázquez & Devesa (1996); Vázquez & Devesa (1996) as *S. pauneroana* (Martinovský) F.M.Vázquez & Devesa. ***S***. ***iranica*** Freitag: ***n* = 22**: Sheidai & Attaei (2005). ***S***. ***juncea*** L.: **2*n* = 44**: Myers (1947) citing an unpubl. count of R.M. Love; Nilsson & Lassen (1971); Vázquez & Devesa (1996) as *S. juncea* var. *cabanasii* F.M.Vázquez & Devesa; Vázquez & Devesa (1996) as *S. juncea* L. var. *juncea*. ***S***. ***kirghisorum*** P.A.Smirn.: **2*n* = 32**: Solntzeva (1967). ***S***. ***krylovii*** Roshev.: **2*n* = 44**: Sokolovskaya & Probatova (1978); Chepinoga & al. (2012); Probatova & al. (2013, 2016b); Chepinoga & Gnutikov (2014); Gnutikov & al. (2017). ***S***. ***lagascae*** Roem. & Schult.: ***n* = 22**: Ferchichi & al. (1994) as *S. lagascae* var. *pubescens* Hack.; **2*n* = 44**: Gould & Soderstrom (1970): Faruqi & al. (1987); Devesa & al. (1991); Vázquez & Devesa (1996) as *S. clausa* subsp. *cazorlensis* F.M.Vázquez & Devesa; Vázquez & Devesa (1996) as *S. clausa* Trab. subsp. *clausa* var. *clausa*; Vázquez & Devesa (1996) as *S. lagascae* var. *australis* Maire; Vázquez & Devesa (1996) as *S. lagascae* var. *lagascae. **S***. ***lessinigiana*** Trin. & Rupr.: ***n* = 22**: Sheidai & Attaei (2005); **2*n* = 44**: Avdulov (1931: 130); Petrova (1968); Lungeanu (1975); Chopanov & Yurtsev (1976); Magulaev (1984); Strid & Andersson (1985) as *Stipa cyllenaea* Strid; Ghukasyan (2004). ***S***. ***lingua*** Junge: **2*n* = 44**: Chopanov & Yurtsev (1976). ***S***. ***offneri*** Breistr.: **2*n* = 44**: Vázquez & Devesa (1996). ***S***. ***orientalis*** Trin. ex Ledeb.: ***n* = 10**: Singhal & al. (2014); **2*n* = 36**: Sokolovskaya & Strelkova (1939) but regarded as miscount by Probatova & Sokolovskaya (1980); **2*n* = 44**: Sokolovskaya & Probatova (1978); Agapova & al. (1993) citing an unpubl. count of O.I. Zakharyeva. ***S***. ***pennata*** L.: ***n* = 22**: Duckert-Henriod (1991) as *S. pennata* subsp. *eriocaulis* (Borbas) Martinovský & Skalický; **2*n* = 44**: Avdulov (1931: 130) as *S. joannis* Čelak.; Tarnavschi (1948) as *S. joannis*; Petrova (1968) as *S. lithophila* P.A.Smirn.; Tarnavschi & Lungeanu (1970a,b) as *S. danubialis* Dihoru & Roman; Tarnavschi & Lungeanu (1970a,b) as *S. eriocaulis* Borbás; Prokudin & al. (1977) citing an unpubl. count of O.A. Petrova; Májovský (1978) as *S. eriocaulis* subsp. *austriaca* (Beck) Martinovský; Andreev (1982) as *S. joannis*; Krasnikov (1991); Vázquez & Devesa (1996) as *S. eriocaulis*; Winterfeld & al. (2015). ***S***. ***pulcherrima*** K.Koch: **2*n* = 44**: Avdulov (1931: 130); Prokudin & al. (1977) citing an unpubl. count of O.A. Petrova; Baden (1983) as *S. epilosa* Martinovský; Strid & Andersson (1985) as *S. epilosa*; Moraldo (1986) as *S. crassiculmis* subsp. *picentina* Martinovský; **2*n* = 66**: Kožuharov & Petrova (1991) as *S. epilosa* Martinovský. ***S***. ***sareptana*** A.K.Becker: **2*n* = 44**: Titova (1935). ***S***. ***tirsa*** Steven: **2*n* = 44**: Avdulov (1931: 130) as *S. stenophylla* Czern. ex Trautv.; Prokudin & al. (1977) citing an unpubl. count of O.A. Petrova; Májovský (1978); Winterfeld & al. (2015). ***S***. ***turkestanica*** Hack.: ***n* = 22**: Sheidai & Attaei (2005). ***S***. ***turkestanica*** subsp. ***trichoides*** (P.A.Smirn.) TzveIev: **2*n* = 40**: Chopanov & Yurtsev (1976) as *S. trichoides* P. A.Smirn. ***S***. ***ucrainica*** P.A.Smirn.: **2*n* = 44**: Avdulov (1931: 130); Prokudin & al. (1977) citing an unpubl. count of O.A. Petrova. ***S***. ***zalesskii*** Wilensky ex Grossh.: **2*n* = 44**: Prokudin & al. (1977) citing an unpubl. count of O.A. Petrova.

***Stipellula*** Röser & Hamasha: **2*n* = 18**, **<u>28</u>**, **ca**. **34**, **<u>36</u>**. — ***S***. ***capensis*** (Thunb.) Röser & Hamasha: **2*n* = 18**: Borgen (1970) as *Stipa retorta* Cav.; **2*n* = ca**. **34**: Reese (1957) as *S. retorta*; ***n* = 18**, **2*n* = 36**: Devesa & al. (1991) as *S. capensis* Thunb.; **2*n* = 36**: Fernandes & Queirós (1969) as *S. capensis*; Dahlgren & al. (1971) as *S. capensis*; Gould & Soderstrom (1970) as *S. capensis*; Queirós (1974) as *S. capensis*; Vázquez & Devesa (1996) as *S. capensis*; Scholz & al. (1998) as *S. capensis. **S***. ***parviflora*** (Desf.) Röser & Hamasha: ***n* = 14**: Ferchichi & al. (1994) as *Stipa parviflora* Desf.; **2*n* = 28**: Lorenzo-Andreu (1951, 1953) as *S. parviflora*; Vázquez & Devesa (1996) as *S. parviflora*.

***Thorneochloa*** Romasch., P.M.Peterson & Soreng: **no chromosome counts available**.

***Timouria*** Roshev.: **no chromosome counts available**.

***Trikeraia*** Bor: **no chromosome counts available**.

---

**Table S1.**
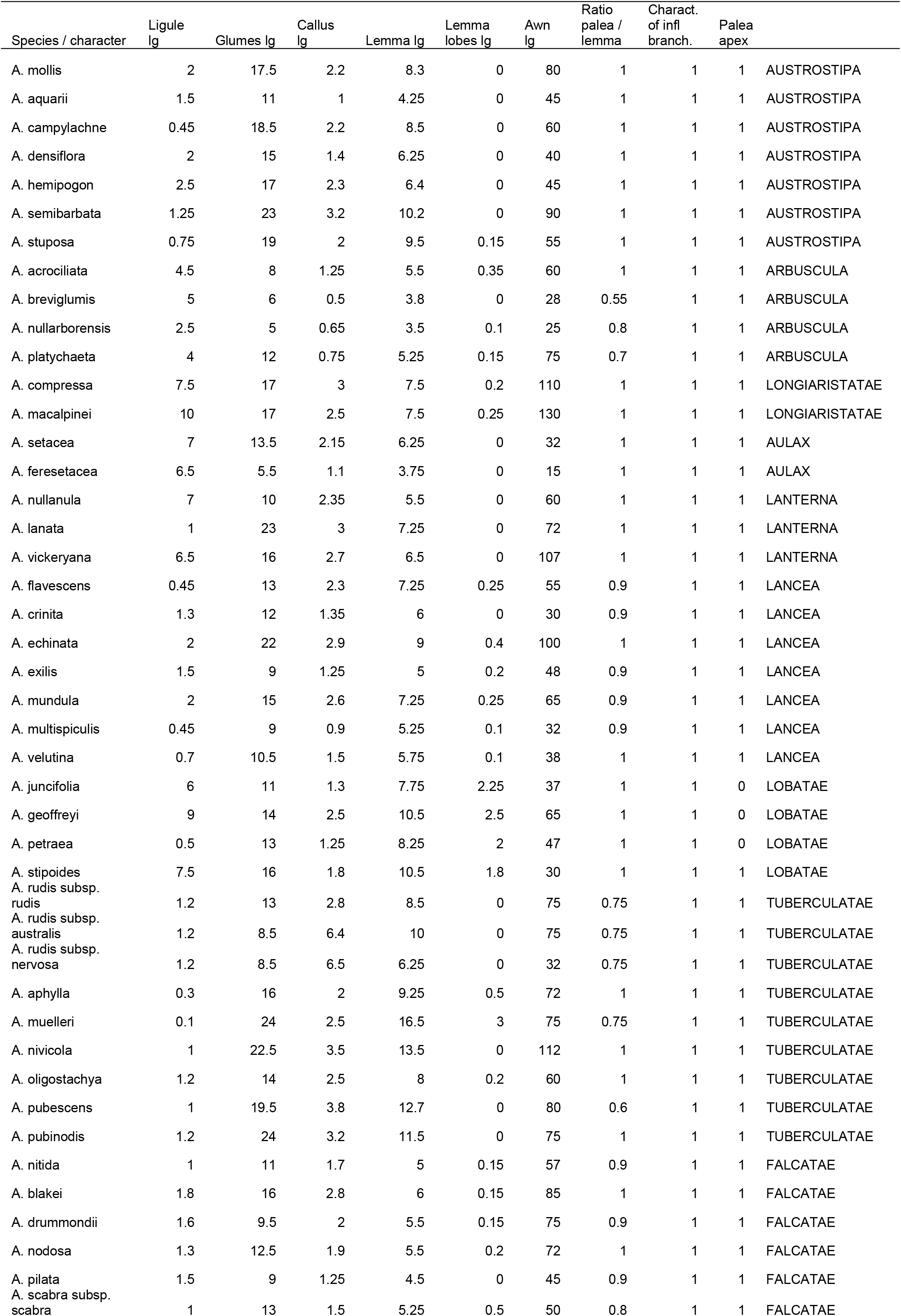
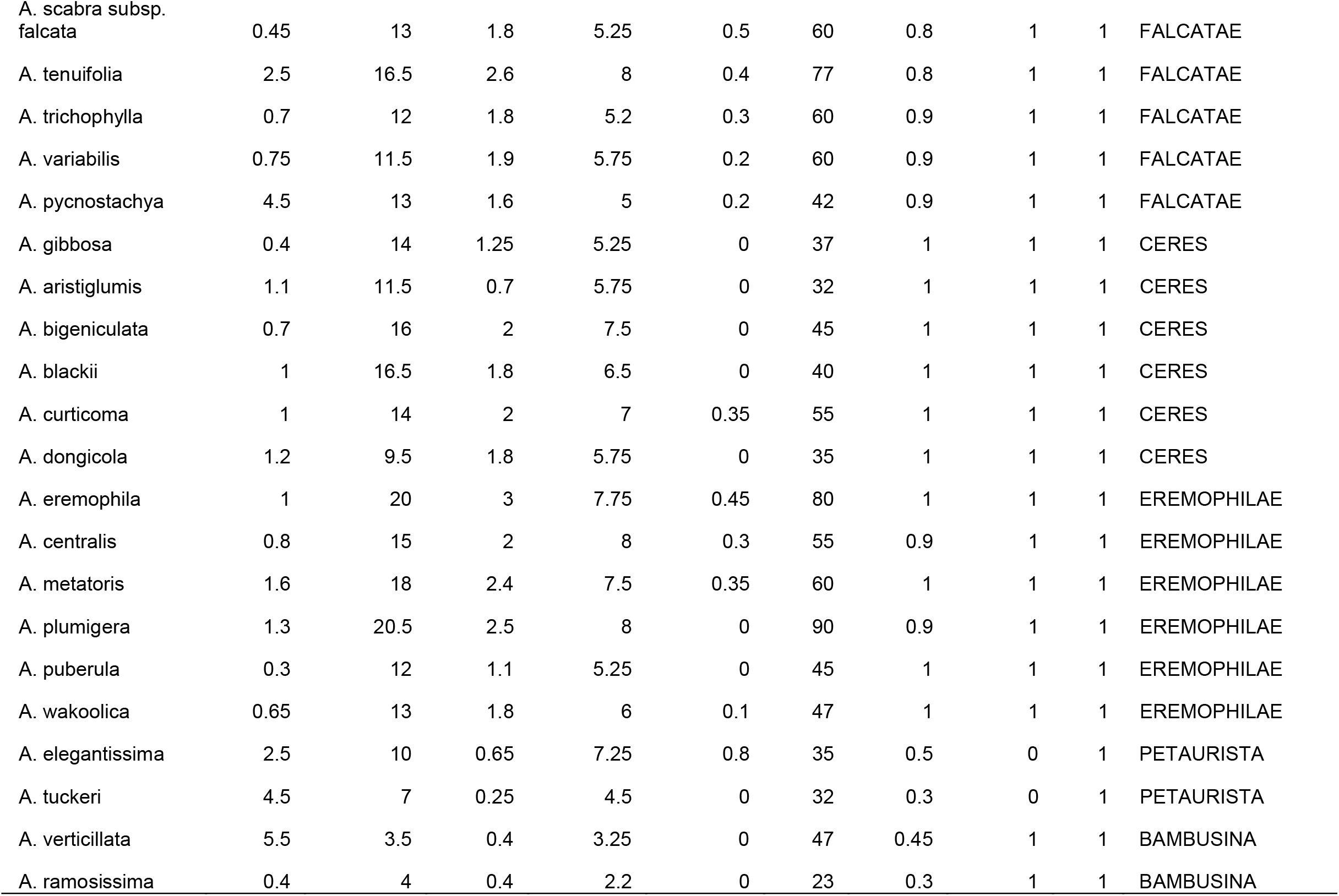
Data matrix used for macromorphological analysis.

**Table S2.** Data matrix used for micromorphological and combined macro- and micromorphological analyses.

| Species / character | Ligule g | Glumes lg | Callus lg | Lemma lg | Lemma lobes lg | Awn lg | Ratio palea / lemma | Charact. of infl. branch. | Basal cells | Hooks | Silica cells | Cork cells |  |
| --- | --- | --- | --- | --- | --- | --- | --- | --- | --- | --- | --- | --- | --- |
| A. mollis | 2 | 17.5 | 2.2 | 8.3 | 0 | 80 | 1 | 0 | 1 | 2 | 2 | 2 | AUSTROSTIPA |
| A. campylachne | 0.45 | 18.5 | 2.2 | 8.5 | 0 | 60 | 1 | 0 | 1 | 1 | 2 | 2 | AUSTROSTIPA |
| A. densiflora | 2 | 15 | 1.4 | 6.25 | 0 | 40 | 1 | 0 | 2 | 2 | 2 | 2 | AUSTROSTIPA |
| A. hemipogon | 2.5 | 17 | 2.3 | 6.4 | 0 | 45 | 1 | 0 | 2 | 2 | 1 | 2 | AUSTROSTIPA |
| A. semibarbata | 1.25 | 23 | 3.2 | 10.2 | 0 | 90 | 1 | 0 | 1 | 2 | 1 | 2 | AUSTROSTIPA |
| A. stuposa | 0.75 | 19 | 2 | 9.5 | 0.15 | 55 | 1 | 0 | 1 | 2 | 2 | 2 | AUSTROSTIPA |
| A. breviglumis | 5 | 6 | 0.5 | 3.8 | 0 | 28 | 0.55 | 0 | 3 | 2 | 1 | 2 | ARBUSCULA |
| A. acrociliata | 4.5 | 8 | 1.25 | 5.5 | 0.35 | 60 | 1 | 0 | 2 | 2 | 2 | 2 | ARBUSCULA |
| A. platychaeta | 4 | 12 | 0.75 | 5.25 | 0.15 | 75 | 0.7 | 0 | 2 | 2 | 1 | 2 | ARBUSCULA |
| A. compressa | 7.5 | 17 | 3 | 7.5 | 0.2 | 110 | 1 | 0 | 1 | 2 | 2 | 2 | LONGIARISTATAE |
| A. macalpinei | 8 | 17 | 2.5 | 7.5 | 0.25 | 130 | 1 | 0 | 1 | 2 | 2 | 2 | LONGIARISTATAE |
| A. setacea | 7 | 13.5 | 2.15 | 6.25 | 0 | 32 | 1 | 0 | 2 | 2 | 2 | 2 | AULAX |
| A. flavescens | 0.45 | 13 | 2.3 | 7.25 | 0.25 | 55 | 0.9 | 0 | 1 | 2 | 1 | 2 | LANCEA |
| A. mundula | 2 | 15 | 2.6 | 7.25 | 0.25 | 65 | 0.9 | 0 | 2 | 2 | 2 | 2 | LANCEA |
| A. stipoides | 7.5 | 16 | 1.8 | 10.5 | 1.8 | 30 | 1 | 0 | 3 | 2 | 1 | 1 | LOBATAE |
| A. rudis subsp. rudis | 1.2 | 13 | 2.8 | 8.5 | 0 | 75 | 0.75 | 0 | 2 | 2 | 2 | 2 | TUBERCULATAE |
| A. rudis subsp. australis | 1.2 | 8.5 | 6.4 | 10 | 0 | 75 | 0.75 | 0 | 2 | 2 | 2 | 2 | TUBERCULATAE |
| A. muelleri | 0.1 | 24 | 2.5 | 16.5 | 3 | 75 | 0.75 | 0 | 2 | 2 | 1 | 2 | TUBERCULATAE |
| A. pubescens | 1 | 19.5 | 3.8 | 12.7 | 0 | 80 | 0.6 | 0 | 1 | 1 | 2 | 2 | TUBERCULATAE |
| A. pubinodis | 1.2 | 24 | 3.2 | 11.5 | 0 | 75 | 1 | 0 | 2 | 2 | 1 | 2 | TUBERCULATAE |
| A. nitida | 1 | 11 | 1.7 | 5 | 0.15 | 57 | 0.9 | 0 | 1 | 2 | 2 | 2 | FALCATAE |
| A. drummondii | 1.6 | 9.5 | 2 | 5.5 | 0.15 | 75 | 0.9 | 0 | 1 | 2 | 2 | 2 | FALCATAE |
| A. nodosa | 1.3 | 12.5 | 1.9 | 5.5 | 0.2 | 72 | 1 | 0 | 1 | 2 | 2 | 2 | FALCATAE |
| A. scabra subsp. scabra | 1 | 13 | 1.5 | 5.25 | 0.5 | 50 | 0.8 | 0 | 1 | 2 | 2 | 2 | FALCATAE |
| A. scabra subsp. falcata | 0.45 | 13 | 1.8 | 5.25 | 0.5 | 60 | 0.8 | 0 | 1 | 2 | 2 | 2 | FALCATAE |
| A. variabilis | 0.75 | 11.5 | 1.9 | 5.75 | 0.2 | 60 | 0.9 | 0 | 1 | 2 | 2 | 2 | FALCATAE |
| A. gibbosa | 0.4 | 14 | 1.25 | 5.25 | 0 | 37 | 1 | 0 | 2 | 2 | 2 | 2 | CERES |
| A. aristiglumis | 1.1 | 11.5 | 0.7 | 5.75 | 0 | 32 | 1 | 0 | 2 | 2 | 2 | 2 | CERES |
| A. bigeniculata | 0.7 | 16 | 2 | 7.5 | 0 | 45 | 1 | 0 | 2 | 2 | 2 | 2 | CERES |
| A. eremophila | 1 | 20 | 3 | 7.75 | 0.45 | 80 | 1 | 0 | 1 | 2 | 2 | 2 | EREMOPHILAE |
| A. elegantissima | 2.5 | 10 | 0.65 | 7.25 | 0.8 | 35 | 0.5 | 1 | 3 | 1 | 2 | 2 | PETAURISTA |
| A. tuckeri | 4.5 | 7 | 0.25 | 4.5 | 0 | 32 | 0.3 | 1 | 3 | 1 | 2 | 2 | PETAURISTA |
| A. verticillata | 5.5 | 3.5 | 0.4 | 3.25 | 0 | 47 | 0.45 | 0 | 2 | 1 | 2 | 2 | BAMBUSINA |
| A. ramosissima | 0.4 | 4 | 0.4 | 2.2 | 0 | 23 | 0.3 | 0 | 3 | 1 | 2 | 2 | BAMBUSINA |

## REFERENCES

Acedo, C. & Llamas, F. 2001. Variation of micromorphological characters of lemma and palea in the genus *Bromus* (Poaceae). Ann. Bot. Fenn. 38: 1–14. https://www.jstor.org/stable/23726828

Agapova, N.D., Arkharova, K.B., Vakhtina, E.A., Zemskova, E.A. & Tarvis, L.V. 1993. Chisla khromosom tsvetkovykh rasteniy flory SSSR: Moraceae–Zygophyllaceae [Chromosome numbers in flowering plants of the flora of the USSR: Moraceae– Zygophyllaceae]. Nauka, St. Petersburg.

Álvarez, I. & Wendel, J.F. 2003. Ribosomal ITS sequences and plant phylogenetic inference. Molec. Phylogen. Evol. 29: 417–434. https://doi.org/10.1016/S1055-7903(03)00208-2

Avdulov, N. 1931. Kario-sistematicheskoe issledovanie semeistva zlakov. [Karyosystematic study of the family of grasses.] Trudy Prikl. Bot. Selekts., Suppl. 44: 1–428.

Bailey, C.D., Carr, T.G., Harris, S.A. & Hughes, C.F. 2003. Characterization of angiosperm nrDNA: polymorphism, paralogy, and pseudogenes. Molec. Phylogen. Evol. 29: 435–455. https://doi.org/10.1016/j.ympev.2003.08.021

Baldwin, B.G., Sanderson, M.J., Porter, J.M., Wojciechowski, M.F., Campbell, C.S. & Donoghue, M.J. 1995. The ITS region of nuclear ribosomal DNA: a valuable source of evidence on angiosperm phylogeny. Ann. Missouri Bot. Gard. 82: 247–277. doi: 10.2307/2399880

Barber, J.C., Hames, K.A., Cialdella, A.M., Giussani, L.M. & Morrone, O. 2009. Phylogenetic relationships of *Piptochaetium* Presl (Poaceae: Stipeae) and related genera as reconstructed from nuclear and chloroplast datasets. Taxon 58: 375–380. https://doi.org/10.1002/tax.582005

Barkworth, M.E. 1983. *Ptilagrostis* in North America and its relationship to other Stipeae. Syst. Bot. 8(4): 395–419. https://doi.org/10.2307/2418359

Barkworth, M.E. 1990. *Nassella* (Gramineae: Stipeae): revised interpretation and nomenclatural changes. Taxon 39: 597–614. https://doi.org/10.2307/1223366

Barkworth, M.E.1993. North American Stipeae (Gramineae): taxonomic changes and other comments. Phytologia 74(1): 1–25. https://www.biodiversitylibrary.org/item/47126#page/3/mode/thumb

Barkworth, M.E. 2007. Stipeae Dumort. Pp. 109–186 in: Barkworth, M.E., Capels, K.M., Long, S., Anderton, L.K. & Piep, M.B. (eds.), Flora of North America north of Mexico, vol. 24, Magnoliophyta: Commelinidae (in part): Poaceae, part 1. New York: Oxford University Press.

Barkworth, M.E. & Everett, J. 1987. Evolution in the Stipeae: Identification and relationships of its monophyletic taxa. Pp. 251–264 in: Soderstrom, T.R., Hilu, K.W., Campbell, C.S. & Barkworth, M.E. (eds.), Grass systematics and evolution. Washington, DC: Smithsonian Institution Press.

Barkworth, M.E. & Torres, M.A. 2001. Distribution and diagnostic characters of *Nassella* (Poaceae: Stipeae). Taxon 50(2): 439−468. https://doi.org/10.2307/1223891

Barkworth, M.E., Arriaga, M.O., Smith, J.F., Jacobs, S.W.L., Valdés-Reyna, J. & Bushman, B.S. 2008. Molecules and morphology in South American Stipeae (Poaceae). Syst. Bot. 33(4): 719–731. https://doi.org/10.1600/036364408786500235

Bayly, M.J. & Ladiges, P.Y. 2007. Divergent paralogues of ribosomal DNA in eucalypts (Myrtaceae). Molec. Phylogen. Evol. 44: 346–356. https://doi.org/10.1016/j.ympev.2006.10.027

Blaner, A., Schneider, J. & Röser, M. 2014. Phylogenetic relationships in the grass family (Poaceae) based on the nuclear single copy locus topoisomerase 6 compared to chloroplast DNA. Syst. Biodivers. 12: 111–124. https://doi.org/10.1080/14772000.2014.890137

Blattner, F.R. 1999. Direct amplification of the entire ITS region from poorly preserved plant material using recombinant PCR. BioTechniques. 27: 1180–1186. https://doi.org/10.2144/99276st04

Borgen, L. 1970. Chromosome numbers of Macaronesian flowering plants. Nytt Mag. Bot. 17: 145–161.

Brassac, J., Jakob, S.S. & Blattner, F.R. 2012. Progenitor-derivative relationships of *Hordeum* polyploids (Poaceae, Triticeae) inferred from sequences of TOPO6, a nuclear low-copy gene region. PLoS ONE 7: e333808.

Buckler, E.S., Ippolito, A. & Holtsford, T.P. 1997. The evolution of ribosomal DNA: divergent paralogues and phylogenetic implications. Genetics 145(3): 821–832.

Bustam, B.M. 2010. Systematic studies of Australian stipoid grasses (*Austrostipa*) based on micro-morphological and molecular characteristics. Biodiversitas 11(1): 9–14.

Catalán, P., Kellogg, E.A. & Olmstead, R.G. 1997. Phylogeny of Poaceae subfamily Pooideae based on chloroplast ndhF gene sequences. Molec. Phylogen. Evol. 8: 150–166. https://doi.org/10.1006/mpev.1997.0416

Chase, M.W. & Hills, H.H. 1991. Silica gel: an ideal material for field preservation of leaf samples for DNA studies. Taxon 40(2): 215−220. https://doi.org/10.2307/1222975

Cialdella, A.M., Guissani, L.M., Aagesen, L., Zuloaga, F.O. & Morrone, O. 2007. A phylogeny of *Piptochaetium* (Poaceae: Pooideae: Stipeae) and related genera based on a combined analysis including *trnL–F, rpl16*, and morphology. Syst. Bot. 32: 545–559. https://doi.org/10.1600/036364407782250607

Cialdella, A.M., Salariato, D.L., Aagesen, L., Giussani, L.M., Zuloaga, F.O. & Morrone, O. 2010. Phylogeny of New World Stipeae (Poaceae): an evaluation of the monophyly of *Aciachne* and *Amelichloa*. Cladistics 26: 563–578. doi: 10.1111/j.1096-0031.2010.00310.x

Cialdella, A.M., Sede, S.M., Romaschenko, K., Peterson, P.M., Soreng, R.J., Zuloaga, F.O. & Morrone, O. 2014. Phylogeny of *Nassella* (Stipeae, Pooideae, Poaceae) based on analyses of chloroplast and nuclear ribosomal DNA and morphology. Syst. Bot. 39(3): 814–828. https://doi.org/10.1600/036364414X681419

Clark, J.W. & Donoghue, P.C.J. 2018. Whole-genome duplication and plant macroevolution. Trends Pl. Sci. 23(10): 933–945. https://doi.org/10.1016/j.tplants.2018.07.006.

Clayton, W.D. 1970. Gramineae (part 1). Pp. 1–176 in: Milne-Redhead, E. & Polhill, R.M. (eds.), Flora of tropical East Africa. London: Crown Agents for Oversea Governments and Administrations.

Clayton, W.D. 1972. Gramineae. Pp. 349–512 in: Hepper, F.N. (ed.), Flora of east tropical Africa, vol. 3(2). London: Crown Agents for Oversea Governments and Administrations.

Curto, M. & Henderson, D.M. 1998. A new *Stipa* (Poaceae: Stipeae) from Idaho and Nevada. Madroño 45(1): 57–63. https://www.jstor.org/stable/41425241

Darlington, C.D. & Wylie, A.P. 1956. Chromosome atlas of flowering plants. Chromosome atlas of flowering plants. London: George Allen and Unwin Ltd.

Davis, J.I. & Soreng, R.J. 2007. A preliminary phylogenetic analysis of the grass subfamily Pooideae subfamily Pooideae (Poaceae), with attention to structural features of the plastid and nuclear genomes, including an intron loss in GBSSI. Aliso 23, 335–348. doi: 10.5642/aliso.20072301.27

Dawson, M.I. & Beuzenberg, E.J. 2000. Contributions to a chromosome atlas of the New Zealand flora. New Zealand J. Bot. 38: 1–23. https://doi.org/10.1080/0028825X.2000.9512671

Decker, H.F. 1964. Affinities of the grass genus *Ampelodesmos*. Brittonia 16: 76–79. https://link.springer.com/content/pdf/10.2307%2F2805186.pdf

Döring, E., Schneider, J., Hilu, K.W. & Röser, M. 2007. Phylogenetic relationships in the Aveneae/Poeae complex (Pooideae, Poaceae). Kew Bull. 62(3): 407–424. https://www.jstor.org/stable/20443367

Edgar, E. & Connor, H.E. 2000. Flora of New Zealand, vol. 5. Gramineae. Lincoln, New Zealand: Manaaki Whenua Press.

Everett, J. & Jacobs, S.W.L. 1990. Notes on *Stipa* (Poaceae) in Australia and Easter Island. Telopea 4: 7–11. dx.doi.org/10.7751/telopea19904912

Everett, J., Jacobs, S.W.L. & Nairn, L. 2009. Tribe Stipeae. Pp. 11–70 in: Wilson, A. (ed.), Flora of Australia, vol. 44A, Poaceae 2. (ABRS: Canberra; CSIRO Publishing: Melbourne).

Fan, X., Zhang, H., Sha, L., Zhang, L., Yang, R., Ding, C. & Zhou, Y. 2007. Phylogenetic analysis among *Hystrix*, *Leymus* and its affinitive genera (Poaceae: Triticeae) based on the sequences of a gene encoding plastid acetyl-CoA carboxylase. Plant Sci. 172: 701– 707. doi: 10.1016/j.plantsci.2006.11.012

Fan, X., Sha, L.-N., Yang, R.-W., Zhang, H.-Q., Kang, H.-Y., Ding, C.-B., Zhang, L., Zheng, Y.-H. & Zhou, Y.-H. 2009. Phylogeny and evolutionary history of *Leymus* (Triticeae; Poaceae) based on a single-copy nuclear gene encoding plastid acetyl-CoA carboxylase. BMC Evol. Biol. 9: 247. doi: 10.1186/1471-2148-9-24

Fedorov, A.A. 1969. Chromosome numbers of flowering plants. Leningrad, U.S.S.R: Izdatelstvo Nauka.

Fish, L., Mashau, A.C., Moeaha, M.J. & Nembudani, M.T. 2015. Identification guide to southern African grasses. An identification manual with keys, descriptions and distributions. Strelitzia 36. Pretoria: South African National Biodiversity Institute.

Freitag, H. 1975. The genus *Piptatherum* (Gramineae) in southwest and south Asia. Notes Roy. Bot. Gard. Edinburgh 33: 341–408.

Freitag, H. 1985. The genus *Stipa* (Gramineae) in southwest and south Asia. Notes Roy. Bot. Gard. Edinburgh 42: 355–489.

Freitag, H. 1989. Piptatherum and Stipa (Gramineae) in the Arabian Peninsula and tropical East Africa. Pp. 115–132 in: Tan, K. (ed.), The Davis and Hedge festschrift. Edinburgh: Edinburgh University Press.

GPWG (Grass Phylogeny Working Group). 2001. Phylogeny and subfamilial classification of the grasses (Poaceae). Ann. Missouri Bot. Gard. 88: 373–457. https://doi.org/10.2307/3298585

Ghukasyan, A.G. 2004. Kariologisheskaya izuchennost zlakov (Poaceae) Armenii. [Extent of karyological study of Armenian grasses (Poaceae).] *Fl. Rastitel’nost’* Rastitel’nye Resursy Armenii [Flora, vegetation and plant resources of Armenia] 15: –8. http://takhtajania.asj-oa.am/id/eprint/297

Hamasha, H., von Hagen, K.B. & Röser, M. 2012. *Stipa* (Poaceae) and allies in the Old World: molecular phylogenetics realigns genus circumscription and gives evidence on the origin of American and Australian lineages. Pl. Syst. Evol. 298(2), 351–367. doi: 10.1007/s00606-011-0549-5

Hammer, O., Harper, D.A.T. & Ryan, P.D. 2001. PAST: paleontological statistic software package for education and data analysis. *Palaeontol*. Electron. 4: 1–9.

Hand, M.L., Cogan, N.O., Stewart, A.V. & Forster, J.W. 2010. Evolutionary history of tall fescue morphotypes inferred from molecular phylogenetics of the *Lolium*-*Festuca* species complex. BMC Evol. Biol. 10, 303. https://doi.org/10.1186/1471-2148-10-303

Hilu, K.W., Alice, L.A. & Liang, H. 1999. Phylogeny of the Poaceae inferred from *mat*K sequences. Ann. Missouri Bot. Gard. 86(4): 835–851. doi: 10.2307/2666171

Hochbach, A., Schneider, J. & Röser, M. 2015. A multi-locus analysis of phylogenetic relationships within grass subfamily Pooideae (Poaceae) inferred from sequences of nuclear single copy gene regions compared with plastid DNA. Molec. Phylogen. Evol. 87: 14–27. https://doi.org/10.1016/j.ympev.2015.03.010

Hochbach, A., Linder, P.H. & Röser, M. 2018. Nuclear genes, *mat*K and the phylogeny of the Poales. Taxon 67(3): 521–536. https://doi.org/10.12705/673.5

Hsiao, C., Jacobs, S.W.L., Chatterton, N.J. & Asay, K.H. 1999. A molecular phylogeny of the grass family (Poaceae) based on the sequences of nuclear ribosomal DNA (ITS). *Austral*. Syst. Bot. 11(5–6): 667–688. https://doi.org/10.1071/SB97012

Huang, S., Sirikhachornkit, A., Su, X., Faris, J., Gill, B., Haselkorn, R. & Gornicki, P. 2002. Genes encoding plastid acetyl-CoA carboxylase and 3-phosphoglycerate kinase of the *Triticum*/*Aegilops* complex and the evolutionary history of polyploid wheat. Proc. Natl. Acad. Sci. USA 99: 8133–8138. doi: 10.1073/pnas.072223799

Jacobs, S.W.L., Everett, J., Connor, H.E. & Edgar, E. 1989. Stipoid grasses in New Zealand. New Zealand J. Bot. 27(4): 569–582. https://doi.org/10.1080/0028825X.1989.10414140

Jacobs, S.W.L. & Everett, J. 1996. *Austrostipa*, a new genus, and new names for Australasian species formerly included in *Stipa* (Gramineae). Telopea 6: 579–595. https://doi.org/10.7751/telopea19963026

Jacobs, S.W.L., Everett, J., Barkworth, M.E. & Hsiao, C. 2000. Relationships within the stipoid grasses (Gramineae). Pp. 75–82 in: Jacobs, S.W.L. & Everett, J. (eds.), Grasses, systematics and evolution. Melbourne, Victoria: CSIRO Publishing.

Jacobs, S., Bayer, R., Everett, J., Arriaga, M., Barkworth, M., Sabin-Badereau, A., Torres, A., Vázquez, F. & Bagnall, N. 2007. Systematics of the tribe Stipeae (Gramineae) using molecular data. Aliso 23: 349–361. doi: 10.5642/aliso.20072301.28

Johnson, L.A. & Soltis, D.E. 1994. *matK* DNA sequences and phylogenetic reconstruction in Saxifragaceae s.str. Syst. Bot. 19(1): 143–156. doi: 10.2307/2419718

Kearse, M., Moir, R., Wilson, A., Stones-Havas, S., Cheung, M., Sturrock, S., Buxton, S., Cooper, A., Markowitz, S., Duran, C., Thierer, T., Ashton, B., Mentjies, P. & Drummond, A. 2012. Geneious Basic: an integrated and extendable desktop software platform for the organization and analysis of sequence data. Bioinformatics 28: 1647– 1649. https://doi.org/10.1093/bioinformatics/bts199

Kožuharov, S.I. & Petrova, A.V. 1991. Chromosome numbers of Bulgarian angiosperms. Fitologija 39: 72–77.

Krawczyk, K., Nobis, M., Nowak, A., Szczecińska, M. & Sawicki, J. 2017. Phylogenetic implications of nuclear rRNA IGS variation in *Stipa* L. (Poaceae). Sci. Rep. 7: 11506. https://doi.org/10.1038/s41598-017-11804-x

Krawczyk, K., Nobis, M., Myszczyński, K., Klichowska, E. & Sawicki, J. 2018. Plastid super-barcodes as a tool for species discrimination in feather grasses (Poaceae: *Stipa*). Sci. Rep. 8: 1924. https://doi.org/10.1038/s41598-018-20399-w

Larkin, M.A., Blackshields, G., Brown, N.P., Chenna, R., McGettigan, P.A., McWilliam, H., Valentin, F., Wallace, I.M., Wilm, A., Lopez, R., Thompson, J.D., Gibson, T.J. & Higgins, D.G. 2007. ClustalW and ClustalX version 2.0. Bioinformatics 23: 2947–2948. https://doi.org/10.1093/bioinformatics/btm404

Liang, H. & Hilu, K.W. 1996. Application of the *matK* gene sequences to grass systematics. Canad. J. Bot. 74(1): 125–134. https://doi.org/10.1139/b96-017

Mathews, S., Tsai, R.C. & Kellogg, E.A. 2000. Phylogenetic structure in the grass family (Poaceae): evidence from the nuclear gene phytochrome B. Amer. J. Bot. 87(1): 96– 107. https://doi.org/10.2307/2656688

Mehra, P.N. & Sharma, M.L. 1975. Pp. 501–502 in: Löve, Á. (ed.), IOPB chromosome number reports XLIX. Taxon 24(4): 501–516. https://doi.org/10.1002/j.1996-8175.1975.tb00341.x

Mehra, P.N. & Sharma, M.L. 1977. Cytological studies on some grasses of Kashmir. Cytologia 42: 111–123. https://doi.org/10.1508/cytologia.42.111

Mejía Saulés, T. & Bisby, F.A. 2003. Silica bodies and hooked papillae in lemmas of *Melica* species (Gramineae: Pooideae). Bot. J. Linn. Soc. 141(4): 447–463. https://doi.org/10.1046/j.1095-8339.2003.00152.x

Murray, B.G., de Lange, P.J. & Ferguson, A.R. 2005. Nuclear DNA variation chromosome numbers and polyploidy in the endemic and indigenous grass flora of New Zealand. Ann. Bot. (Oxford) 96: 1293–1305. https://doi.org/10.1093/aob/mci281

Myers, W.M. 1947. Cytology and genetics of forage grasses. Bot. Rev. (Lancaster) 13(6–7): 319–422. https://www.jstor.org/stable/4353363 and https://www.jstor.org/stable/4353364

Nieto Feliner, G. & Rosselló, J.A. 2007. Better the devil you know? Guidelines for insightful utilization of nrDNA ITS in species-level evolutionary studies in plants. Molec. Phylogen. Evol. 44(2): 911–919. https://doi.org/10.1016/j.ympev.2007.01.013

Nilsson, Ö. & Lassen, P. 1971. Chromosome numbers of vascular plants from Austria, Mallorca and Yugoslavia. Bot. Not. 124: 270–276.

Nobis, M. 2013. Taxonomic revision of the *Stipa lipskyi* group (Poaceae: *Stipa* section *Smirnovia*) in the Pamir Alai and Tian-Shan Mountains. Pl. Syst. Evol. 299: 1307– 1354. https://doi.org/10.1007/s00606-013-0799-5

Nobis, M., Gudkova, P.D., Baiakhmetov, E., Żabicka, J., Krawczyk, K. & Sawicki, J. 2019a. Hybridisation, introgression events and cryptic speciation in *Stipa* (Poaceae): a case study of the *Stipa heptapotamica* hybrid-complex. Perspect. Pl. Ecol. Evol. Syst. 39: 125457. https://doi.org/10.1016/j.ppees.2019.05.001

Nobis, M., Gudkova, P.D. & Nowak, A. 2019b. *Neotrinia* gen. nov. and *Pennatherum* sect. nov. in *Achnatherum* (Poaceae: Stipeae). Turczaninowia 22(1): 37–41. https://doi.org/10.14258/turczaninowia.22.1.5

Nobis, M., Gudkova, P.D. & Pendry, C. 2019c. Synopsis of the tribe Stipeae (Poaceae) in Nepal. PhytoKeys 128: 97–119. https://doi.org/10.3897/phytokeys.128.34637

Nobis, M., Gudkova, P.D., Nowak, A., Sawicki, J. & Nobis, A. 2020. A synopsis of the genus *Stipa* (Poaceae) in Middle Asia, including a key to species identification, an annotated checklist, and phytogeographic analyses. Ann. Missouri Bot. Gard. 105(1): 1–63. https://doi.org/10.3417/2019378

Olonova, M.V., Barkworth, M.E. & Gudkova, P.D. 2016. Lemma micromorphology and the systematics of Siberian species of *Stipa* (Poaceae). Nordic J. Bot. 34(3): 322–334. https://doi.org/10.1111/njb.00881

Ortúñez, E. & de la Fuente, V. 2010. Epidermal micromorphology of the genus *Festuca* L. (Poaceae) in the Iberian Peninsula. Pl. Syst. Evol. 284: 201–218. https://doi.org/10.1007/s00606-009-0248-7

Peñailillo, P. 2002. El género *Jarava* Ruiz et Pav. (Stipeae-Poaceae): delimitación y nuevas combinaciones. *Gayana*, Bot 59: 27−34.

Peñailillo, P. 2003. Jarava Ruiz et Pav. In: Soreng, R.J., Peterson, P.M., Davidse, G., Judziewicz, E.J., Zuloaga, F.O., Filgueiras, T.S. & Morrone, O. (eds.), Catalogue of New World grasses (Poaceae): IV. subfamily Pooideae. Contr. U.S. Natl. Herb. 48: 402−409.

Peterson, P.M., Romaschenko, K., Soreng, R.J. & Valdés Reyna, J. 2019. A key to the North American genera of Stipeae (Poaceae, Pooideae) with descriptions and taxonomic names for species of *Eriocoma*, *Neotrinia*, *Oloptum*, and five new genera: *Barkworthia*, ×*Eriosella*, *Pseudoeriocoma*, *Ptilagrostiella*, and *Thorneochloa*. PhytoKeys 126: 89−125. https://doi.org/10.3897/phytokeys.126.34096

Petrova, O.A. 1968. Khromosomnyy sostav nekotorykh zlakov flory Ukrainy v svyasi s usloviami ikh proizrastaniya. [Chromosomal composition of some Ukrainian grasses according to their growing conditions.] Pp. 37–39 in: Nikitin, V.N. (ed.), Biologicheskaya nauka v universitetakh I pedagogicheskikh institutakh Ukrainy za 50 let. [Biological science in universities and pedagogical institutes of Ukraine for 50 years]. Charkov: Charkov University.

Prokudin, Yu.N., Vovk, A.G, Petrova, O.A, Ermolenko, E.D. & Vernichenko, Yu.V. (eds.) 1977. Zlaki Ukrainy. [Grasses of Ukraine.] Kiev: Naukova Dumka.

Razafimandimbison, S.G., Kellogg, E.A. & Bremer, B. 2004. Recent origin and phylogenetic utility of divergent ITS putative pseudogenes: a case study from the Naucleeae (Rubiaceae). Syst. Biol. 53(2): 177–192. https://doi.org/10.1080/10635150490423278

Romaschenko, K., Peterson, P.M., Soreng, R.J., Garcia-Jacas, N., Futoma, O. & Susanna, A. 2008. Molecular phylogenetic analysis of the American Stipeae (Poaceae) resolves *Jarava* sensu lato polyphyletic: evidence for a new genus, *Pappostipa*. J. Bot. Res. Inst. Texas 2(1): 165–192. https://www.jstor.org/stable/41971613

Romaschenko, K., Peterson, P.M., Soreng, R.J., Garcia-Jacas, N. & Susanna, A. 2010. Phylogenetics of Stipeae (Poaceae, Pooideae) based on plastid and nuclear DNA sequences. Pp. 511–537 in: Seberg, O., Petersen, G., Barfod, A.F. & Davis, J.I. (eds.), Diversity, phylogeny, and evolution in the monocotyledons. Aarhus: Aarhus University Press.

Romaschenko, K., Peterson, P.M., Soreng, R.J., Futorna, O. & Susanna, A. 2011. Phylogenetics of *Piptatherum* s.l. (Poaceae: Stipeae): evidence for a new genus, *Piptatheropsis*, and resurrection of *Patis*. Taxon 60(6): 1703–1716. https://doi.org/10.1002/tax.606015

Romaschenko, K., Peterson, P.M., Soreng, R.J., Garcia-Jacas, N., Futorna, O. & Susanna, A. 2012. Systematics and evolution of the needle grasses (Poaceae: Pooideae: Stipeae) based on analysis of multiple chloroplast loci, ITS, and lemma micromorphology. Taxon 61(1): 18–44. https://doi.org/10.1002/tax.611002

Romaschenko, K., Garcia-Jacas, N., Peterson, P.M., Soreng, R.J., Vilatersana, R. & Susanna, A. 2013. Miocene–Pliocene speciation, introgression, and migration of *Patis* and *Ptilagrostis* (Poaceae: Stipeae). Molec. Phylogen. Evol. 70(2014): 244–259. https://doi.org/10.1016/j.ympev.2013.09.018

Romaschenko, K., Garcia-Jacas, N., Peterson, P.M., Soreng, R.J., Vilatersana, R. & Susanna, A. 2014. Miocene–Pliocene speciation, introgression, and migration of *Patis* and *Ptilagrostis* (Poaceae: Stipeae). Molec. Phylogen. Evol. 70: 244–259. https://doi.org/10.1016/j.ympev.2013.09.018

Saarela, J.M., Wysocki, W.P., Barrett, C.F., Soreng, R.J., Davis, J.I., Clark, L.G., Kelchner, S.A., Pires, J.C., Edger, P.P., Mayfield, D.R. & Duvall, M.R. 2015. Plastid phylogenomics of the coolseason grass subfamily: clarification of relationships among early-diverging tribes. AoB Plants 7: plv046. doi: 0.1093/aobpla/plv046.

Saarela, J.M., Burke, S.V., Wysocki, W.P., Barrett, M.D., Clark, L.G., Craine, J.M., Peterson, P.M., Soreng, R.J., Vorontsova, M.S. & Duvall, M.R. 2018. A 250 plastome phylogeny of the grass family (Poaceae): topological support under different data partitions. PeerJ. 6: e4299. doi: 10.7717/peerj.4299.

Schneider, J., Döring, E., Hilu, K.W. & Röser, M. 2009. Phylogenetic structure of the grass subfamily Pooideae based on comparison of plastid *matK* gene–3’*trnK* exon and nuclear ITS sequences. Taxon 58: 405–424. https://doi.org/10.1002/tax.582008

Schneider, J., Winterfeld, G., Hoffmann, M.H. & Röser, M. 2011. Duthieeae, a new tribe of grasses (Poaceae) identified among the early diverging lineages of subfamily Pooideae: molecular phylogenetics, morphological delineation, cytogenetics, and biogeography. Syst. Biodivers. 9: 27–44. https://doi.org/10.1080/14772000.2010.544339

Schneider, J., Winterfeld, G. & Röser, M. 2012. Polyphyly of the grass tribe Hainardieae (Poaceae: Pooideae): identification of its different lineages based on molecular phylogenetics, including morphological and cytogenetic characteristics. Organisms Diversity Evol. 12: 113–132. https://doi.org/10.1007/s13127-012-0077-3

Sclovich, S.E., Giussani, L.M., Cialdella, A.M. & Sede, S.M. 2015. Phylogenetic analysis of *Jarava* (Poaceae, Pooideae, Stipeae) and related genera: testing the value of the awn indumentum in the circumscription of Jarava. Pl. Syst. Evol. 301: 1625–1641. https://doi.org/10.1007/s00606-014-1175-9

Sha, L., Fan, X., Yang, R., Kang, H., Ding, C., Zhang, L., Zheng, L. & Zhou, Y. 2010. Phylogenetic relationships between *Hystrix* and its closely related genera (Triticeae; Poaceae) based on nuclear *Acc1*, *DMC1* and chloroplast *trnL–F* sequences. Molec. Phylogen. Evol. 54: 327–335. https://doi.org/10.1016/j.ympev.2009.05.005.

Snow, N. 1996. The phylogenetic utility of lemmatal micromorphology in *Leptochloa* s.l. and related genera in subtribe Eleusininae (Poaceae, Chloridoideae, Eragrostideae). Ann. Missouri Bot. Gard. 83: 504–529. doi: 10.2307/2399991

Sokal, R.R. & Sneath, P.H. 1963. Principles of numerical taxonomy. San Francisco: W.H. Freeman.

Soreng, R.J. & Davis, J.I. 2000. Phylogenetic structure in Poaceae subfamily Pooideae as inferred from molecular and morphological characters: misclassification versus reticulation. Pp. 61–74 in: Jacobs, S.W.L. & Everett, J. (eds.), Grasses: systematics and evolution. Melbourne: CSIRO Publishing.

Soreng, R.J., Peterson, P.M., Davidse, G., Judziewicz, E.J., Zuloaga, F.O., Filgueiras, T.S. & Morrone, O. 2003. Catalogue of New World grasses (Poaceae): IV. subfamily Pooideae. Contr. U.S. Natl. Herb. 48: 1–730.

Soreng, R.J., Davis, J.I. & Voionmaa, M.A. 2007. A phylogenetic analysis of Poaceae tribe Poeae sensu lato based on morphological characters and sequence data from three plastid-encoded genes: evidence for reticulation, and a new classification of the tribe. Kew Bull. 62(3): 425–454. https://www.jstor.org/stable/20443368

Soreng, R.J., Peterson, P.M., Romaschenko, K., Davidse, G., Teisher, J.K., Clark, L.G., Barberá, P., Gillespie, L.J. & Zuloaga, F.O. 2017. A worldwide phylogenetic classification of the Poaceae (Gramineae) II: an update and a comparison of two 2015 classifications. J. Syst. Evol. 55(4): 259–290. https://doi.org/10.1111/jse.12262

Stebbins, G.L. & Love, R.M. 1941. A cytological study of California forage grasses. Amer. J. Bot. 28(5): 371−382. https://doi.org/10.1002/j.1537-2197.1941.tb07983.x

Štorchová, H., Hrdličková, R., Chrtek, J., Tetera, M., Fitze, D. & Fehrer, J. 2000. An improved method of DNA isolation from plants collected in the field and conserved in saturated NaCl/CTAB solution. Taxon 49(1): 79–84. https://doi.org/10.2307/1223934

Syme, A.E. 2011. Diversification rates in the Australasian endemic grass *Austrostipa*: 15 million years of constant evolution. Pl. Syst. Evol. 298: 221–227. doi: 10.1007/s00606-011-0539-7

Syme, A.E., Murphy, D.J., Holmes, G.D., Gardner, S., Fowler, R. & Cantrill, D.J. 2012. An expanded phylogenetic analysis of *Austrostipa* (Poaceae: Stipeae) to test infrageneric relationships. *Austral*. Syst. Bot. 25: 1–10. doi:10.1071/SB10049

Tateoka, T. 1954. Karyotaxonomic studies in Poaceae, II. Rep. (Annual) Natl. Inst. Genet. 5: 68–69.

Tateoka, T. 1955. Karyotaxonomy in Poaceae III. Further studies of somatic chromosomes. Cytologia 20(4): 296–306. https://doi.org/10.1508/cytologia.20.296

Tateoka, T. 1956. Notes on some grasses I. Bot. Mag. (Tokyo) 69 (nos. 817–818): 311–315. https://doi.org/10.15281/jplantres1887.69.311

Terrell, E.E. & Wergin, W.P. 1981. Epidermal features and silica deposition in lemmas and awns of *Zizania* (Gramineae). Amer. J. Bot. 68(5): 697–707. https://doi.org/10.1002/j.1537-2197.1981.tb12402.x

Terrell, E.E., Peterson, P.M. & Wergin, W.P. 2001. Epidermal features and spikelet micromorphology in *Oryza* and related genera (Poaceae: Oryzeae). Smithsonian Contr. Bot. 91: 1–50. https://doi.org/10.5479/si.0081024X.91

Thomasson, J.R. 1978. Epidermal patterns of the lemma in some fossil and living grasses and their phylogenetic significance. Science 199: 975–977. doi: 10.1126/science.199.4332.975

Thomasson, J.R. 1981. Micromorphology of the lemma in *Stipa robusta* and *Stipa viridula* (Gramineae: Stipeae): taxonomic significances. S. W. Naturalist 26(2): 211–214. doi: 10.2307/3671126

Thomasson, J.R. 1986. Lemma epidermal features in the North American species of *Melica*, and selected species of *Briza*, *Catabrosa*, *Glyceria*, *Neostapfia*, *Pleuropogon* and *Schizachne* (Gramineae). Syst. Bot. 11: 253–262.

Tkach, N., Röser, M., Suchan, T., Cieślak, E., Schönswetter, P. & Ronikier, M. 2019. Contrasting evolutionary origins of two mountain endemics: *Saxifraga wahlenbergii* (Western Carpathians) and *S. styriaca* (Eastern Alps). B. M. C. Evol. Biol. 19: 18. https://doi.org/10.1186/s12862-019-1355-x

Tkach, N., Schneider, J., Doering, E., Wölk, A., Hochbach, A., Nissen, J., Winterfeld, G., Meyer, S., Gabriel, J., Hoffmann, M.H. & Röser, M. 2020. Phylogenetic lineages and the role of hybridization as driving force of evolution in grass supertribe Poodae. Taxon 69 (in press). https://doi.org/10.1002/tax.12204

Tzvelev, N.N. 1976. *Zlaki SSSR*. [*Grasses of the Soviet Union*.] Leningrad: Nauka.

Tzvelev, N.N. 1977. O proiskhozhdenii i evolutsii kovyley (*Stipa* L.). [On the origin and evolution of feathergrasses (*Stipa* L.).] Pp. 139–150 in: Lebedev, D.V. & Karamysheva, Z.V. (eds.), Problemy ekologii, geobotaniki, botanicheskoi geografii i floristiki. Leningrad: Academiya Nauk SSSR.

Tzvelev, N.N. 2011. Zametki o tribe kovylevykh (Stipeae Dumort., Poaceae). [Notes on the tribe Stipeae Dumort. (Poaceae).] Novosti Sist. Vyssh. Rast. 43: 20–29.

Valdés-Reyna, J. & Hatch, S.L. 1991. Lemma micromorphology in the Eragrostideae (Poaceae). Sida 14(4): 531–549. https://www.jstor.org/stable/41961070

Vázquez, F.M & Devesa, J.A. 1996. Revisión del género *Stipa* L. y *Nassella* Desv. (Poaceae) en la Península Ibérica e Islas Baleares. Acta Bot. Malac. 21: 125–189

Vázquez Pardo, F.M. & Gutiérrez Esteban, M. 2011. Classification of species of *Stipa* with awns having plumose distal segments. Telopea 13: 155–176.

Vickery, J.W., Jacobs, S.W.L. & Everett, J. 1986. Taxonomic studies in *Stipa* (Poaceae) in Australia. Telopea 3: 1–132.

Williams, A.R. 2011. *Austrostipa* (Poaceae) subgenus *Lobatae* in Western Australia. Telopea 13: 177–192.

Winterfeld, G. 2006. Molekular-cytogenetische Untersuchungen an Hafergräsern (Aveneae) und anderen Poaceae. Stapfia 86: 1–170.

Winterfeld, G., Schneider, J., Becher, H., Dickie, J. & Röser, M. 2015. Karyosystematics of the Australasian stipoid grass *Austrostipa* and related genera: chromosome sizes, ploidy, chromosome base numbers, and phylogeny. *Austral*. Syst. Bot. 28: 145–159. http://doi.org/10.1071/SB14029

Wölk, A. & Röser, M. 2014. Polyploid evolution, intercontinental biogeographical relationships and morphology of the recently described African oat genus *Trisetopsis* (Poaceae). Taxon 63: 773–788. https://doi.org/10.12705/634.1

Wölk, A. & Röser, M. 2017. Hybridization and long-distance colonization in oat-like grasses of South and East Asia, including an amended circumscription of *Helictotrichon* and the description of the new genus *Tzveleviochloa* (Poaceae). Taxon 66: 20–43. https://doi.org/10.12705/661.2

Wu, Z.-Y. & Phillips, S.M. 2006. Stipeae. Pp. 188–212 in: Wu, Z.-Y., Raven, P.H. & Hong, D.Y. (eds.), Flora of China, vol. 22. Beijing: Science Press, St. Louis: Missouri Botanical Garden Press.

Zhang, Z.-S., Li, L.-L. & Chen, W.-L. 2018. Chromosome number and karyotype of Phaenospermateae and Duthieeae (Poaceae), with reference to their systematic implications. Nordic J. Bot. 36: e01918. doi:10.1111/njb.01918

## REFERENCES

Agapova, N.D., Arkharova, K.B., Vakhtina, E.A., Zemskova, E.A. & Tarvis, L.V. 1993. Chisla khromosom tsvetkovykh rasteniy flory SSSR: Moraceae–Zygophyllaceae [Chromosome numbers in flowering plants of the flora of the USSR: Moraceae–Zygophyllaceae]. St. Petersburg: Nauka, 430 pp.

Andreev, V.N. 1982. Pp. 575–576 in: Löve, Á. (ed.), IOPB chromosome number reports LXXVI. Taxon 31(3): 574–598. https://doi.org/10.1002/j.1996-8175.1982.tb03560.x

Avdulov, N.P. 1928. Systematicheskaya karyologiya semeystva Gramineae. Dnyevnik vsesoyuznogo s’ezda botanikov v Leningrade (1928 g.), vol. 1: 65–67. Not seen, cited after Avdulov (1931).

Avdulov N.P. 1931. Kario-sistematicheskoe issledovanie semeistva zlakov. [Karyosystematic study of the family of grasses.] *Trudy Prikl. Bot. Selekts.*, Suppl. 44: 1–428.

Baden, C. 1983. Chromosome numbers in some Greek angiosperms. Willdenowia 13(2): 335–336. https://www.jstor.org/stable/3995849

Belyaeva, V.A. & Siplivinsky, V.N. 1977. Khromosomnye chisla i taksonomiya nekotorykh vidov baikalskoy flory. III. [Chromosome numbers and taxonomy of some species of Baikal flora. III.]. Bot. Zhurn. (Moscow & Leningrad) 62(8): 1132–1142.

Borgen, L. 1970. Chromosome numbers of Macaronesian flowering plants. Nytt Mag. Bot. 17: 145– 161.

Bowden, W.M. 1960. Chromosome numbers and taxonomic notes on northern grasses: III. Twenty-five genera. Canad. J. Bot. 38(4): 541–557. https://doi.org/10.1139/b60-049

Bowden, W.M. & Senn, H.A. 1962. Chromosome numbers in 28 grass genera from South America. Canad. J. Bot. 40(8): 1115–1124. https://doi.org/10.1139/b62-102

Brown, W.V. 1949. A cytological study of cleistogamous *Stipa leucotricha*. Madroño 10(4): 97–107. https://www.jstor.org/stable/41422661

Brown, W.V. 1951. Chromosome numbers of some Texas grasses. Bull. Torrey Bot. Club. 78(4): 292–299. doi: 10.2307/2481991

CCDB (The Chromosome Counts Database). 2020. Version 1.46. http://ccdb.tau.ac.il/home/. Accessed 21 Apr 2020.

Chepinoga, V.A. & Gnutikov, A.A. 2014. P. 1387 in: Marhold, K. (ed.), IAPT/IOPB chromosome data 18. Taxon 63(6): 1387–1393. https://doi.org/10.12705/636.37

Chepinoga, V.V., Gnutikov, A.A., Lubogoschinsky, P.I., Isaikina, M.M. & Konovalov, A.S. 2012. Pp. 981–982 in: Marhold, K. (ed.), IAPT/IOPB chromosome data 13. Taxon 61(4): 889–902. https://doi.org/10.1002/tax.614023

Chopanov, P. & Yurtsev, V.N. 1976. Khoromosomnye chisla nekotorykh zlakov Turkmenii. II. [Chromosome numbers of some grasses of Turkmenia. II.] Bot. Zhurn. (Moscow & Leningrad) 61(9): 1240–1244.

Covas, G. 1945. Número de cromosomas de algunas Gramíneas argentinas. Revista Argent. Agron. 12: 315–317.

Covas, G. & Bocklet, M. 1945. Número de cromosomas de algunas Gramineae-Stipinae de la flora Argentina. Revista Argent. Agron. 12: 261−265.

Dahlgren, P., Karlsson, T. & Lassen, P. 1971. Studies on the flora of the Balearic Islands I. Chromosome numbers in Balearic angiosperms. Bot. Not. 124: 249–269.

Darlington, C.D. & Wylie, A.P. 1956. Chromosome atlas of flowering plants. New York: Macmillan.

Davidse, G. & Pohl, R.W. 1974. Chromosome numbers, meiotic behavior, and notes on tropical American grasses (Gramineae). Canad. J. Bot. 52(2): 317–328. https://doi.org/10.1139/b74-042

Davidse, G. & Pohl, R.W. 1978. Chromosome numbers of tropical American grasses (Gramineae) 5. Ann. Missouri Bot. Gard. 65(2): 637–649. doi: 10.2307/2398863

Dawson, M.I. & Beuzenberg, E.J. 2000. Contributions to a chromosome atlas of the New Zealand flora – 36. Miscellaneous families, New Zealand J. Bot. 38(1): 1–23. https://doi.org/10.1080/0028825X.2000.9512671

Dedecca, D.M. 1954. Studies on the California species of *Stipa* (Gramineae). Madroño 12(5): 129–139. https://www.jstor.org/stable/41422801

Delay, C. 1947. Recherches sur la structure des noyaux quiescents chez les Phanérogames. Rev. Cytol. Cytophysiol. Vég. 9(1–4): 169–222; 10(1–4): 103–229.

Delay, C. 1950. Nombres chromosomiques chez les Phanérogames (I-re liste: 1938 à 1950). Rev. Cytol. Cytophysiol. Vég. 12(1–2): 1–368. Not seen, cited after Gould (1966) and Fedorov (1969).

Devesa, J.A., Ruiz, T., Viera, M.C., Tormo, R., Vázquez, F., Carrasco, J.P., Ortega, A. & Pastor, J. 1991. Contribución al conocimiento cariológico de las Poaceae en Extremadura (España) III. Bol. Soc. Brot. Série 2, 64: 35–74.

de Winter, B. 1965. The South African Stipeae and Aristideae (Gramineae). An anatomical, cytological and taxonomic study. Bothalia 8(3):199–404.

Duckert-Henriod, M.M. 1991. Pp. 229–236 in: Kamari, G., Felber, F. & Garbari, F. (eds.), Mediterranean chromosome number reports 1. Fl. Medit 1: 223–245. http://www.herbmedit.org/flora/01-223.pdf

Edgar, E. & Connor, H.E. 2000. Flora of New Zealand, vol. 5, Gramineae. Lincoln, New Zealand: Manaaki Whenua Press.

Erst, A.S., Mitrenina, E.Yu., Sukhorukov, A.P., Kuzmin, I.V., Gudkova, P.D., Tashev, A.N., Xiang, K. & Wang, W. 2019. P. 1126 in: Marhold, K. & Kučera, J. (eds.), IAPT chromosome data 30. Taxon 68: 1124–1130. https://doi.org/10.1002/tax.12156

Faruqi, S.A., Quraish, H.B. & Inamuddin, M. 1987. Studies in Libyan grasses. X. Chromosome number and some interesting features (1). Ann. Bot. (Rome) 45(2): 75–102.

Fedorov, A.A. (ed.). 1969. Khromosomnye chisla tsvetkovykh rasteniiy. [Chromosome numbers of flowering plants]. Leningrad: Nauka, 927 pp.

Ferchichi, A., Nabli, N.A. & Delay, J. 1994. Prospection caryologique de la famille des Poaceae en Tunisie steppique. Acta Bot. Gallica. 141(3): 327–341. https://doi.org/10.1080/12538078.1994.1051516

Fernandes, A. & Queirós, M. 1969. Contribution à la connaissance cytotaxinomique des Spermatophyta du Portugal. I. Gramineae. Bol. Soc. Brot. Série 2, 43: 20–144.

Gallego, M.J. & Talavera, S. 1994. Números cromosomáticos de plantas occidentales, 696–707. Anales Jard. Bot. Madrid. 51(2): 280.

Gervais, C. 1966. Nombres chromosomiques chez quelques graminées alpines. Bull. Soc. Neuchâtel. Sci. Nat. 89: 87–100. http://doi.org/10.5169/seals-88960

Ghukasyan, A.G. 2004. Kariologisheskaya izuchennost zlakov (Poaceae) Armenii. [Extent of karyological study of Armenian grasses (Poaceae).] Fl. Rastitel’nost’ Rastitel’nye Resursy Armenii [Flora, vegetation and plant resources of Armenia] 15: 74–84. http://takhtajania.asj-oa.am/id/eprint/297

Gnutikov, A.A., Myakoshina, Yu.A., Punina, E.O. & Rodionov, A.V. 2017. A karyological study of grasses (Poaceae) of Altai. II. Turczaninowia 20(2): 16–22. https://doi.org/10.14258/turczaninowia.20.2.2

Gohil, R.N. & Koul, K.K. 1986. SOCGI plant chromosome number reports IV, Gramineae (Poaceae). J.Cytol. Genet. 21: 155.

Gould, F.W. 1958. Chromosome numbers in southwestern grasses. Amer. J. Bot. 45(10): 757–767. https://doi.org/10.1002/j.1537-2197.1958.tb10608.x

Gould, F.W. 1965. Chromosome numbers in some Mexican grasses. Bol. Soc. Bot. México. 29: 49– 62. https://doi.org/10.17129/botsci.1088

Gould, F. W. 1966. Chromosome numbers of some Mexican grasses. Canad. J. Bot. 44(12): 1683– 1696. https://doi.org/10.1139/b66-181

Gould, F.W. 1968. Chromosome numbers of Texas grasses. Canad. J. Bot. 46(10): 1315–1325. https://doi.org/10.1139/b68-175

Gould, F.W. & Soderstrom, T.R. 1970. Pp. 104–104 in: Löve, Á. (ed.), IOPB chromosome number reports. XXV. Taxon 19: 102–113. https://doi.org/10.1002/j.1996-8175.1970.tb02985.x

Gusik, M.B. 1973. Sutochnyi ritm i ekologiya tsveteniya i opyleniya zlakov stepei Zabaikalya i Khakassii [Daily rhythm and ecology of flowering and pollination in the grasses of the steppes of Transbaikal and Khakassia regions). Ph.D. thesis abstract, Perm. Not seen, cited after Tzvelev (1976).

Gusik, M.B. & Levkovskiy, V.P. 1979. Khromosomnye chisla dikorastushchikh zlakov Zabaikalya i Khakassii. [Chromosome numbers of wild grasses of Transbaikal and Khakassia regions]. Ekologia opylenia. Perm. 4: 26–32. Not seen, cited after Agapova & al. (1993).

Gupta, R.C., Chauhan, H.S., Saggoo, M.I.S. & Kaur, N. 2014. Cytomorphology of some grasses (Poaceae) from Lahaul-Spiti (Himachal Pradesh), India. Biolife 2(4): 1234–1247.

Hatch, S.L. 1980. Chromosome numbers of some grasses from the southwestern United States and Mexico. The Southwestern Naturalist 25(2): 278–280. doi: 10.2307/3671259

Hatch, S.L. & Bearden, D.A. 1983. *Stipa curvifolia* (Poaceae) – studies on a rare taxon. Sida 10(2): 184–187. https://www.jstor.org/stable/23909549

Hitchcock, C.L. & Spellenberg, R.W. 1968. A new *Oryzopsis* from Idaho. Brittonia 20: 162–165. https://doi.org/10.2307/2805618

Johnson, B.L. 1945. Cytotaxonomic studies in *Oryzopsis*. Bot. Gaz. 107(1): 1–32. https://www.jstor.org/stable/2472124

Johnson, B.L. 1960. Natural hybrids between *Oryzopsis* and *Stipa*. I. Oryzopsis hymenoides × Stipa speciosa. Amer. J. Bot. 47: 736–742. https://doi.org/10.1002/j.1537-2197.1960.tb07159.x

Johnson, B.L. 1962a. Natural hybrids between *Oryzopsis* and *Stipa*. II. Oryzopsis hymenoides × Stipa nevadensis. Amer. J. Bot. 49: 540–546. https://doi.org/10.1002/j.1537-2197.1962.tb14978.x

Johnson, B.L. 1962b. Amphiploidy and introgression in *Stipa*. Amer. J. Bot. 49: 253–262. https://doi.org/10.1002/j.1537-2197.1962.tb14935.x

Johnson, B.L. 1963. Natural hybrids between *Oryzopsis* and *Stipa*. III. Oryzopsis hymenoides × Stipa pinetorum. Amer. J. Bot. 50: 228–234. https://doi.org/10.1002/j.1537-2197.1963.tb12228.x

Johnson, B.L. & Rogler, G.A. 1943. A cyto-taxonomic study of an intergeneric hybrid between *Oryzopsis hymenoides* and *Stipa viridula*. Amer. J. Bot. 30(1): 49–56. https://doi.org/10.1002/j.1537-2197.1943.tb14731.x

Kaur, H., Gupta, R.C. & Kumari, S. 2011a. P. 1789 in: Marhold, K. (ed.), IAPT/IOPB chromosome data 12. Taxon 60 (6): 1784–1796. https://doi.org/10.1002/tax.606033

Kaur, H., Kumari, S., & Gupta, R.C. (2011b). New chromosome numbers reported in grasses from Himachal Pradesh (India). Chromosome Bot. 6(1): 13–16. doi:10.3199/iscb.6.13

Kerguélen, M. 1975. Les Gramineae (Poaceae) de la flore française. Essai de mise au point taxonomique et nomenclaturale. Lejeunia, nouv. sér. 75: 1–343.

Korobkov, A.A., Kotseruba, V.V., Probatova, N.S., Shatokhina, A.V. & Rudyka, E.G. 2013. pp. 1075–1077 in: Marhold, K. (ed.), IAPT/IOPB chromosome data 15. Taxon 62 (5): 1073–1083. https://doi.org/10.12705/625.16

Kožuharov, S. & Kuzmanov, B. 1968. Cytotaxonomic studies on Bulgarian Gramineae. C. R. Acad. Bulg. Sci. 21: 269–272.

Krogulevich, R.E. 1971. Rol poliploidii v genezise vysokogornoy flory Stanovogo nagorya. Pp. 115–214 in: Ekologiya flory Zabaikalya. Irkutsk. Not seen, cited after Probatova & Sokolovskaya (1980) and Murín & al. (1984).

Krogulevich, R.E. 1972. Vliyanie botaniko-geograficheskikh faktorov na poliploidiyu [Effect of botanical-geographic factors on polyploidy]. Pp: 12–13. in: Nauchnye chteniya pamyati M.G. Popova. lrkutsk. Not seen, cited after Tzvelev (1976).

Krasnikov, A.A. 1991. Chisla khromosom nekotorykh vidov sosudistykh rasteniy Novosibirskoy oblasti. [Chromosome numbers in some species of vascular plants from Novosibirsk region.] Bot. Zhurn. (Moscow & Leningrad) 76(3): 476–479.

Kumar, R., Kumari, V., & Singhal, V.K. 2018. Pp. 1238–1239 in: Marhold, K., Kučera, J. (eds.), IAPT chromosome data 28. Taxon 67 (6): 1235–1245. https://doi.org/10.12705/676.39

Kumar, V. & Subramaniam, B. 1987. Chromosome atlas of flowering plants of the Indian subcontinent, vol. 2, Monocotyledons and bibliography. Calcutta: Botanical Survey of India. Pp. 465–1095.

Kumari, V., Kumar, R. & Singhal, V.K. 2019. P. 1379 in: Marhold, K., Kučera, J. (eds.), IAPT chromosome data 31. Taxon 68 (6): 1374–1380. https://doi.org/10.1002/tax.12176

Kuzmanov, B. 1993. Chromosome numbers of Bulgarian angiosperms: an introduction to a chromosome atlas of the Bulgarian flora. Fl. Medit. 3: 19–163.

Labadie, J.P. 1979a. Pp. 628–629 in: Löve, Á. (ed.), IOPB chromosome number reports. LXV. Taxon 28: 627–637.

Labadie, J. 1979b. Étude caryosystematique de quelques espèces de la flore d’Algérie. Naturalia monspeliensia, sér. Bot. 32: 1–11.

de Litardière, R. 1950. Nombres chromosomiques de diverses graminées. Bol. Soc. Brot. sér. 2, 24: 79–87.

Lorenzo-Andreu, A. 1951. Cromosomas de plantas de la estepa de Aragón. III. An. Estac. Exp. Aula Dei 2(2): 195–203. http://hdl.handle.net/10261/38176

Lorenzo-Andreu, A. 1953. Notas sobre cariología. Pp. 87–91 in: Servicio del Esparto (ed.), Estudios y experiencias sobre el esparto, 2.^a^ parte. Madrid: Ministerios de Industria, Comercio y Agricultura. Not seen, cited after Vazques & Devesa (1991)

Löve, Á. & Kjellqvist, E. 1973. Cytotaxonomy of Spanish plants. II. Moncotyledons. Lagascalia 3(2): 147–182.

Löve, Á. & Löve, D. 1961. Chromosome numbers of central and northwest European plant species. Opera Bot. 5: 1–581. Not seen, cited after Fedorov (1969).

Löve, Á. & Löve, D. 1974. Cytotaxonomical atlas of the Slovenian flora. Lehre: Cramer.

Löve, Á. & Löve, D. 1981. Pp. 509–511 in: Löve, Á. (ed.), Chromosome number reports. LXXI. Taxon 30(2): 507–516. https://doi.org/10.1002/j.1996-8175.1981.tb00780.x

Love, R.M. 1946. Interspecific hybridization in *Stipa* L. I. Natural hybrids. Amer. Naturalist. 80 (no. 788): 189–192. https://doi.org/10.1086/281376

Love, R.M. 1954. Interspecific hybridization in *Stipa* II. Hybrids of *S. cernua*, *S. lepida*, and *S. pulchra*. Amer. J. Bot. 41: 107–110. https://doi.org/10.1002/j.1537-2197.1954.tb14311.x

Lungeanu, I. 1975. Pp. 503–504 in: Löve, Á. (ed.), IOPB chromosome number reports XLIX. Taxon 24(4): 501–516. https://doi.org/10.1002/j.1996-8175.1975.tb00341.x

Lungeanu, I. 1980. Pp. 164–165 in: Löve, Á. (ed.), IOPB chromosome number reports. LXVI. Taxon 29(1): 163–169. https://doi.org/10.1002/j.1996-8175.1980.tb00607.x

Luque, T. & Diaz Lifante, Z. 1991. Chromosome numbers of plants collected during Iter Mediterraneum I in the SE of Spain. Bocconea 1: 303–364. http://www.herbmedit.org/bocconea/1-303.pdf

Magulaev, A.Yu. 1984. Tzitotaksonomicheskoe izuchenie nekotorykh tzvetkovykh rasteniy Severnogo Kavkasa. [Cytotaxonomic study in some flowering plants of the North Caucasus.] Bot. Zhurn. (Moscow & Leningrad) 69(4): 511–517.

Májovskýp, J. (ed.). 1974. Index of chromosome numbers of Slovakian flora. (Part 3). Acta Fac. Rerum Nat. Univ. Comen., Bot. 22: 1–20.

Májovský, J. (ed.). 1978. Index of chromosome numbers of Slovakian flora (Part 6). Acta Fac. Rerum Nat. Univ. Comen., Bot. 26: 1–42.

Marhold, K., Mártonfi, P., Mereďa, P. & Mráz, P. (eds.). 2007. Chromosome number survey of the ferns and flowering plants of Slovakia. Bratislava: Veda.

Mehra, P.N. & Sharma, M.L. 1975a. Cytological studies in some central and eastern Himalayan grasses. IV. The Arundinelleae, Eragrosteae, Isachneae, Chlorideae, Sporoboleae, Meliceae, Stipeae, Arundineae and Garnotieae. Cytologia 40: 453–462. https://doi.org/10.1508/cytologia.40.453

Mehra, P.N. & Sharma, M.L. 1975b. Pp. 501–502 in: Löve, Á. (ed.), IOPB chromosome number reports XLIX. Taxon 24(4): 501–516. https://doi.org/10.1002/j.1996-8175.1975.tb00341.x

Mehra, P.N. & Sunder, S. 1969. Cytological studies in the north Indian grasses. Part II. Res. Bull. Panjab Univ. Sci. 20: 503–539.

Měsíček, J. & Javůrcová-Jarolímová, V. 1992. List of chromosome numbers of the Czech vascular plants. Praha: Academia. 144 pp.

Měsíček, J. & Soják, J. 1972. Chromosome studies in Mongolian plants. Preslia 44: 334–358.

Moinuddin, M., Vahidy, A.A. & Ali, S.I. 1994. Chromosome counts in Arundinoideae, Chloridoideae, and Pooideae (Poaceae) from Pakistan. Ann. Missouri Bot. Gard. 81(4): 784– 791. doi: 10.2307/2399923

Moraldo, B. 1986. Il genere *Stipa* L. (Gramineae) in Italia. Webbia 40(2): 203–278. https://doi.org/10.1080/00837792.1986.10670388

Murín, A., Háberová, I. & Žamsran, C. 1984. Further karyological studies of the Mongolian flora. Folia Geobot. Phytotax.19: 28–39. https://www.jstor.org/stable/4180474

Murray, B.G., de Lange, P.J. & Ferguson, A.R. 2005. Nuclear DNA variation, chromosome numbers and polyploidy in the endemic and indigenous grass flora of New Zealand. Ann. Bot. (Oxford) 96: 1293–1305. https://doi.org/10.1093/aob/mci281

Nielsen, E.L. 1939. Grass studies III. Additional somatic chromosome complements. Amer. J. Bot. 26(6): 369–372. https://doi.org/10.1002/j.1537-2197.1939.tb09289.x

Ono, H. & Tateoka, T. 1953. Karyotaxonomy in Poaceae I. Chromosomes and taxonomic relations in some Japanese grasses. Bot. Mag. (Tokyo) 66: 18–27.

Parodi, L.R. 1946. Gramineas bonaerenses. Clave para la determinación de los géneros y enumeración de las especies, 4th ed. Buenos Aires: Centro Estudiantes de Agronomía. Not seen, cited after Fedorov (1969) and Romaschenko & al. (2012).

Petrova, O.A. 1968. Khromosomnyi sostav nekotorykh zlakov flory Ukrainy v svyasi s usloviami ikh proizrastaniya. [Chromosomal composition of some Ukrainian grasses according to their growing conditions.] Pp. 37–39 in: Nikitin, V.N., (ed.), Biologicheskaya nauka v universitetakh I pedagogicheskikh institutakh Ukrainy za 50 let. [Biological science in universities and pedagogical institutes of Ukraine for 50 years]. Charkov: Charkov University. Not seen, cited after Prokudin & al. (1977) and Agapova & al. (1993).

Podlech, D. & Dieterle, A. 1969. Chromosomenstudien an afghanischen Pflanzen. Candollea 24: 185–243.

Pohl, R.W. 1954.The allopolyploid *Stipa latiglumis*. Madroño 12(5): 145–150. https://www.jstor.org/stable/41422803

Pohl, R.W. & Davidse, G. 1971. Chromosome numbers of Costa Rican grasses. Brittonia 23(3): 293– 324. doi: 10.2307/2805632

Probatova, N.S. & Seledets, V. 2008. Pp. 553–558 in: Marhold, K. (ed.), IAPT/IOPB chromosome data 5. Taxon 57(2): 553–562. https://doi.org/10.2307/25066021

Probatova, N.S. & Sokolovskaya, A.P. 1980. K kariotaksonomicheskomu isucheniyu zlakov gornogo Ataya. [To the karyotaxonomic study of the grasses of Mountain Altai.] Bot. Zhurn. (Moscow & Leningrad) 65(4): 509–520.

Probatova, N.S., Rudyka, E.G., Pavlova, N.S., Verkholat, V.P. & Nechaev, V.A. 2006. Chisla khromosom vidov rasteniy iz Primorskogo kraya, Priamurya i Magadanskoy oblasti. [Chromosome numbers of plants of the Primorsky Territory, the Amur River basin and Magadan region.] Bot. Zhurn. (Moscow & Leningrad) 91(3): 491–509.

Probatova, N.S., Rudyka, E.G., Seledets, V.P. & Nechaev, V.A. 2008. Pp. 1268–1271 in: Marhold, J. (ed.), 2008. IAPT/IOPB chromosome data 6. Taxon 57(4): 1267–1273. https://doi.org/10.1002/tax.574017

Probatova, N.S., Sedelets, V.P., Rudyka, E.G., Gnutikov, A.A., Kozhevnikova, Z.V., Barkalov & V.Yu. 2009. Pp.1284–1288 in: Marhold, K. (ed.), IAPT/IOPB chromosome data 8. Taxon 58(4): 1281–1289. https://doi.org/10.1002/tax.584017

Probatova, N.S., Kazanovsky, S.G., Rudyka, E.G., Barkalov, V.Yu., Seledets, V.P. & Nechaev, V.A. 2011. Pp. 1790–1794 in: Marhold, K. (ed.), IAPT/IOPB chromosome data 12. Taxon 60 (6): 1784–1796. https://doi.org/10.1002/tax.606033

Probatova, N.S., Kazanovsky, S.G., Shatokhina, A.V. Rudyka, E.G., Verkhozina, A.V. & Krivenko, D.A. 2012. Pp. 1342–1344 in: Marhold, K. (ed.), IAPT/IOPB chromosome data 14. Taxon 61(6): 1336–1345. https://doi.org/10.1002/tax.616027

Probatova, N.S., Kazanovsky, S.G., Rudyka, E.G., Gnutikov, A.A. & Verkhozina, A.V. 2013. Pp. 1080–1081 in: Marhold, K. (ed.), IAPT/IOPB chromosome data 15. Taxon 62(5): 1073–1083. https://doi.org/10.12705/625.16

Probatova, N.S., Kazanovsky, S.G., Barkalov, V.Yu, Rudyka, E.G. & Shatokhina, A.V. 2015. Pp. 1348–1349 in: Marhold, K. (ed.), IAPT/IOPB chromosome data 20. Taxon 64(6): 1344–1350. https://doi.org/10.12705/646.42

Probatova, N.S., Kazanovsky, S.G., Barkalov, V.Yu. & Nechaev, V.A. 2016a. Pp. 1202–1204 in: Marhold, K., Kučera, J. (eds.), IAPT/IOPB chromosome data 22. Taxon Vol. 65(5): 1200–1207. https://doi.org/10.12705/655.40

Probatova, N.S., Krivenko, D.A. & Ebel, A.L. 2016b. Pp. 1204–1205 in: Marhold, K., Kučera, J. (eds.), IAPT/IOPB chromosome data 22. Taxon Vol. 65(5): 1200–1207. https://doi.org/10.12705/655.40

Prokudin, Yu.N, Vovk, A.G, Petrova, O.A, Ermolenko, E.D. & Vernichenko, Yu.V. (ed.). 1977. Zlaki Ukrainy [Grasses of Ukraine]. Kiev: Naukova Dumka.

Queirós, M. 1973. Contribuição para o conhecimento citotaxonómico das Spermatophyta de Portugal. I. Gramineae, supl. 1. Bol. Soc. Brot. Ser. 2, 47: 77–103.

Queirós, M. 1974. Contribuição para o conhecimento citotaxonómico das Spermatophyta de Portugal. I. Gramineae, supl. 2. Bol. Soc. Brot. Ser. 2, 48: 81–98.

Reeder, J.R. 1967. Notes on Mexican grasses VI. Miscellaneous chromosome numbers. Bull. Torrey Bot. Club 94(1): 1–17. https://www.jstor.org/stable/2483595

Reeder, J.R. 1968. Notes on Mexican Grasses VIII. Miscellaneous chromosome numbers – 2. *Bull. Torrey Bot*. Club 95(1): 69–86. https://www.jstor.org/stable/2483808

Reeder, J.R. 1977. Chromosome numbers in western grasses. Amer. J. Bot 64(1): 102–110. https://doi.org/10.1002/j.1537-2197.1977.tb07611.x

Reeder, J.R. 1984. Pp. 132–133 in: Löve, Á. (ed.), Chromosome number reports LXXXII. Taxon 33(1): 126–134. https://doi.org/10.1002/j.1996-8175.1984.tb02474.x

Reeder, J.R. & Reeder, C.G. 1968. *Parodiella*, a new genus of grasses from the high Andes. Bol. Soc. Argent. Bot. 12: 268–283.

Reese, G. 1957. Über die Polyploidiespektren in der nordsaharischen Wüstenflora. Flora 144(4): 598– 634.

Rudyka, E.G. 1990. Chisla khromosom sosudistykh rasteniy iz raslichnykh regionov SSSR. [Chromosome numbers of vascular plants from the various regions of the USSR.] Bot. Zhurn. (Moscow & Leningrad) 75(12): 1783–1786.

Saura, F. 1943. Cariología de Gramíneas. Géneros Paspalum, Stipa, Poa, Andropogon y Phalaris. Rev. Fac. Agron. Vet. (Buenos Aires) 10: 344–353. Not seen, cited after Myers (1947) and Bowden & Senn (1962).

Saura, F. 1948. Cariología de Gramíneas en Argentina. Rev. Fac. Agron. Vet. (Buenos Aires) 12: 56–67. Not seen, cited after Bowden & Senn (1962).

Scholz, H., Oberprieler, C. & Vogt, R. 1998. Chromosome numbers of North African phanerogams. VII. Some notes on North African Gramineae. Lagascalia 20: 265–275.

Sharma, M.L. & Sharma, K. 1978. Pp. 390–391 in: Löve, Á. (ed.), IOPB chromosome number reports LXI. Taxon 27(4): 375–392. https://doi.org/10.1002/j.1996-8175.1978.tb04261.x

Sharma, M.L. & Sharma, K. 1979. Cytological studies in the north Indian grasses. Cytologia 44: 861–872.

Shechter, Y. & Johnson, B.L. 1966. A new species of *Oryzopsis* (Gramineae) from Wyoming. Brittonia 18(4): 342–347. doi: 10.2307/2805150

Sheidai, M. & Attaei, S. 2005. Meiotic studies of some *Stipa* (Poaceae) species and populations in Iran. Cytologia 70(1): 23–31.

Singhal, V.K., Kumari, V. & Kumar, P.V. 2014. Cytomorphological diversity in some selected members of Poaceae from Parvati Valley in Kullu district of Himachal Pradesh, India. Pl. Syst. Evol. 300(6): 1385–1408. doi: 10.1007/s00606-013-0969-5

Skalińska, M., Pogan, E. & Jankun, A. 1968. Dalsze badania nad kariologią flory polskiej Cz. VII. [Further studies in chromosome numbers of Polish angiosperms. Seventh contribution.] *Acta Biol. Cracov.*, Ser. Bot. 11(2): 199–224.

Sokolovskaya, A.P. & Probatova, N.S. 1977. Kariologicheskoe issledovanie zlakov (Poaceae) yuzhnoy chasti Sovetskogo Dalnego Vostoka. [Karyological investigation of grasses (Poaceae) in southern part of the Soviet Far East.] Bot. Zhurn. (Moscow & Leningrad) 62(8): 1143–1153.

Sokolovskaya, A.P. & Probatova, N.S. 1978. Khromosomnye chisla nekotorykh zlakov (Poaceae) flory SSSR. II. [Chromosome numbers of some grasses (Poaceae) of the U.S.S.R. flora. II.] Bot. Zhurn. (Moscow & Leningrad) 63(9): 1247–1257.

Sokolovskaya, A.P. & Strelkova, O.S. 1939. Geograficheskoye raspredelenie poliploidov. I. Issledovanie rastitelnosti Pamira. Uch. zap. Leningr. gor. univ., ser. biol. 9: 35; Tr. Petergofsk. boil. inst. 17: 42–63. Not seen, cited after Fedorov (1969).

Sokolovskaya, A.P. & Strelkova, O.S. 1948. Geograficheskoye raspredelenie poliploidov. II. Issledovanie flory Altaya. *Uch. zap. Ped. Inst. im. Gertzena*, Leningrad 66: 179–193. Not seen, cited after Fedorov (1969).

Solntzeva, M.P. 1967. Mikrosporogenez i razvitie muzhskogo gametofita u kovyley. [The microsporogenesis and the development of the male gamethophyte in feather-grasses.] Bot. Zhurn. (Moscow & Leningrad) 52(12): 1757–1772.

Spellenberg, R. 1968. Notes on *Oryzopsis hendersonii* (Gramineae). Madroño 19(7) 283–286. https://www.jstor.org/stable/41423321

Spellenberg, R. 1970. Pp. 112–113 in: Löve, A. (ed.), IOPB chromosome number reports. XXV. Taxon 19(1): 102–113. https://doi.org/10.1002/j.1996-8175.1970.tb02985.x

Stepanov, N.V. 1994. Chisla khromosom nekotorykh taksonov vysshikh rasteniy flory Krasnoyarskogo kraya. [Chromosome numbers of some higher plants taxa of the flora of Krasnoyarsk region.] Bot. Zhurn. (Moscow & Leningrad) 79(2): 135–139.

Strid, A. 1983. Pp. 139–140 in: Löve, A. (ed.), IOPB chromosome number reports LXXVIII. Taxon 32(1): 138–141. https://doi.org/10.1002/j.1996-8175.1983.tb02410.x

Strid, A. 1987. Chromosome numbers of Turkish mountain plants. Notes Roy. Bot. Gard. Edinburgh 44(2): 351–356.

Strid, A. & Andersson, I.A. 1985. Chromosome numbers of Greek mountain plants. An annotated list of 115 species. Bot. Jahrb. Syst. 107(1–4): 203–228.

Strobl, W. & Wittmann, H. 1985. Beiträge zur Kenntnis von Verbreitung, Soziologie und Karyologie von Achnatherum calamagrostis (L.) P.B. im Bundesland Salzburg (Österreich). Ber. Bayer. Bot. Ges. 56: 95–102.

Tarnavschi, I.T. 1948. Die Chromosomenzahlen der Anthophyten-Flora von Rumänien mit einem Ausblick auf das Polyploidie-Problem. *Bot. Mus. Bot. Univ. Cluj* 28, suppl. 1–130. [Buletinul Grădinii Botanice şi al Muzeului Botanic dela Universitatea din Cluj]. Not seen, cited after Fedorov (1969) and Vázquez & Devesa (1996).

Tarnavschi, I.T. & Lungeanu, I. 1970a. Pp. 609–610 in: Löve, A. (ed.), IOPB chromosome number reports XXVIII. Taxon 19: 608–610. https://doi.org/10.1002/j.1996-8175.1970.tb03062.x

Tarnavschi, I.T. & Lungeanu, I. 1970b. Chromosomenzahlen von einigen in Rumänien wildwachsenden Anthophyten. Rev. Roum. Biol., Bot. 15(6): 381–383.

Tateoka, T. 1957. Notes on some grasses VI. Bot. Mag. (Tokyo) 70: 119–125. https://doi.org/10.15281/jplantres1887.70.119

Tateoka, T. 1962. A cytological study of some Mexican grasses. Bull. Torrey Bot. Club. 89(2): 77– 82. doi: 10.2307/2482528

Tateoka, T. 1967. P. 561–564 in: Löve, A. (ed.), IOPB chromosome number reports. XIV. Taxon 16(6): 552–571. https://doi.org/10.1002/j.1996-8175.1967.tb02139.x

Tateoka, T. 1986. Chromosome numbers in the tribe Stipeae (Poaceae) in Japan. *Bull. Natl. Sci. Mus. Tokyo*, B, 12: 151–154.

Titova, N.N. 1935. Poiski rastitelnoy drosofily. [Searches for plant fruit flies.]. Sov. botanika 2: 61–62. Not seen, cited after Fedorov (1969).

Tzvelev, N.N. 1976. Zlaki SSSR. Leningrad: Nauka.

Vázquez, F.M. & Devesa, J.A. 1996. Revisión del género *Stipa* L. y *Nassella* Desv. (Poaceae) en la Península Ibérica e Islas Baleares. Acta Bot. Malac. 21: 125–189.

Valencia, J. I. & Costas, M. 1968. Estudios citotaxonómicos sobre *Piptochaetium* (Gramineae). Bol. Soc. Argent. Bot. 12: 168–179.

Verlaque, R., Reynaud, C. & Aboucaya, A. 1997. Pp. 240–246 in: Kamari, G., Felber, F. & Garbari, F. (eds.), Mediterranean chromosome number reports 7. Fl. Medit 7: 197–275. http://www.herbmedit.org/flora/7-197.pdf

Winterfeld, G., Schneider, J., Becher, H., Dickie, J. & Röser, M. 2015. Karyosystematics of the Australasian stipoid grass *Austrostipa* and related genera: chromosome sizes, ploidy, chromosome base numbers, and phylogeny. *Austral*. Syst. Bot. 28: 145–159. https://doi.org/10.1071/SB14029

Zhang, Z.S., Li, L.L. & Chen, W.L. 2017. *Ptilagrostis contracta* (Stipeae, Poaceae), a new species endemic to Qinghai-Tibet Plateau. PloS ONE 12(1): e0166603. https://doi.org/10.1371/journal.pone.0166603

